# Tetradic Dynamics of Dyadic Sensorimotor Coordination: A Multiscale EEG Hyperscanning Study

**DOI:** 10.64898/2026.06.21.732435

**Authors:** Emmanuel Olarewaju, Lena Palaniyappan, Guillaume Dumas

## Abstract

How do leader and follower roles shape the brain mechanisms that support coordinated action between people? This question has therapeutic relevance for conditions such as schizophrenia and autism spectrum disorder, where the capacity for reciprocal social coordination is a defining vulnerability. Here, we propose a tetradic framework and examine sensorimotor coordination in 16 healthy adult pairs (15 included in EEG analyses) using dual-brain EEG hyperscanning and a 2×2 within-subject design crossing Role (Leader/Follower) and Condition (*Mirroring*/*Matching*). *Mirroring* required resonance with a partner’s movement; *Matching* required its controlled transformation. This contrast was designed to dissociate automatic from controlled coordination. Behaviourally, *Mirroring* produced a robust Follower reaction-time advantage: pooling an independent behavioural-only cohort (combined N=64) revealed a smaller Leader advantage, consistent with role-dependent attention–inhibition gating. At the neural level, tonic power and significant phasic condition clusters were greater during *Matching* than *Mirroring*, while role-dependent phasic clusters suggested an anticipatory sequence around Leader responses and post-response gating and rebound around Follower responses. Information-theoretic decomposition identified Leader-unique predictive information in superior temporal regions and a separate frontal-pole interaction, Follower-unique predictive information in insular and superior frontal regions, sparse Synergy in the right rostral anterior cingulate cortex, and Redundancy spanning multiple cortical regions. These findings characterize tetradic coordination within dyads as a multiscale, role-asymmetric architecture in which top-down predictive control and bottom-up adaptive regulation are functionally dissociable. The tetradic framework provides a scaffold for this dissociation, and the role-specific signatures it reveals may offer candidate biomarkers for clinical populations in whom interpersonal coordination is disrupted.

**Highlights:**

- Leader–follower roles structure dyadic coordination mechanisms
- Mirroring yields a robust Follower RT advantage and a smaller Leader advantage
- Matching-dominant neural power spans tonic and phasic condition analyses
- Phasic dynamics suggest Leader anticipation, then Follower inhibition and rebound
- Information decomposition reveals role-specific inter-brain predictive architecture

## 1. Introduction

Two individuals are seated face-to-face across a table. One raises their hand; the other follows moments later, doing the same. The movements appear trivial, yet they reveal a fundamental structured asymmetry: one initiates, the other responds. When the task requires the responder to produce the corresponding *Matching* action (the same anatomical body-side response: Left side ↔ Left side/Right side ↔ Right side) rather than the *Mirroring* action (the opposite anatomical body-side response: Left side ↔ Right side/Right side ↔ Left side), something changes: top-down control is exerted, prepotent responses are modulated, and what would otherwise unfold as automatic imitation becomes a controlled transformation (Brass et al., 2001; Heyes, 2011; Kelso, 1995; Olarewaju, 2021). This distinction between *Mirroring* and *Matching* is situated at the center of a broader question that has occupied social neuroscience for decades (Brass et al., 2001; Cacioppo & Decety, 2011; Rizzolatti & Craighero, 2004; Schilbach et al., 2013; Sebanz et al., 2006): how do leader-follower roles influence the neural and behavioural mechanisms by which two individuals coordinate their actions? The answer to this question will help us understand how interpersonal dynamics become fragile and disconnected at different times and for different reasons.

The ease with which one person’s movement becomes another’s response, depends on a system of attention and inhibitory control whose disruption is a core feature of schizophrenia and autism spectrum disorder (Bolis et al., 2023; Olarewaju et al., 2023). These conditions are characterized by a failure to calibrate behaviours in social contexts: to know when to *Mirror* and when to *Match*, when to follow and when to initiate. These interdependent cycles of anticipation and adaptation fluctuate over time, vary with role demands, and manifest heterogeneously across neural, behavioural, and interpersonal scales. Understanding the baseline architecture of this system is, therefore, a prerequisite for disentangling what may go wrong within it.

A critical challenge in this enterprise is the problem of scale. Coordination unfolds simultaneously across multiple timescales (Kelso, 1995). When neural activity is averaged across repeated trials, complementary task conditions (automatic vs. effortful) may engage overlapping cortical networks, reflecting the global demands of sustained social interaction. At the timescale of successive responses, those same networks may also exhibit role-dependent temporal dynamics as the momentary demands on attention and inhibitory control fluctuate within each trial.

Previous hyperscanning studies have tended to collapse across these scales, averaging inter-brain processes over extended epochs and thereby conflating sustained network activity with transient event-driven dynamics (Babiloni & Astolfi, 2014; Hasson et al., 2012; Lindenberger et al., 2009). A complementary challenge is the problem of decomposition: inter-brain coupling is not a unitary phenomenon but a mixture of synergistic (*our*), unique (*mine* and *yours*), and redundant (*everyone*’*s*) informational components whose role-specific contributions have not previously been disentangled (Chidichimo et al., 2026).

We address both challenges using a tetradic framework (Kelso, 1995; Olarewaju, 2021) to characterize leader–follower sensorimotor coordination at multiple scales utilizing multimodal data. This framework formalizes face-to-face interaction as a four-quadrant spatiotemporal field defined by two axes: the left–right hemispheric axis, which divides each participant’s brain and body into halves, and the self–other interpersonal axis, which divides the dyad into distinct, but coupled, agents (see Figure 1).

**Figure 1.**
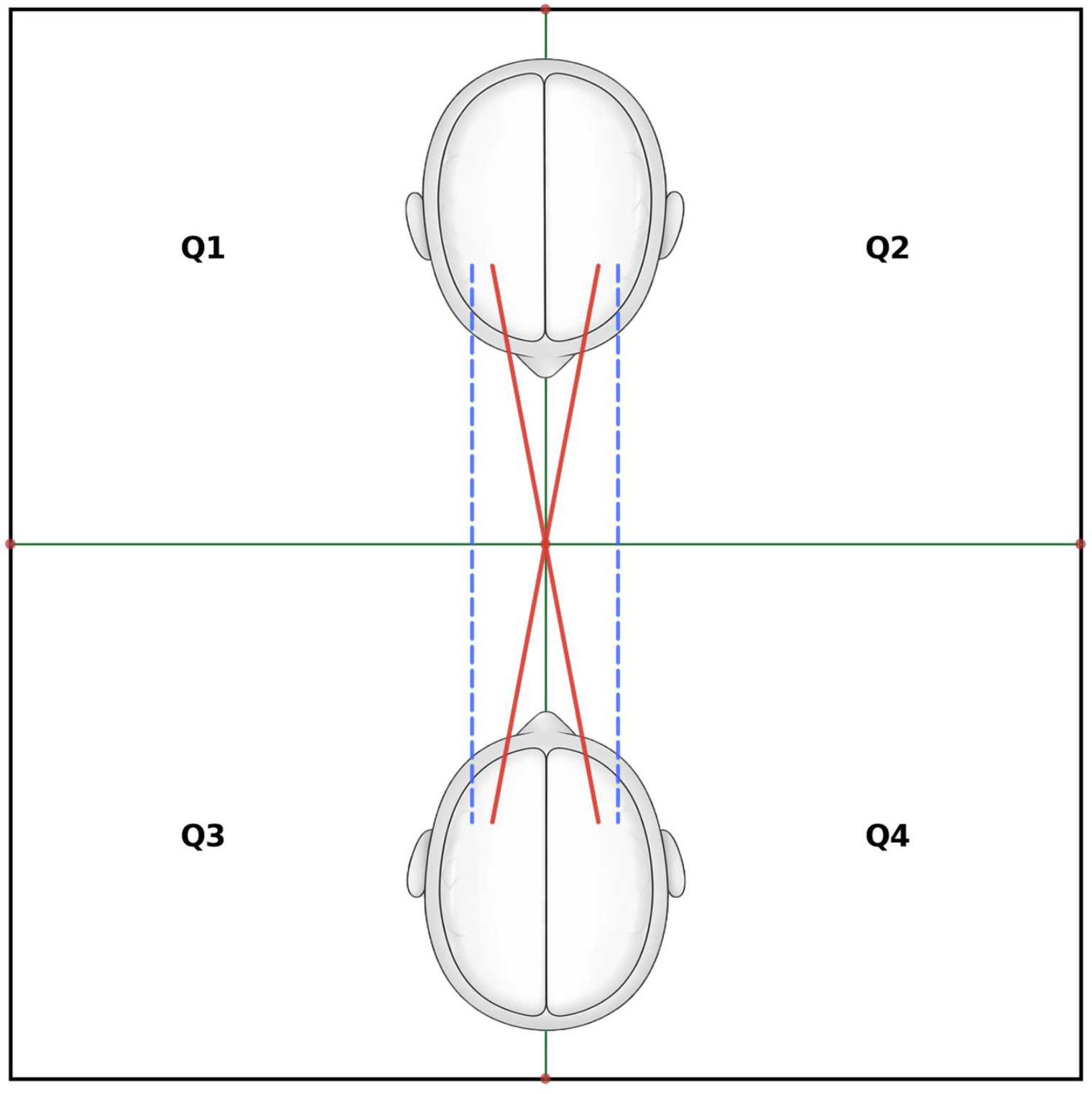
Tetradic geometry of embodied coordination. Apical view of a face-to-face dyad divided into four quadrants along two axes: Participant 1 occupies Quadrants 1 and 2 (top); Participant 2 occupies Quadrants 3 and 4 (bottom). The horizontal green axis denotes the self–other boundary; the vertical green axis denotes the shared left–right field boundary onto which each participant’s hemispheric midline is projected. Homologous axes (Q1–Q4, Q2–Q3; red solid lines) connect the same hemispheres across partners, corresponding to contralateral alignment in shared space (opposite sides of the shared left–right field): the geometric substrate of *Matching*. Cross-hemisphere axes (Q1–Q3, Q2–Q4; blue dashed lines) connect opposite hemispheres, corresponding to ipsilateral alignment in shared space (the same side of the shared left–right field): the geometric substrate of *Mirroring*. This fourfold organization defines a biaxial neural topology that maps hemispheric (left–right) and interpersonal (self–other) relations within a geometric spatiotemporal frame. Adapted from Olarewaju (2021).

Within this fourfold geometry, two fundamental coordination modes emerge within an interpersonal field (see Figure 2 for the geometric model of dyadic exchange; see Figure 1 for the tetradic organization of the dyadic field). *Mirroring*, characterized by movement alignment on the same side of shared interpersonal space and cross-hemispheric inter-brain pairing, reflects automatic sensorimotor resonance: low-inhibition coupling in which the partner’s action is reproduced along the shared visual-motor plane. In a face-to-face dyad, *mirroring* preserves the spatial reflection of the partner’s movement, but because each hemisphere controls the contralateral (left ↔ right) body side, this produces cross-hemispheric interbrain pairing (e.g., left precentral gyrus in one person with right precentral gyrus in the other).

**Figure 2.**
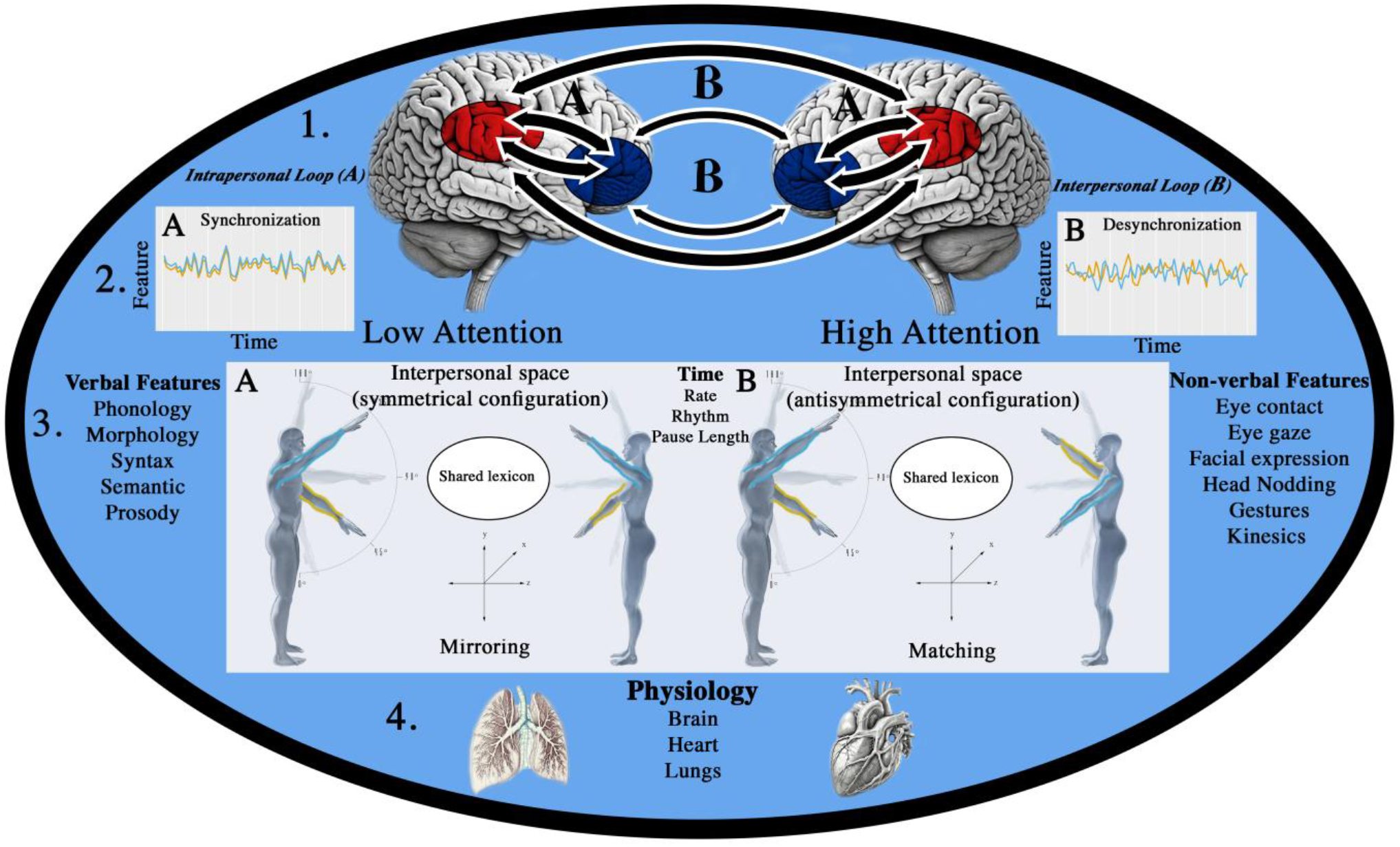
Geometric model of multiscale dyadic coordination. This side-view schematic embeds a dyadic entity in a tetrad, situating brain and body dynamics along orthogonal axes of self/other, left/right, and internal/external. (1) Neural connectivity encompasses (1A) intrapersonal loops within individuals and (1B) interpersonal loops between individuals, linking frontal (blue) and temporoparietal (red) regions along a shared self–other axis. (2) These loops support (2A) synchronization and (2B) desynchronization of verbal and nonverbal features, reflected in oscillatory transitions between in-phase (symmetrical-*mirroring*) and anti-phase (antisymmetrical-*matching*) dynamics. (3) Attention–inhibition gating structures recurrence in interpersonal space: (3A) low-attention/disinhibited *Mirroring* aligns movements on the same side of shared space and is represented by opposite-side body and cross-hemispheric brain pairings (left–right/right–left); (3B) high-attention/inhibitory *Matching* aligns movements on opposite sides of shared space and is represented by same-side body and homologous brain pairings (left–left/right–right). (4) Temporal parameters govern both verbal and nonverbal exchange and reciprocally modulate cardiorespiratory physiology. Adapted from Olarewaju (2021).

*Matching*, characterized by movement alignment on opposite sides of shared interpersonal space and homologous inter-brain pairing, reflects controlled inhibition of the *mirrored* form: higher-order regulation in which each partner aligns their effector system with a spatial rule rather than with the partner’s movement. Here, the same anatomical (right ↔ right/left ↔ left) body-side response is preserved across people. Because the same body side is controlled by the same hemisphere in each person, *matching* corresponds to homologous interbrain hemisphere pairs (e.g., left precentral gyrus to left precentral gyrus).

The tetradic framework integrates across four large-scale brain networks whose roles in social coordination are well established. The mirror-neuron system, centred on the inferior frontal gyrus (pars opercularis), inferior parietal lobule, and premotor cortex, provides the sensorimotor substrate for automatic resonance and imitation (Iacoboni, 2009; Rizzolatti & Craighero, 2004). Inputs from the superior temporal sulcus convey the bottom-up social signals (gaze direction, gesture kinematics) that drive this system (Allison et al., 2000; Grossman et al., 2000). The salience network (anterior insula, dorsal anterior cingulate cortex) regulates attentional allocation and the inhibitory gating of automatic responses, determining at each moment whether imitation is expressed or suppressed as a function of social context (Menon, 2011; Seeley et al., 2007). Frontoparietal regions mediate flexible, top-down executive monitoring and role-appropriate behavioural adjustment (Cole et al., 2013; Vincent et al., 2007). The default mode network (medial prefrontal cortex, posterior cingulate cortex, angular gyrus) instantiates the self–other relational frame from which the polarity of *Mirroring* and *Matching* derives its meaning, merging the present interaction within internal generative models of “me” and “you” (Buckner et al., 2008; Friston, 2010; Heyes, 2011; Raichle et al., 2001).

When embedded in the four-quadrant (tetradic) geometry, intra- and interpersonal dynamics through these multiscale networks form an integrated top-down/bottom-up attentional loop that alternates between stability (*Mirroring*) and controlled variability (*Matching*) across scales of space and time (Auffray & Nottale, 2008; Kelso, 1995; Olarewaju, 2021).

This work has four aims. First, we establish a robust, multilevel behavioural characterization of role asymmetry in attention–inhibition gating. Second, we identify the tonic neural correlates of the Condition dimension and determine whether this neural pattern is role-invariant or role-differentiated, thereby bridging the behavioural and neural accounts of attention–inhibition gating. Third, we extend this analysis across roles to characterize the phasic, time-resolved neural signatures of *Mirroring* and *Matching*, testing how the Condition effect is expressed around response-period onset and resolving the initiation–response–feedback cycle of leader–follower coordination. Fourth, we decompose the inter-brain information structure, isolating synergistic, unique, and redundant components as indices of the distributed predictive architecture of leader–follower coordination.

We addressed these aims using a dual-brain electroencephalography (EEG) hyperscanning paradigm in which healthy dyads completed a leader–follower sensorimotor coordination task designed to dissociate *Mirroring* from *Matching* within a 2 × 2 (Role × Condition) within-dyad factorial design. We use complementary analytic levels: (1) mixed-effects tonic ROI × band models capturing sustained network activation across the task; (2) spatiotemporal cluster permutation tests and frequency–space analyses capturing phasic, event-locked dynamics; (3) Integrated Information Decomposition (ΦID) (Chidichimo et al., 2026; Luppi et al., 2022) decomposing inter-brain dependencies into synergistic (sts), leader-unique (xtx), follower-unique (yty), and redundant (rtr) components. A combined behavioural cohort (the current hyperscanning sample plus an independent behavioural study conducted under equivalent procedures) was additionally analyzed to assess the replicability of the primary behavioural effects (see Glossary for conceptual and methodological terms used throughout the article).

### Glossary

| **Conceptual Terms** | **Definitions** |
|---|---|
| Tetradic Framework | The tetrad forms a fourfold relational geometry that structures dyadic interaction through ordered opposition (e.g., polarity). This architecture defines the left-right/self-other embedding of interpersonal space and time by providing a formal scaffold for understanding role asymmetry (leading and following) and its expression through (top-down controlled) antisymmetrical (*matching*) and (bottom-up automatic) symmetrical (*mirroring*) dynamics of face-to-face coordination (Kelso, 1995; Olarewaju, 2021). |
| Tetradic Brain–Body Alignment | In a face-to-face dyad, *Mirroring* pairs opposite anatomical body sides (left–right/right–left). The movements occupy the same side of shared space and correspond to cross-hemispheric pairing across the two brains. *Matching* pairs the same anatomical body sides (left–left/right–right). The movements occupy opposite sides of shared space and correspond to homologous hemisphere pairing. |
| Scale-Dependent Organization | A principle derived from scale-relativity theory (Auffray & Nottale, 2008), stating that relational structure is conserved across levels of organization while its geometric expression varies with scale through fractal-like patterns. In this work, the tetradic form is preserved across neural, behavioural, and interpersonal levels without implying mechanistic identity or scale collapse. |
| Metastability | A principle of brain–body coordination in which systems dynamically balance stability and flexibility, enabling rapid switching between coordination states (Kelso, 1995). In the context of this work, it describes how dyads move between stable (symmetric) *mirroring* and variable (antisymmetric) *matching* across leader–follower asymmetry. |
| Attention–Inhibition Gating | A regulatory principle describing how social interaction balances automatic and controlled behaviour. In the tetradic framework, this balance is modulated by attention and inhibition, which structures the role asymmetries of leading and following across neural, behavioural, and interpersonal levels. |
| **Statistical and Methodological Terms** |  |
| Region of Interest (ROI) | A predefined cortical region used for analysis, defined here using the Desikan–Killiany atlas. |
| Boundary Element Model (BEM) | A forward model used in EEG source reconstruction to estimate how electrical potentials propagate through the head. |
| Dynamic Statistical Parametric Mapping (dSPM) | A noise-normalized EEG inverse solution that estimates cortical source activity as standardized values. |
| Linear Mixed-Effects Model (LMM) | A statistical model that accounts for both fixed effects (e.g., role, condition) and random effects (e.g., participant, dyad) in repeated-measures data. |
| Likelihood-Ratio Test (LRT) | A test used to compare nested statistical models to determine whether the inclusion of additional parameters significantly improves model fit. |
| Spatiotemporal Cluster Permutation Testing | A nonparametric statistical approach that identifies significant effects in EEG data by clustering adjacent samples in time, space, and/or frequency. This analysis reduces the multiple-comparisons problem while preserving temporal and spatial structure. |
| Space–Frequency Analysis | A method in which EEG signals are decomposed into canonical frequency bands (theta, alpha, beta, gamma). The power in each band is averaged over the entire trial duration, and effects are then assessed across brain regions (space) and frequency bands. |
| Integrated Information Decomposition (ΦID) | A framework that decomposes joint predictive information from paired past variables to paired future variables into non-overlapping informational components reflecting distinct modes of contribution (synergy, unique-X, unique-Y, and redundancy). |
| Synergy (ΦID-sts) | Information about the joint future that is available only when both pasts are considered together, reflecting emergent dyadic integration.<br>*Note:* In this work, synergy specifies the dyad as a joint predictive system; synergistic dynamics therefore reorganize both global coordination and local differentiation, altering individual contributions (unique-X, unique-Y), shared structure (redundancy), and the system's capacity to adapt. |
| Unique-X (ΦID-xtx) | Predictive information about the joint future that is unique to the past of the Leader (*X*) after conditioning on the past of the Follower (*Y*). |
| Unique-Y (ΦID-yty) | Predictive information about the joint future that is unique to the past of the Follower (*Y*) after conditioning on the past of the Leader (*X*). |
| Redundancy (ΦID-rtr) | Information shared across both the Leader's and Follower's past states about the joint future was estimated using the minimum mutual information redundancy function (MMI). |

## 2. Results

Thirty-two healthy adults (16 dyads) completed a face-to-face leader–follower sensorimotor coordination task during simultaneous dual-stream EEG hyperscanning, constituting the primary neurobehavioural cohort (*N*RT = 32; *N*EEG = 30). A previously recorded behavioural-only sample (Olarewaju (2021), Experiment 1A; *N*RT = 32), collected using the same task structure, was combined with the current hyperscanning dataset to form a merged behavioural cohort (*N*RT = 64) for replication analyses. Both cohorts completed a 2 × 2 within-subject design crossing Role (Leader/Follower) × Condition (*Mirroring*/*Matching*). Reaction-time (RT) differences between conditions index attention–inhibition gating at the behavioural level; dual-stream EEG analyses then extend this framework across tonic, phasic, and information-theoretic levels of neural organization. Results are presented in the following order: behavioural asymmetry (#2.1), an overview of the neural analytic strategy (#2.2), tonic EEG activity (#2.3), phasic EEG dynamics (#2.4), inter-brain information decomposition via ΦID (#2.5), and a cross-method synthesis (#2.6). All figures, tables, and supplementary materials are referenced throughout.

### 2.1 Behavioural Asymmetry: Role-Dependent Attention–Inhibition Gating

#### 2.1.1 Condition effect and role asymmetry at the aggregate level (N_RT_ = 32)

A 2 × 2 repeated-measures ANOVA on condition-wise participant median RTs (*N*RT = 32) revealed a significant main effect of Condition (*F*(1, 31) = 18.25, *p* = .0002), confirming that RTs differed systematically between the *Mirroring* and *Matching* conditions. The main effect of Role was not significant (*F*(1, 31) = 2.95, *p* = .096), nor was the Role × Condition interaction (*F*(1, 31) = 1.93, *p* = .174; see Table 1). These results indicate that the role-specific asymmetry was not yet detectable at the aggregate median level in this cohort. Bonferroni-corrected post-hoc comparisons of participant median RTs revealed that Followers responded significantly faster during *Mirroring* than *Matching* (*t*(31) = 5.83, *p*adj < .001, *d* = 1.03; see Figure 3a). For Leaders, the corresponding difference was negligible and non-significant (*t*(31) = 1.24, *p*adj = .896, *d* = 0.22). Model-based RT differences for these contrasts are reported separately in Table 2. This pattern of differential condition sensitivity (strong in Followers, absent in Leaders) constitutes the first behavioural signature of role-specific attention–inhibition gating. The overall Mirroring RT advantage persisted at the dyad level (Supplementary Section S1.3; Table S8).

**Figure 3.**
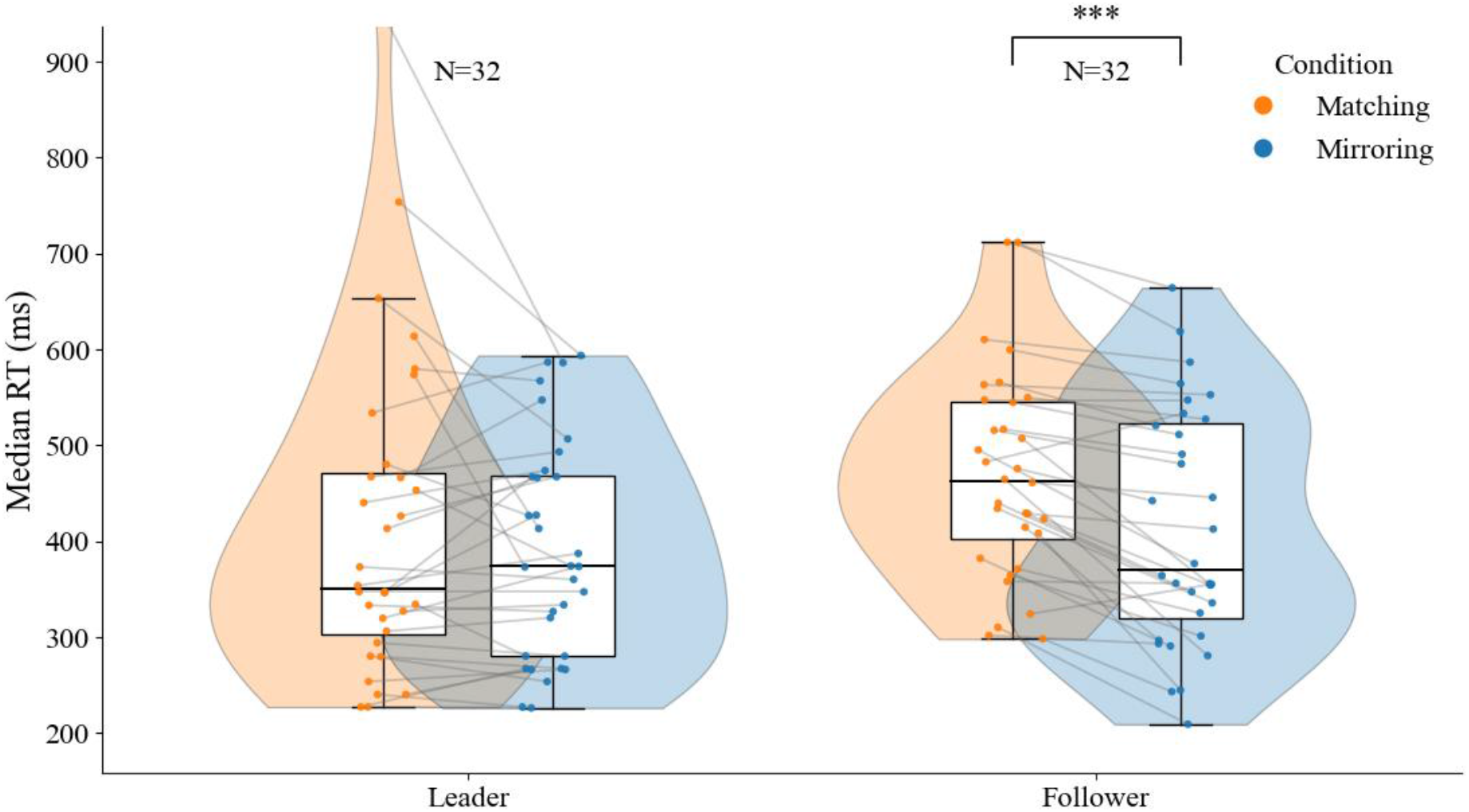
Behavioural Polarity (NRT = 32) *Note*. Violin plots show the distributions of participant median response times for Leaders and Followers across the *Matching* and *Mirroring* conditions. Boxplots indicate the median and interquartile range. Individual data points represent participant medians. Grey connecting lines link each participant’s *Matching* and *Mirroring* values, illustrating within-participant changes across conditions. Asterisks denote Bonferroni-adjusted significance of paired comparisons (pₐ*d*ⱼ < 0.05; **pₐ*d*ⱼ < 0.001).

**Table 1.**
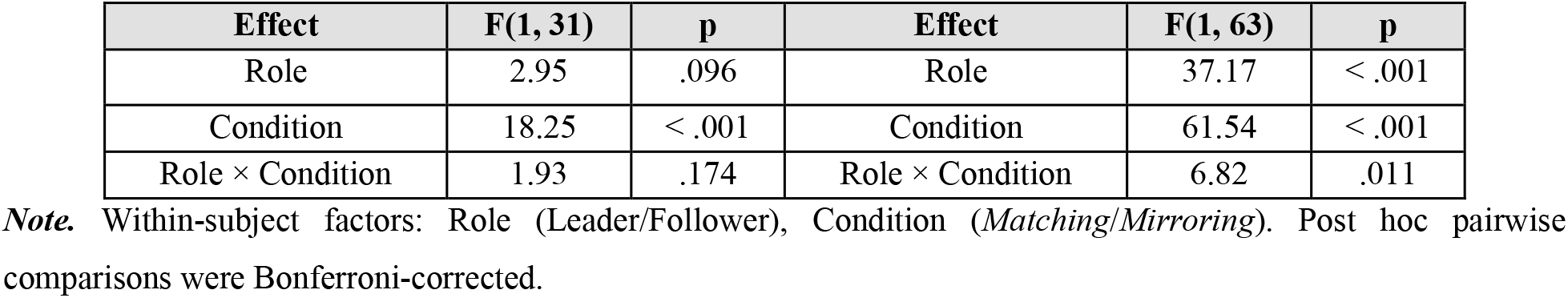
2 × 2 Repeated-Measures ANOVA on Participant Median RTs: Primary Cohort (N_RT_ = 32) and Replication Cohort (N_RT_ = 64)

| **Effect** | **F(1, 31)** | **p** | **Effect** | **F(1, 63)** | **p** |
|---|---|---|---|---|---|
| Role | 2.95 | .096 | Role | 37.17 | < .001 |
| Condition | 18.25 | < .001 | Condition | 61.54 | < .001 |
| Role × Condition | 1.93 | .174 | Role × Condition | 6.82 | .011 |

***Note.*** Within-subject factors: Role (Leader/Follower), Condition (*Matching*/*Mirroring*). Post hoc pairwise comparisons were Bonferroni-corrected.

**Table 2.** Condition Contrasts by Role: Model-Based RT Differences and Participant-Median Tests in the Primary (N_RT_ = 32) and Replication (N_RT_ = 64) Cohorts.

| **Contrast** | **N** | **Δ (ms)** | **95% CI (ms)** | **t** | **p_adj** | **Cohen’s d** |
|---|---|---|---|---|---|---|
| Follower: Mirroring − Matching | 32 | −59.7 | [−77.9, −40.6] | 5.83 | < .001 | 1.03 |
| Leader: Mirroring − Matching | 32 | −6.1 | [−29.5, 18.7] | 1.24 | .896 | 0.22 |
| Follower: Mirroring − Matching | 64 | −69.1 | [−93.9, −42.4] | 9.60 | < .001 | 1.20 |
| Leader: Mirroring − Matching | 64 | −26.6 | [−46.0, −5.8] | 2.82 | .026 | 0.35 |

***Note.*** Δ and 95% CIs are model-based differences in milliseconds (*Mirroring − Matching*), estimated from fitted trial-level mixed-effects models and back-transformed from log RT. The t statistics, Bonferroni-adjusted p values, and Cohen’s d come from paired comparisons of participant median RTs; these statistics are not tests of the model-based Δ estimates.

#### 2.1.2 Trial-level mixed-effects modelling reveals a Role × Condition interaction (N_RT_ = 32)

To assess whether the role-specific asymmetry was detectable at the level of individual trials and to account for within-participant, within-dyad, and across-condition variance, a linear mixed-effects model (LMM) was fitted to log-transformed trial-level RTs. Fixed effects included Role, Condition, and their interaction; random effects included participant intercepts, slopes were supported by the data, and a dyad-level variance component. A significant Role × Condition interaction emerged (β = -0.123, SE = 0.043, z = -2.86, p = .004; see Table 3), confirming that Followers’ differential condition sensitivity is reliable once trial-level variability and nested dependencies are modelled explicitly. Inclusion of the interaction substantially improved model fit relative to a model without it (likelihood-ratio test: 2ΔLL = 158.42, df = 1, p < .001). Back-transformed predicted cell means were: Follower-*Matching*: 461 ms, Follower-*Mirroring*: 402 ms, Leader-*Matching*: 400 ms, Leader-*Mirroring*: 394 ms, yielding a Follower *Mirroring* advantage of Δ = -59.7 ms (95% CI [-77.9, -40.6]; ratio = 0.871) versus a Leader effect of Δ = -6.1 ms (95% CI [-29.5, 18.7]; ratio = 0.985). Bayesian LMM estimates corroborated these results: the Follower contrast yielded a posterior P(Δ < 0) = 1.000, while the Leader contrast placed 86.1% of posterior mass within the ±5% region of practical equivalence (ROPE).

**Table 3.** Fixed Effects of the Trial-Level Linear Mixed-Effects Model (N_RT_ = 32; log-transformed RT, Gaussian likelihood)

| **Effect** | **β (log scale)** | **SE** | **z** | **p** | **95% CI** |
|---|---|---|---|---|---|
| Intercept | 6.025 | 0.042 | 142.85 | < .001 | [5.942, 6.107] |
| Role (Follower − Leader) | 0.081 | 0.043 | 1.85 | .064 | [−0.005, 0.166] |
| Condition (Mirroring − Matching) | −0.077 | 0.017 | −4.40 | < .001 | [−0.111, −0.043] |
| Role × Condition | −0.123 | 0.043 | −2.86 | .004 | [−0.208, −0.039] |
| Interaction LRT: 2ΔLL = 158.42, df = 1, p < .001 |  |  |  |  |  |

***Note.*** Sum-coded contrasts: Role (Leader = −0.5, Follower = +0.5), Condition (*Matching* = −0.5, *Mirroring* = +0.5). Random effects included participant intercepts, slopes were supported by the data, and a dyad-level variance component. β coefficients are on the log scale.

#### 2.1.3 Individual-level consistency of the Follower Mirroring advantage

Within-participant trial-level contrasts assessed how consistently the condition effect appeared across individuals. Twenty-five of 32 Followers (78%) showed statistically significant within-participant *Mirroring*-faster contrasts (Bonferroni-adjusted α = .025), and 28 of 32 (88%) were directionally faster during *Mirroring*, many with medium-to-large within-person effect sizes. Leaders showed mixed individual-level directions: 17 were faster during *Mirroring* and 15 during *Matching* (Figure 3b). These data confirm that the Follower advantage reflects a robust individual-level tendency and that Leaders, as a group, do not exhibit a consistent condition preference at this sample size.

**Figure 3b.**
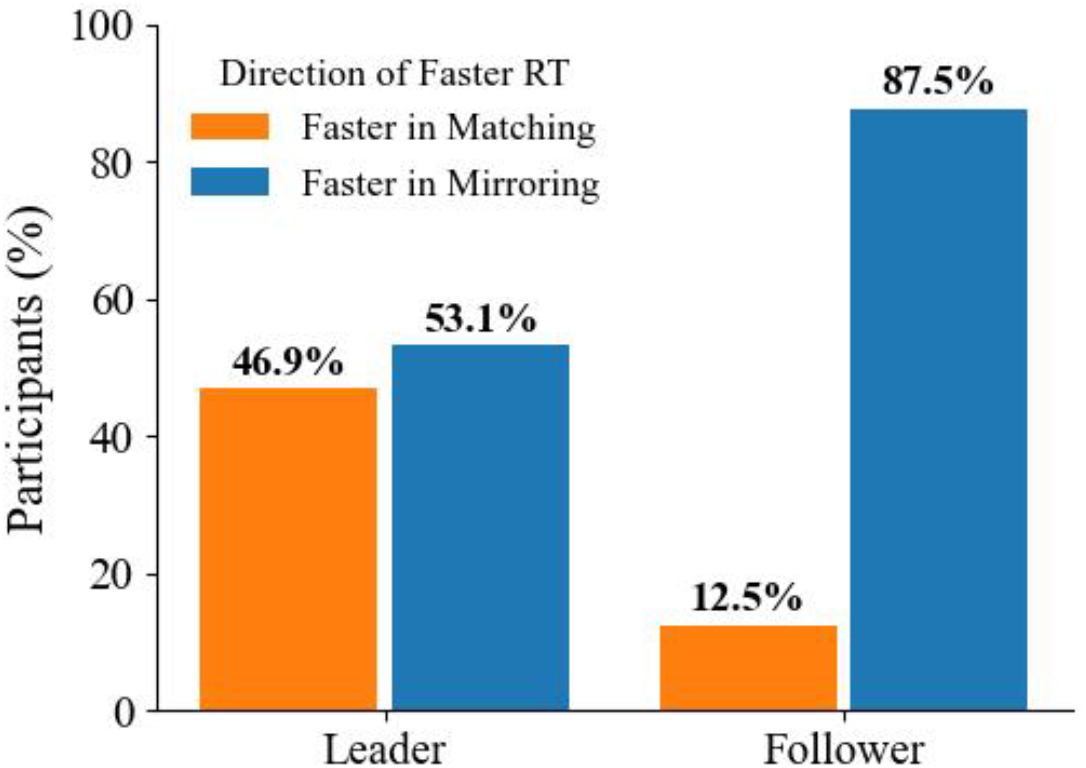
Individual-Level Behavioural Polarity by Role (NRT = 32) *Note.* Bars indicate the percentage of participants whose mean response time was faster in *Mirroring* or *Matching*. This visualization summarizes the direction of the individual-level differences, independent of their magnitude.

#### 2.1.4 Replication in the combined behavioural cohort (N_RT_ = 64)

To assess the generalizability of the role-differentiated pattern, the hyperscanning cohort was combined with the independent sample. In this merged cohort (*N*RT = 64), a 2 × 2 repeated-measures ANOVA revealed significant main effects of Role (*F*(1, 63) = 37.17, *p* < .001, η^2^*p* = .37) and Condition (*F*(1, 63) = 61.54, *p* < .001, η^2^*p* = .49), with a small but significant Role × Condition interaction (*F*(1, 63) = 6.82, *p* = .011, η^2^*p* = .10; see Table 1). Post-hoc contrasts of participant median RTs confirmed a large Follower *Mirroring* advantage (*d* = 1.20) and a smaller but now-significant Leader effect (*d* = 0.35; see Figure 3c). The corresponding model-based Mirroring − Matching RT differences were −69.1 ms for Followers and −26.6 ms for Leaders (Table 2). Equivalence testing (TOST) indicated that the Leader effect failed both the ±5% log-scale and ±25 ms raw-scale equivalence bounds, confirming that it is neither practically null nor as large as the Follower effect. Bayesian models corroborated the directionality of both effects (Follower: P(Δ < 0) = 1.000; Leader: P(Δ < 0) = .994), though the Leader contrast failed the ±5% ROPE criterion (P = .116), leaving its practical significance equivocal. These replication data confirm the primary behavioural finding while underscoring that the role asymmetry in condition sensitivity is substantially more pronounced in Followers than in Leaders.

**Figure 3c.**
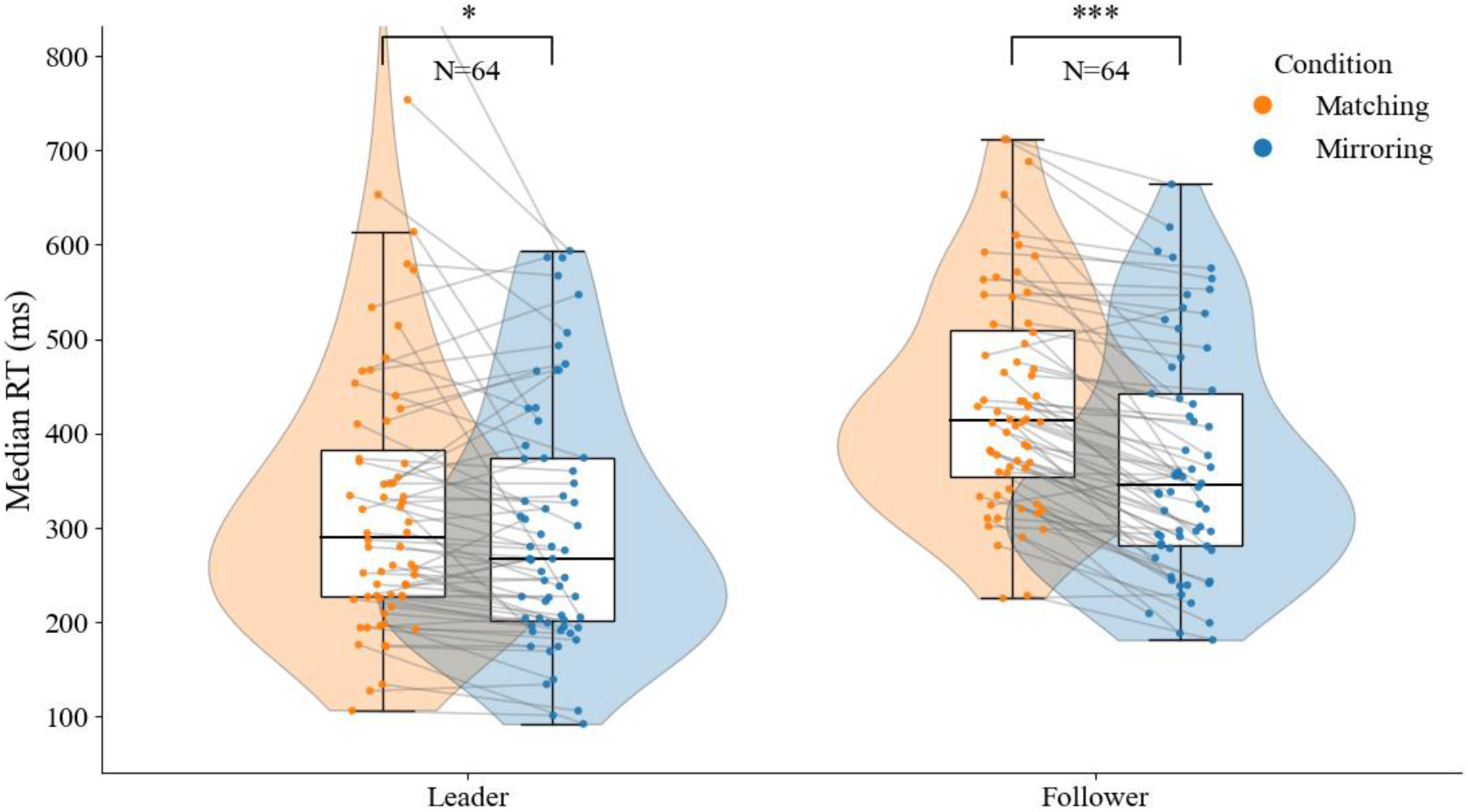
Behavioural Polarity (NRT = 64) *Note.* Violin plots show the distributions of participant median response times for Leaders and Followers across the *Matching* and *Mirroring* conditions in the combined cohort. Boxplots indicate the median and interquartile range. Individual data points represent participant medians. Grey connecting lines link each participant’s *Matching* and *Mirroring* values, emphasizing within-participant shifts across conditions. Asterisks denote Bonferroni-adjusted significance of paired comparisons (*p*ₐ*d*ⱼ < 0.05; \*\**p*ₐ*d*ⱼ < 0.001).

### 2.2 Neural Correlates of Attention–Inhibition Gating: Analytic Overview

The dual-stream EEG hyperscanning analyses (*N***_EEG_** = 30) preserve the Role × Condition contrast established behaviourally and proceed in two stages. The first stage examines tonic neural activity using mixed-effects models that identify stable, trial-averaged patterns of region- and frequency-specific neural energy across experimental conditions. The second stage examines phasic dynamics through two complementary time-resolved approaches: (1) spatiotemporal cluster permutation testing, which identifies condition-related differences around response-period onset and their resolution into role-specific temporal dynamics; (2) inter-brain Integrated Information Decomposition (ΦID), which decomposes the joint predictive structure of the dyad into atomic informational components.

Together, these analyses target the neural expression of attention–inhibition gating at multiple levels of temporal and spatial organization, demonstrating how the behavioural polarity between *Mirroring* and *Matching* is grounded in, and reorganized across, the dual-brain system. Details of EEG preprocessing, source reconstruction, parcellation, and analytic pipelines are reported in the Methods.

### 2.3 Tonic Neural Activity: A Widespread Condition Effect

Trial-level mixed-effects models fitted to ROI × frequency-band power estimates from individual epochs (408 total; 68 Desikan–Killiany ROIs × 6 frequency bands) with RT entered as a covariate revealed a broadly distributed Condition effect. Of the 408 features examined, 279 survived false discovery rate (FDR) correction (*q* < .05, Benjamini–Hochberg). Of these, 257 features showed greater neural activity during *Matching* than *Mirroring*, and only 22 showed the reverse; *Matching*-related effects were thus strongly dominant (see Figure 4; Table 4; Supplementary Table S4). These *Matching*-dominant features were concentrated within the frontoparietal and salience (cingulo-opercular) networks regions implicated in precision-weighted executive control and inhibitory gating alongside beta- and gamma-range modulation in the posterior cingulate and medial orbitofrontal cortices, consistent with engagement of default-mode network components implicated in contextual self–other evaluation during controlled coordination. In a dyad-level RT-adjusted sensitivity analysis, Condition estimates remained predominantly Matching-positive, although no feature survived FDR correction (Supplementary Section S5.1; Table S8).

**Figure 4.**
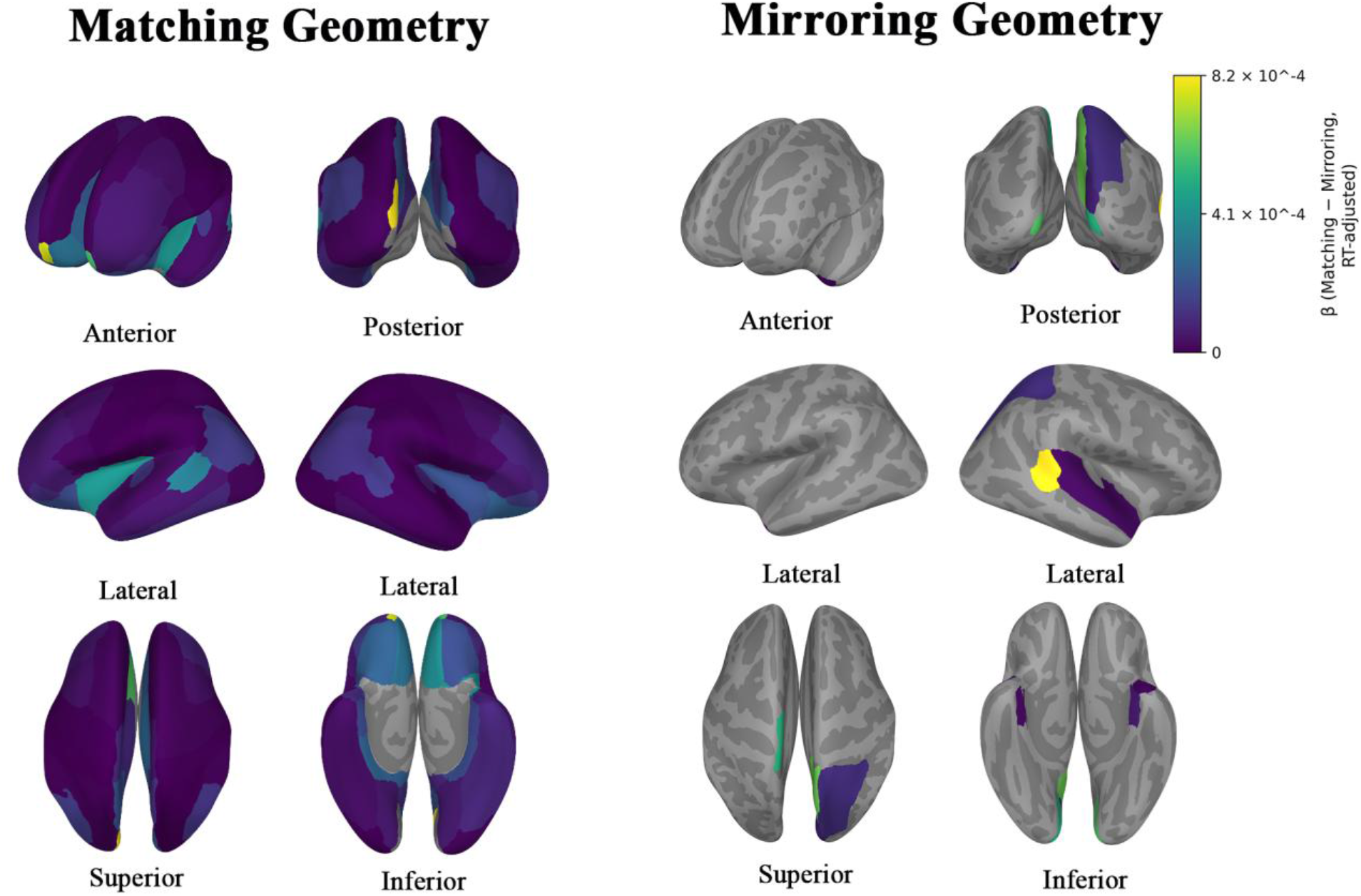
Neural geometry of tonic coordination. Aggregated ROI × frequency-band effects from mixed-effects models were used to contrast the *Matching* and *Mirroring* conditions (FDR q < .05). Colour intensity of the brain maps reflects the magnitude of the RT-adjusted β coefficient across conditions. The brighter the colour, the larger the absolute effects. Here, average neural energetic demand differences are predominantly expressed across the frontoparietal control network and salience (cingulo-opercular) systems during *Matching*. In contrast, *Mirroring*-related effects are concentrated in midline default-mode regions. The maps summarize condition effects across roles. Role-specific patterns showed near-complete overlap for *Matching* > *Mirroring* (Jaccard ≈ 0.95), but more moderate overlap for *Mirroring* > *Matching* (Jaccard ≈ 0.57). Grey regions indicate ROIs that did not survive correction.

**Table 4.**
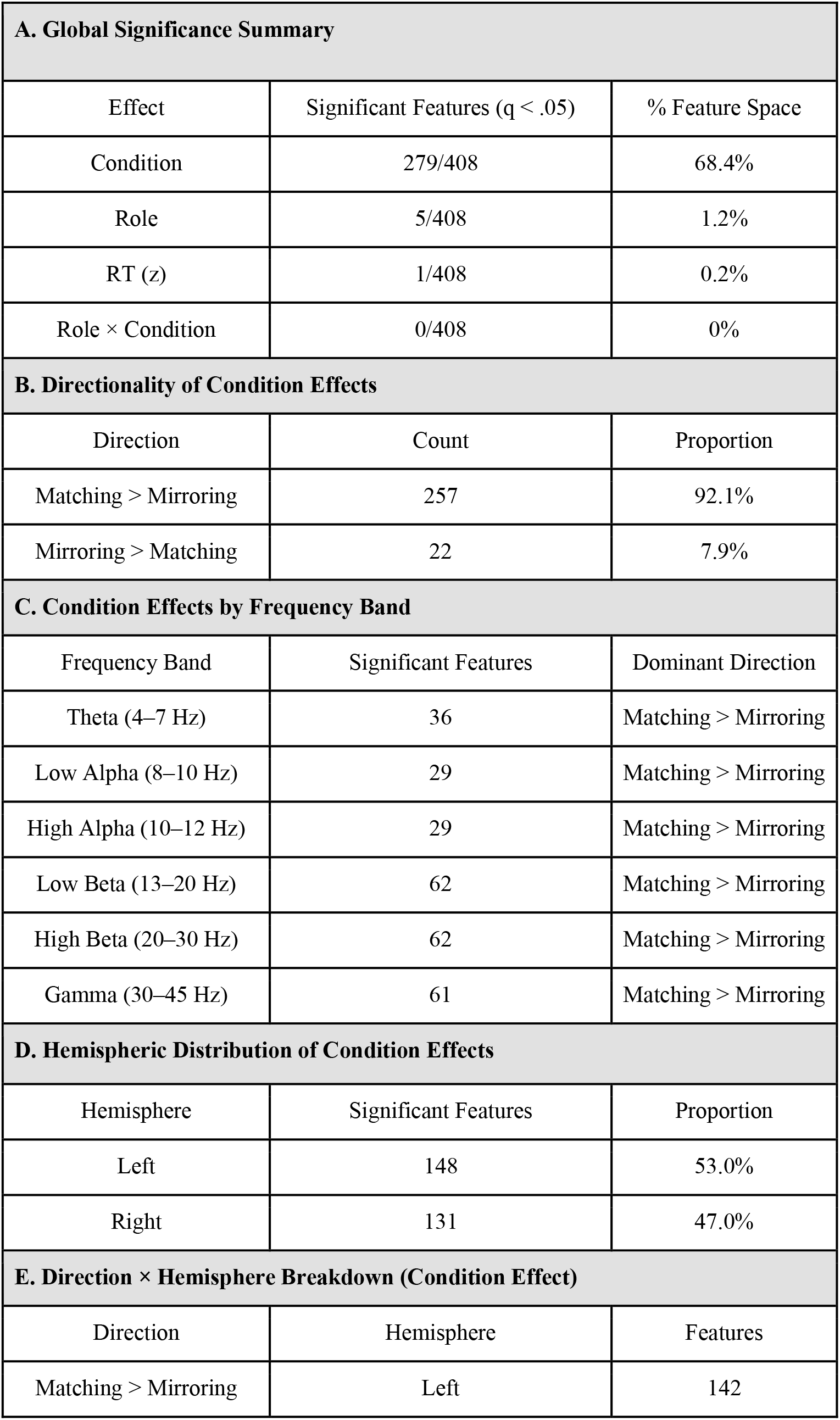
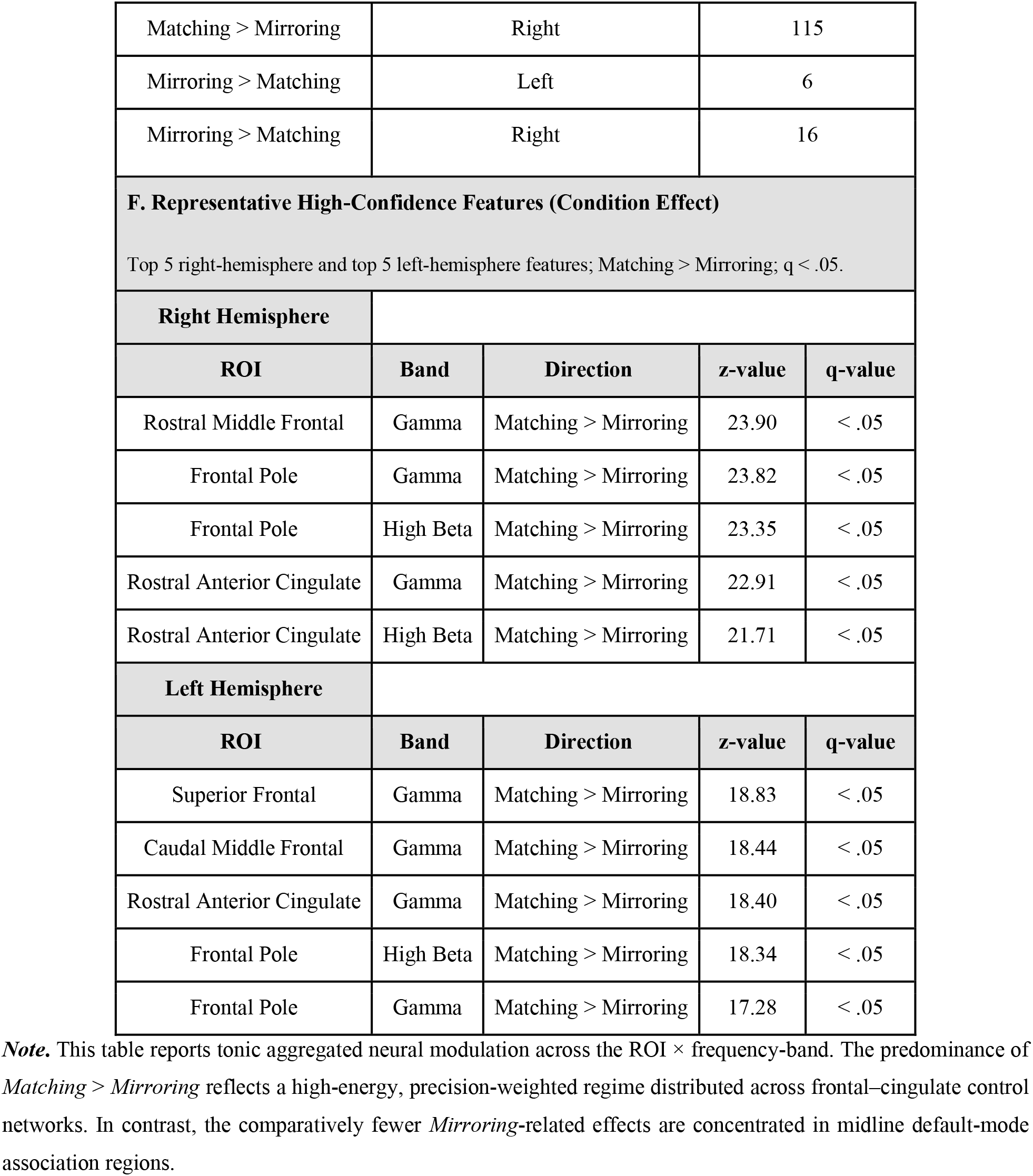
Mixed-Effects ROI × Frequency Band Models (RT-adjusted tonic effects)

| **A. Global Significance Summary** |  |  |  |  |
|---|---|---|---|---|
| Effect | Significant Features (q < .05) |  | % Feature Space |  |
| Condition | 279/408 |  | 68.4% |  |
| Role | 5/408 |  | 1.2% |  |
| RT (z) | 1/408 |  | 0.2% |  |
| Role × Condition | 0/408 |  | 0% |  |
| **B. Directionality of Condition Effects** |  |  |  |  |
| Direction | Count |  | Proportion |  |
| Matching > Mirroring | 257 |  | 92.1% |  |
| Mirroring > Matching | 22 |  | 7.9% |  |
| **C. Condition Effects by Frequency Band** |  |  |  |  |
| Frequency Band | Significant Features |  | Dominant Direction |  |
| Theta (4–7 Hz) | 36 |  | Matching > Mirroring |  |
| Low Alpha (8–10 Hz) | 29 |  | Matching > Mirroring |  |
| High Alpha (10–12 Hz) | 29 |  | Matching > Mirroring |  |
| Low Beta (13–20 Hz) | 62 |  | Matching > Mirroring |  |
| High Beta (20–30 Hz) | 62 |  | Matching > Mirroring |  |
| Gamma (30–45 Hz) | 61 |  | Matching > Mirroring |  |
| **D. Hemispheric Distribution of Condition Effects** |  |  |  |  |
| Hemisphere | Significant Features |  | Proportion |  |
| Left | 148 |  | 53.0% |  |
| Right | 131 |  | 47.0% |  |
| **E. Direction × Hemisphere Breakdown (Condition Effect)** |  |  |  |  |
| Direction | Hemisphere |  | Features |  |
| Matching > Mirroring | Left |  | 142 |  |
| Matching > Mirroring | Right |  | 115 |  |
| Mirroring > Matching | Left |  | 6 |  |
| Mirroring > Matching | Right |  | 16 |  |
| **F. Representative High-Confidence Features (Condition Effect)**<br>Top 5 right-hemisphere and top 5 left-hemisphere features; Matching > Mirroring; q < .05. |  |  |  |  |
| **Right Hemisphere** |  |  |  |  |
| **ROI** | **Band** | **Direction** | **z-value** | **q-value** |
| Rostral Middle Frontal | Gamma | Matching > Mirroring | 23.90 | < .05 |
| Frontal Pole | Gamma | Matching > Mirroring | 23.82 | < .05 |
| Frontal Pole | High Beta | Matching > Mirroring | 23.35 | < .05 |
| Rostral Anterior Cingulate | Gamma | Matching > Mirroring | 22.91 | < .05 |
| Rostral Anterior Cingulate | High Beta | Matching > Mirroring | 21.71 | < .05 |
| **Left Hemisphere** |  |  |  |  |
| **ROI** | **Band** | **Direction** | **z-value** | **q-value** |
| Superior Frontal | Gamma | Matching > Mirroring | 18.83 | < .05 |
| Caudal Middle Frontal | Gamma | Matching > Mirroring | 18.44 | < .05 |
| Rostral Anterior Cingulate | Gamma | Matching > Mirroring | 18.40 | < .05 |
| Frontal Pole | High Beta | Matching > Mirroring | 18.34 | < .05 |
| Frontal Pole | Gamma | Matching > Mirroring | 17.28 | < .05 |

***Note.*** This table reports tonic aggregated neural modulation across the ROI × frequency-band. The predominance of *Matching > Mirroring* reflects a high-energy, precision-weighted regime distributed across frontal–cingulate control networks. In contrast, the comparatively fewer *Mirroring*-related effects are concentrated in midline default-mode association regions.

Critically, this neural energetic pattern was largely invariant across Leader and Follower roles. Decomposition of simple effects using independently FDR-corrected role-specific models revealed near-complete spatial overlap in ROI × band features for the dominant contrast (*Matching* > *Mirroring*; Jaccard ≈ 0.95), with more moderate overlap observed for the weaker contrast (*Mirroring* > *Matching*; Jaccard ≈ 0.57); see Figure 4. Despite these differences in spatial extent, effect magnitudes were effectively identical across roles (median |Δβ| ≈ 0), indicating that the tonic demand of *Matching* versus *Mirroring* is encoded in a shared neural field. These results suggest that, at the tonic timescale, condition-related energetic demands are predominantly role-invariant, with only minor role-dependent variation in the spatial distribution of weaker effects. This role-invariance is addressed in the Discussion.

### 2.4 Phasic Neural Dynamics Across Conditions and Roles

To examine how the tonic Condition pattern is expressed across time, spatiotemporal cluster permutation tests (5,000 permutations; cluster-level FWER within each band × contrast map, α = .05) were applied to source-level data aligned to response-period onset (−1.0 to +1.0 s). Clusters were defined over contiguous time points and spatially adjacent ROIs, with neighbourhood structure derived from ROI centroid distances. A complementary frequency–space cluster analysis (collapsing across time) was conducted as a corroborating check.

Phasic analyses identified greater source-level power during *Matching* than *Mirroring* in significant condition clusters extending across the entire response-period-aligned epoch. Clusters were identified in the alpha (8–12 Hz), beta (13–30 Hz), and gamma (30–45 Hz) ranges, with cluster-level FWER *p* values ≤ .0122 (estimated mean *d*ᶻ ≈ 0.49–0.51; peak |*d*ᶻ| ≈ 0.80–0.91; see Figure 5, Table 5, Supplementary section S6, and Supplementary Table S5a). In the complementary space × frequency analysis, only Condition clusters survived (Supplementary Table S5b). Together, these findings show that *Matching* is associated with greater neural power than *Mirroring*, both in the time-averaged tonic analysis and response-period-aligned phasic clusters. To test whether the response-period-aligned clusters contained temporal variation beyond this sustained difference, we centered each participant’s Condition contrast across the epoch. Significant high-beta and gamma clusters were identified in the centered data within their respective maps, indicating limited time-varying Condition modulation not captured by the tonic average (Supplementary Section S6.5). The robustness of these Condition clusters was examined in RT-controlled and across-map sensitivity analyses (Supplementary Sections S6.3–S6.4; Table S7).

**Figure 5.**
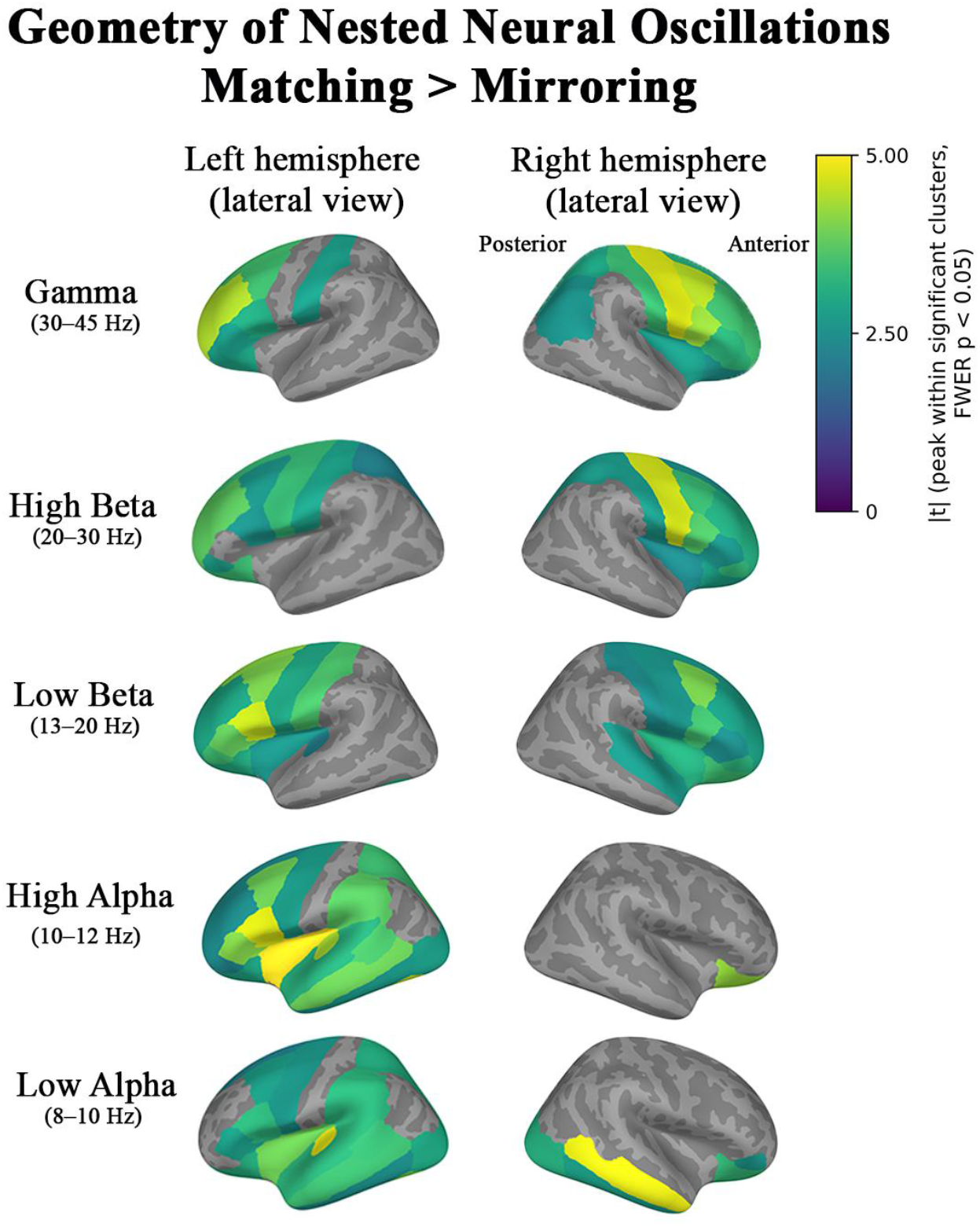
Neural geometry of phasic coordination. Spatiotemporal cluster-based permutation analysis was applied to source-level data aligned to response-period onset (−1.0 to +1.0 s; 5,000 permutations), contrasting *Matching* and *Mirroring* conditions across ROIs and frequency bands (cluster-level FWER *p* < .05). A complementary space × frequency analysis was conducted separately. Colour intensity of the brain maps reflects the peak absolute *t*-value (|*t*|) for each ROI within significant spatiotemporal clusters, indicating effect magnitude independent of sign. Direction of effects and anatomical coverage are reported in Supplementary Tables S5a and S5b. Grey regions indicate ROIs that did not participate in a significant spatiotemporal cluster.

**Table 5.** Spatiotemporal Cluster Statistics (Condition-based Phasic Effects)

| **Contrast** | **Direction of Effect** | **Frequency Band** | **Time Window (ms)** | **Cluster p (FWER)** | **Mean dz** | **Peak \|dz\|** | **Dominant Neural Regions (compressed from ROI lists)** |
|---|---|---|---|---|---|---|---|
| Condition | Matching > Mirroring | Low Alpha (8–10 Hz) | −1000 → 1000 | 0.0004 | 0.50 | 0.91 | Lateral occipital, fusiform, inferotemporal, orbitofrontal, anterior cingulate, parahippocampal |
| Condition | Matching > Mirroring | High Alpha (10–12 Hz) | −1000 → 1000 | 0.0014 | 0.49 | 0.84 | Orbitofrontal, anterior cingulate, middle frontal, temporal association, lateral occipital |
| Condition | Matching > Mirroring | Low Beta (13–20 Hz) | −1000 → 1000 | 0.0076 | 0.49 | 0.88 | Anterior cingulate, orbitofrontal, premotor, postcentral |
| Condition | Matching > Mirroring | High Beta (20–30 Hz) | −1000 → 1000 | 0.0122 | 0.50 | 0.85 | Anterior cingulate, superior parietal, premotor, postcentral |
| Condition | Matching > Mirroring | Gamma (30–45 Hz) | −1000 → 1000 | 0.0090 | 0.51 | 0.80 | Medial and lateral frontal, anterior and posterior cingulate, superior parietal |

***Note.*** Cluster-based permutation statistics were computed on epochs aligned to response-period onset (−1.0 to +1.0 s; 5,000 permutations). Reported p-values are cluster-level familywise error–corrected using the maximum cluster statistic across permutations within each band × contrast map (α = 0.05); no correction across maps was applied. Mean *d*<sup>z</sup> reflects the cluster-averaged standardized effect size, and Peak \|*d*<sup>z</sup>\| denotes the maximum local standardized effect within each cluster. Time windows indicate the contiguous temporal extent of each significant cluster.

#### 2.4.1 Role-specific Phasic Coordination: Initiation–Response–Feedback

To understand how these condition-related dynamics unfold across roles, we turn to role-specific spatiotemporal clusters (see Table 6; Supplementary Table S5a). Here, we detected a pre-onset sequence consistent with control processes around Leader responses (see Figure 6a). Before Leader response-period onset, theta (4–7 Hz) was observed (−763 to −275 ms, *p* = 0.0148, *dz* ≈ 0.51), predominantly in frontal midline, premotor, and orbitofrontal cortices, regions associated with cognitive control and conflict monitoring (Cavanagh & Frank, 2014). Low-alpha (−723 to −268 ms, *p* = 0.0122, *dz* ≈ 0.49) and high-alpha (−718 to −277 ms, *p* = 0.0402, *dz* ≈ 0.50) emerged almost concurrently in orbitofrontal, rostral cingulate, and temporal-pole regions. Given the task demands preceding Leader responses, phasic activity in these regions during these intervals is consistent with proactive task maintenance (Brinkman et al., 2014). Finally, a high-beta cluster (−459 to +208 ms, *p* = 0.0262, *dz* ≈ 0.46) appeared over frontoparietal cortices, consistent with the controlled initiation of action (Tzagarakis et al., 2010).

**Figures 6a–b.**
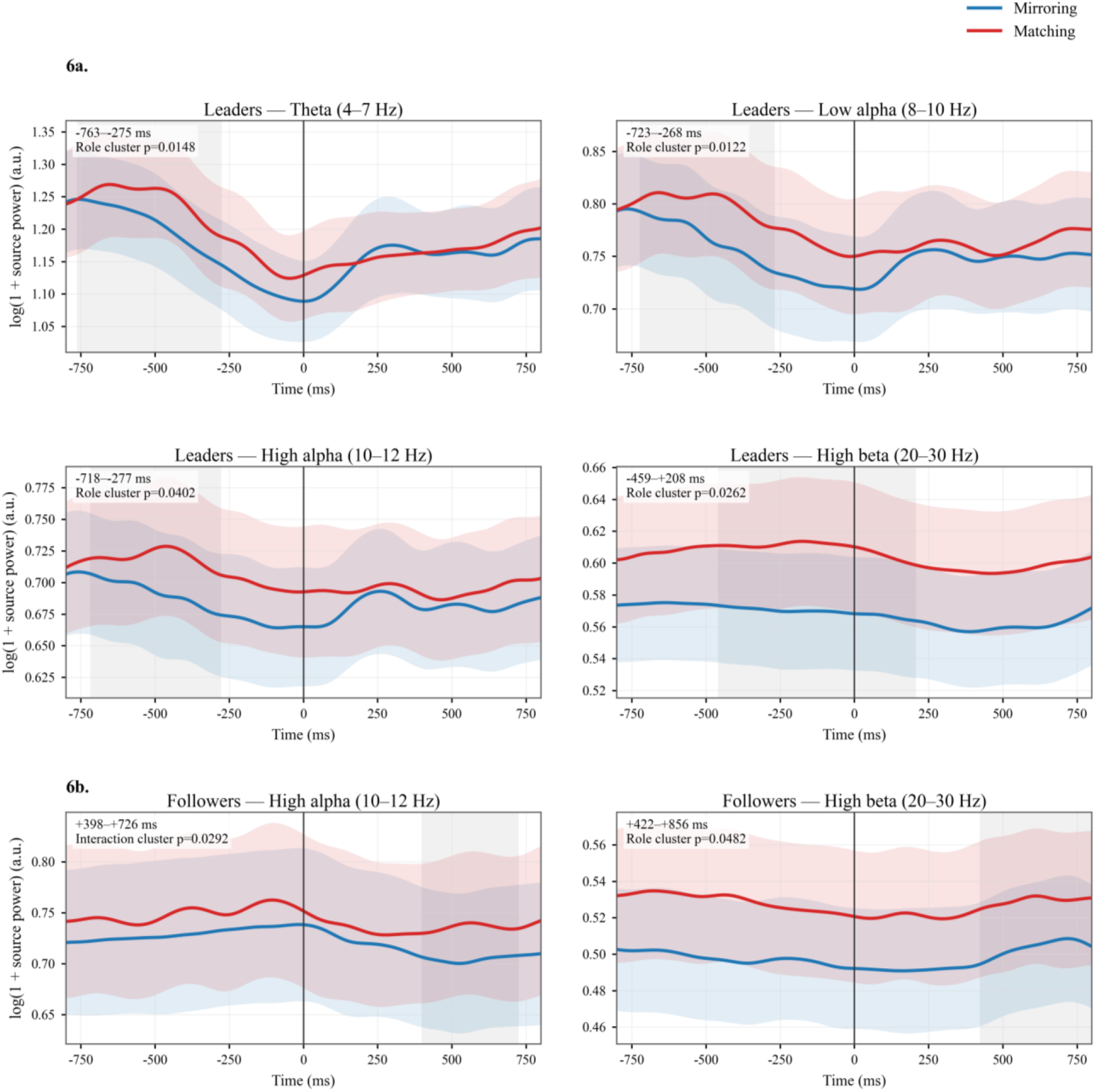
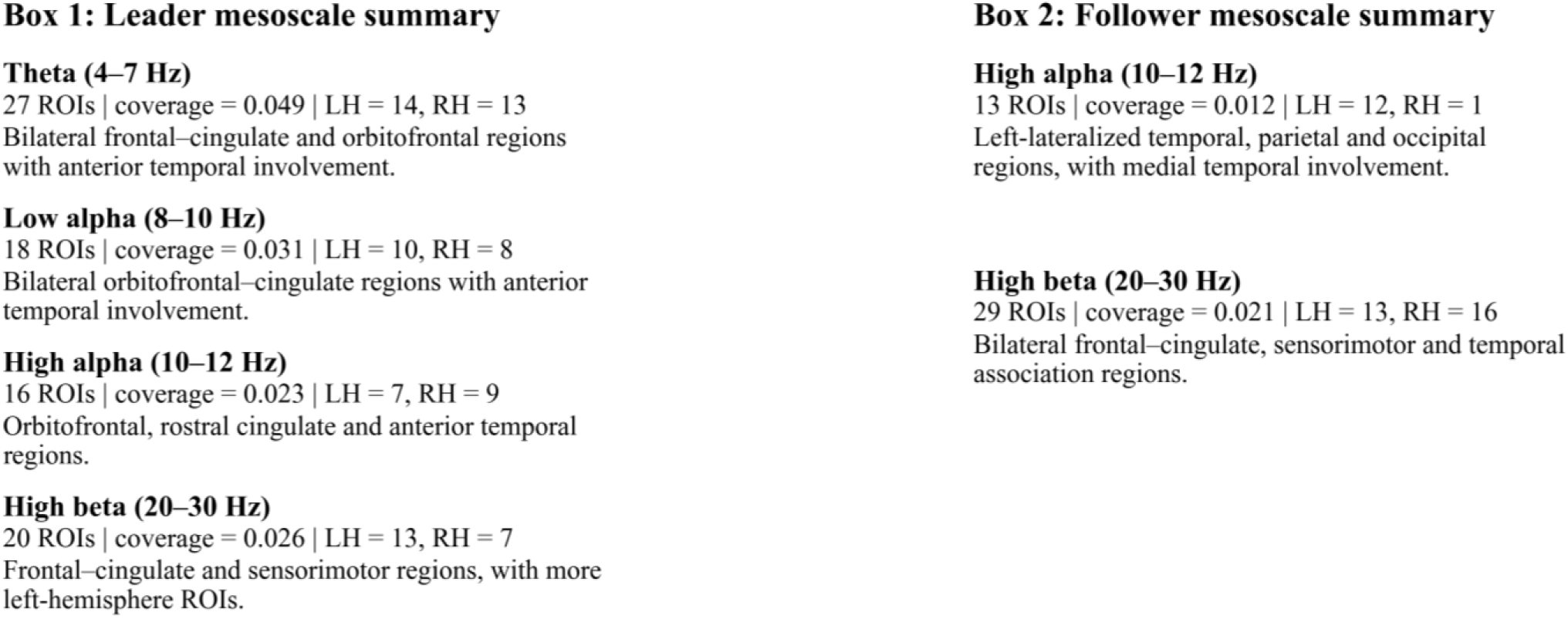
Role-specific time-resolved initiation–response–feedback. Figure 6a shows Leader time-resolved power estimates for theta (4–7 Hz) and low alpha (8–10 Hz) in the top row, and high alpha (10–12 Hz) and high beta (20–30 Hz) in the middle row. Figure 6b shows Follower estimates for high alpha (10–12 Hz) and high beta (20–30 Hz) in the bottom row. Blue traces indicate *Mirroring*; red traces indicate *Matching*. Coloured bands denote ±SEM across participants. Grey shading marks significant spatiotemporal Role or Role × Condition clusters identified by permutation testing; the cluster-level p value and contrast are specified in each panel. These clusters do not test the separation of the two condition traces within a role. The vertical black line marks the response-period onset (0 ms). The display spans −800 to +800 ms; the Follower high-beta cluster continues to +856 ms. Box 1 summarizes the four displayed Leader clusters by frequency band, reporting ROI count, spatiotemporal coverage, hemispheric distribution, and dominant anatomical pattern. Box 2 provides the corresponding summary for the two displayed Follower-related clusters. Table 6 and Supplementary Table S5a also report two significant Leader Role clusters not shown here: high alpha after onset (15–775 ms) and low beta spanning onset (−322 to +456 ms). Cluster ROI memberships are listed in Supplementary Table S5a.

**Table 6.** Spatiotemporal Cluster Statistics (Role-based Phasic Effects)

| **Contrast** | **Direction of Effect** | **Frequency Band** | **Time Window (ms)** | **Cluster p (FWER)** | **Mean dz** | **Peak \|dz\|** | **Dominant Neural Regions (compressed from ROI lists)** |
|---|---|---|---|---|---|---|---|
| Role | Leader > Follower (before onset) | Theta (4–7 Hz) | −763 → −275 | 0.0148 | 0.51 | 0.89 | Frontal midline (rostral anterior cingulate), premotor, orbitofrontal |
| Role | Leader > Follower (before onset) | Low Alpha (8–10 Hz) | −723 → −268 | 0.0122 | 0.49 | 0.72 | Orbitofrontal cortex, rostral anterior cingulate, temporal pole |
| Role | Leader > Follower (before onset) | High Alpha (10–12 Hz) | −718 → −277 | 0.0402 | 0.50 | 0.72 | Orbitofrontal cortex, rostral anterior cingulate, temporal pole |
| Role | Leader > Follower (after onset) | High Alpha (10–12 Hz) | 15 → 775 | 0.0060 | 0.47 | 0.71 | Lateral occipital, lingual, fusiform, middle temporal, posterior cingulate |
| Role | Leader > Follower (spanning onset) | Low Beta (13–20 Hz) | −322 → 456 | 0.0044 | 0.48 | 0.68 | Precuneus, superior parietal, lateral occipital, posterior cingulate, postcentral |
| Role | Leader > Follower (spanning onset) | High Beta (20–30 Hz) | −459 → 208 | 0.0262 | 0.46 | 0.86 | Dorsolateral frontal, anterior cingulate, precentral/postcentral |
| Role | Follower > Leader<sup>†</sup> (after onset) | High Beta (20–30 Hz) | 422 → 856 | 0.0482 | −0.46 | 0.74 | Middle temporal, fusiform, postcentral, anterior cingulate, frontal pole |
| Interaction | (Follower: Match−Mirror) > (Leader: Match−Mirror) (after onset) | High Alpha (10–12 Hz) | 398 → 726 | 0.0292 | 0.51 | 0.90 | Temporal pole, parahippocampal, precuneus, isthmus cingulate, lateral occipital |

***Note.*** Cluster-based permutation statistics were computed on epochs aligned to response-period onset (−1.0 to +1.0 s; 5,000 permutations). Reported p-values are cluster-level familywise error–corrected using the maximum cluster statistic across permutations within each band × contrast map (α = 0.05); no correction across maps was applied. Mean d<sup>z</sup> reflects the cluster-averaged standardized effect size, and Peak \|d<sup>z</sup>\| denotes the maximum local standardized effect within each cluster. Time windows indicate the contiguous temporal extent of each significant cluster. <sup>†</sup>The later high-beta cluster is consistent with a post-movement beta rebound following action completion.

Around Follower responses, we found a high-alpha (10–12 Hz) Role × Condition interaction after response-period onset (398–726 ms, *p* = 0.0292, mean *dz* ≈ 0.51; peak |*dz*| ≈ 0.90): the *Matching*–*Mirroring* difference was greater during Follower than Leader response periods. This cluster involved the temporal pole, parahippocampal cortex, precuneus, isthmus cingulate, and lateral occipital cortex. This cluster is consistent with inhibitory gating during the Follower response period (Jensen & Mazaheri, 2010; Klimesch, 2012) (see Figure 6b). A smaller high-beta (20–30 Hz) effect after response-period onset also favoured Follower over Leader response periods (422–856 ms, *p* = 0.0482, mean *dz* ≈ −0.46). This neural activity is consistent with a role-specific rebound following action completion (Pfurtscheller & Lopes da Silva, 1999). RT-controlled, across-map, and dyad-level sensitivity analyses are reported in Supplementary Sections S6.3–S6.4 and S6.6 and Tables S7–S8.

These phasic relationships reveal a role-asymmetric sequence, from anticipatory activity to activity following response-period onset. Around Leader responses, pre-response theta–alpha and peri-response beta activity may reflect anticipatory preparation and action initiation, with cue-related processing also contributing. For Follower responses, the high-alpha *Matching*–*Mirroring* contrast after Follower response-period onset was greater than for Leader response periods, consistent with condition-dependent inhibitory gating. A later high-beta effect may reflect post-action rebound. Together, these findings delineate the phasic signatures of the initiation–response–feedback cycle in leader–follower coordination. Next, we examine whether related role-specific structure is recoverable at the level of information geometry.

### 2.5 Inter-brain Information Decomposition (ΦID): Role-Specific Information Geometry

#### 2.5.1 Overview and survivor identification

To characterize the informational architecture of inter-brain coordination, Integrated Information Decomposition (ΦID) (Chidichimo et al., 2026; Luppi et al., 2022) was applied to paired ROI time series across both brains. ΦID decomposes joint predictive information from past dual-brain activity to future dual-brain activity into 16 atoms. Of these, we examined four atoms: *synergy* (sts), information available only from joint states in both past and future; *Unique-X* (xtx), information unique to the Leader across past and future; *Unique-Y* (yty), information unique to the Follower across past and future; and *redundancy* (rtr), information shared across both partners’ past and future. A linear-Gaussian model with the Minimum Mutual Information (MMI) function was fitted to homologous (Q1–Q4, Q2–Q3) and cross-hemispheric (Q1–Q3, Q2–Q4) ROI pairs spanning the neural geometry of the tetradic field (see Figure 1 for details).

In total, four contrasts were examined across ROI-pair × frequency-band × atom × geometric maps: *Matching* minus *Mirroring* averaged across response periods; Leader period minus Follower period within *Mirroring*; Leader period minus Follower period within *Matching*; and Leader period minus Follower period averaged across conditions. A separate response-period × condition interaction tested whether the *Matching*–*Mirroring* difference varied between Leader and Follower response periods. A three-tier control framework was applied: 5,000-permutation temporal cluster inference within each map, supplemented with trial-shuffle and time-shift surrogate tests. These constraints reduced the initial candidate set to a specific subset of inter-brain dependencies.

Within the four-contrast survivor set, the four selected informational atoms were represented (yty: n = 2; xtx: n = 2; rtr: n = 10; sts: n = 2), spanning both geometric axes (cross-hemispheric: n = 7; homologous: n = 9) and all six frequency bands. Episodes were brief (median ≈ 120 ms) and occurred before and after response-period onset. The separate response-period × condition interaction yielded three additional targets (rtr: n = 2; xtx: n = 1), spanning cross-hemispheric (n = 2) and homologous (n = 1) geometries. Summary statistics and spatial distributions are reported in Table 7.

**Table 7.** Integrated Information Decomposition (ΦID) Tier-A & B Survivor Summary by Atom, Functional Role, Frequency Band, Tetradic Geometry, and Neural Regions.

| **ΦID atom** | **Functional role** | **Survivors (n)** | **Tier A** | **Tier B** | **Frequency Band(s)** | **Tetradic geometry** | **Neural regions** |
|---|---|---|---|---|---|---|---|
| **Four condition and response-period contrasts** |  |  |  |  |  |  |  |
| Synergy (sts) | Emergent dyadic information | 2 | 1 | 1 | Theta | Homologous (2) | Rostral anterior cingulate |
| Unique-X (xtx) | Leader-specific predictive information | 2 | 1 | 1 | Theta, low beta | Cross-hemispheric (1); homologous (1) | Banks of the superior temporal sulcus; superior temporal cortex |
| Unique-Y (yty) | Follower-specific predictive information | 2 | 1 | 1 | Low alpha, high beta | Cross-hemispheric (1); homologous (1) | Insula; superior frontal cortex |
| Redundancy (rtr) | Shared predictive information | 10 | 7 | 3 | Theta, low alpha, high alpha, low beta, gamma | Cross-hemispheric (5); homologous (5) | Postcentral, medial orbitofrontal, isthmus cingulate, paracentral, lateral occipital, supramarginal, temporal pole, entorhinal, pars triangularis |
| **ΦID atom** | **Functional role** | **Survivors (n)** | **Tier A** | **Tier B** | **Frequency Band(s)** | **Tetradic geometry** | **Neural regions** |
| **Separate response-period × condition interaction** |  |  |  |  |  |  |  |
| Unique-X (xtx) | Leader-specific predictive information | 1 | 0 | 1 | Gamma | Homologous (1) | Frontal pole |
| Redundancy (rtr) | Shared predictive information | 2 | 1 | 1 | Theta, low alpha | Cross-hemispheric (2) | Caudal middle frontal; inferior temporal cortex |

***Note.*** Tetradic geometry counts indicate the number of survivors in homologous (Q1–Q4, Q2–Q3) or cross-hemispheric (Q1–Q3, Q2–Q4) pairings. For Unique-X, one pairing in each part of the table was identified in both geometries before surrogate controls; only the geometry shown reached Tier A or B. Tier A denotes targets meeting trial-shuffle recurrence in at least four dyads and time-shift recurrence in at least three dyads. Tier B denotes targets meeting the trial-shuffle criterion but not both Tier A criteria. Target-level details, including exact time windows relative to response-period onset, are reported in Supplementary Table S6a.

#### 2.5.2 Synergy (sts): Emergent dyadic information

The synergistic component (sts; n = 2), reflecting information that arises only when both partners’ pasts and futures are considered together, was found exclusively in the homologous geometry (see Figure 7a; Table 7). This pattern indicates that synergistic inter-brain information may depend on the preservation of hemispheric and self–other separation, with emergent information arising from interactions across distinct components of the dyadic system. Both surviving targets involved the right rostral anterior cingulate cortex, a region associated with cognitive–affective integration (Tang et al., 2019), and showed greater Follower- than Leader-period synergy before response-period onset: one within *Mirroring* (−950 to −813 ms) and one in the Leader–Follower comparison averaged over *Mirroring* and *Matching* (−951 to −774 ms).

**Figure 7a–d.**
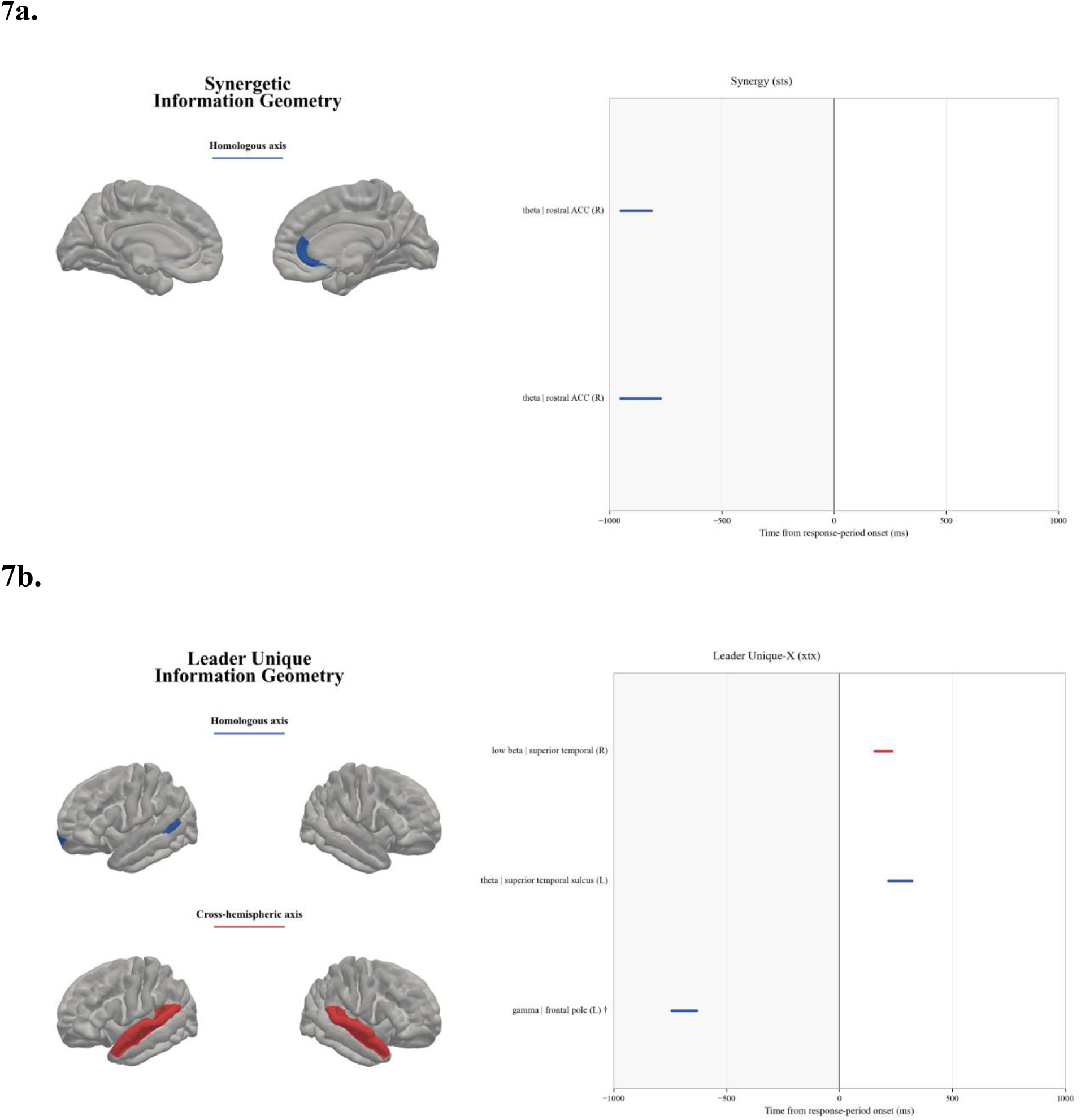
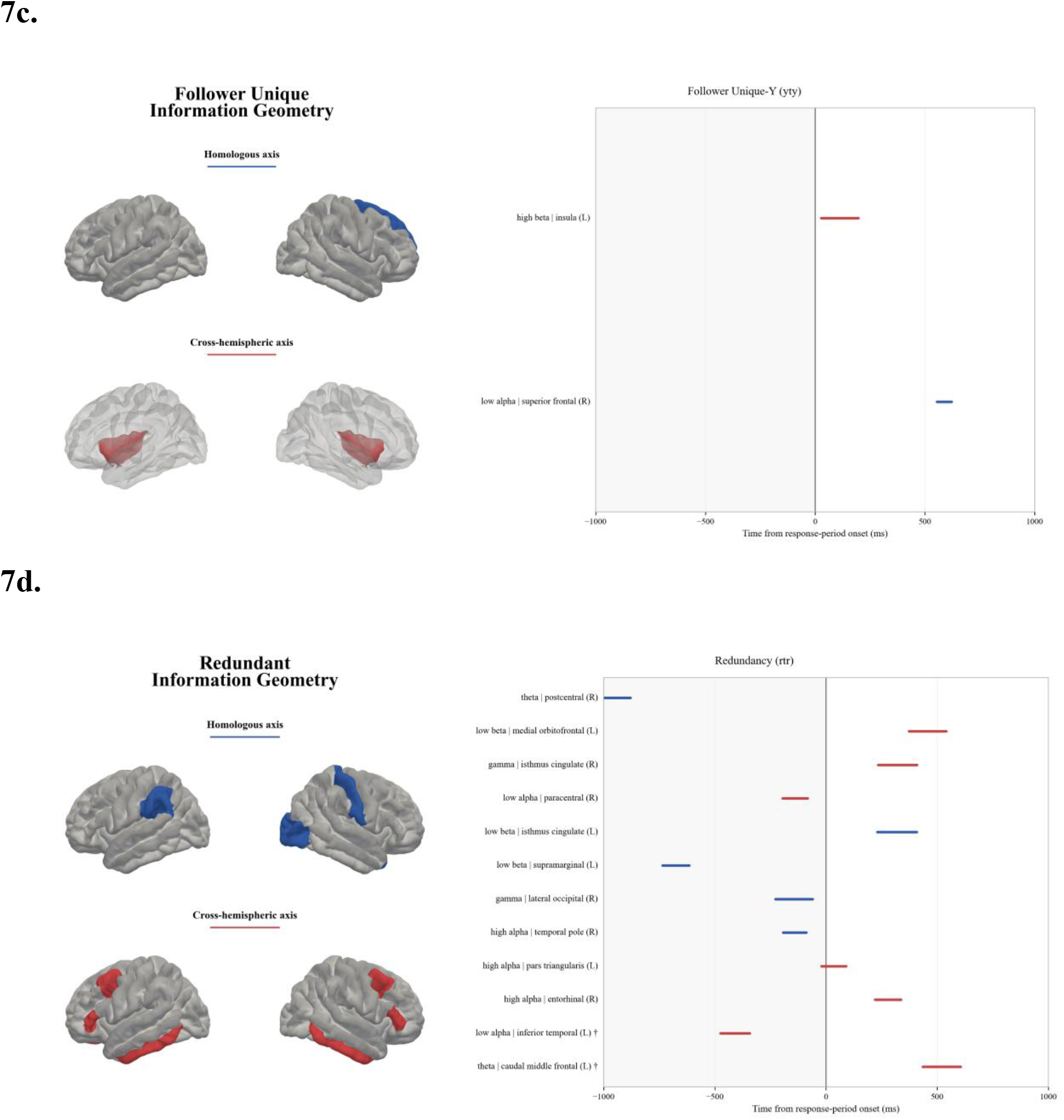
ΦID survivor windows across time, frequency, and tetradic geometry. Time-resolved survivor windows are shown across four ΦID atoms: Synergy (sts; Figure 7a), Leader Unique-X (xtx; Figure 7b), Follower Unique-Y (yty; Figure 7c), and Redundancy (rtr; Figure 7d). The left panel displays cortical maps of regions participating in surviving ΦID targets. Maps are shown separately for homologous and cross-hemispheric axes. Geometric categories are indicated by both panel position and colour-coded headings that match the raster colour scheme; blue denotes homologous and red denotes cross-hemispheric pairings. The right panel presents the corresponding time-resolved survivor windows for each atom, shown relative to response-period onset (0 ms). Horizontal segments indicate the intervals of Tier A or Tier B targets for each ΦID atom, frequency band, ROI pairing, and geometry. The dagger ^(†)^ identifies targets from the separate response-period × condition interaction. Contrast directions, exact time windows, and surrogate recurrence counts are reported in Supplementary Table S6a.

#### 2.5.3 Unique-X and Unique-Y: Role-specific predictive information

Unique-X survivors (xtx; n = 2), reflecting predictive information unique to the Leader across past and future, were distributed across cross-hemispheric (n = 1) and homologous (n = 1) geometries and localized to the right superior temporal cortex and left bank of the superior temporal sulcus (see Figure 7b; Table 7). In the Leader–Follower comparison averaged over *Mirroring* and *Matching*, the homologous theta target was greater during Leader than Follower response periods (217–321 ms), whereas the cross-hemispheric low-beta target was greater during Follower than Leader response periods (156–233 ms).

Moreover, we found a separate Unique-X interaction target localized to the left frontal pole in the homologous geometry (gamma; −743 to −631 ms) (see Figure 7b; Table 7). Here, the *Matching*–*Mirroring* difference was greater during Follower than Leader response periods.

Unique-Y survivors (yty; n = 2), reflecting predictive information unique to the Follower across past and future, were distributed across cross-hemispheric (n = 1) and homologous (n = 1) geometries and localized to the left insula and right superior frontal cortex (see Figure 7c; Table 7). The cross-hemispheric high-beta insula target was greater during *Matching* than *Mirroring* (27–197 ms); the homologous low-alpha superior-frontal target was greater during Follower than Leader response periods in the comparison averaged over both conditions (555–622 ms). Both effects occurred after response-period onset.

#### 2.5.4 Redundancy (rtr): Shared predictive scaffold

The redundant component (rtr; n = 10) reflects information shared across both partners’ pasts and futures, indexing the mutual predictive structure underpinning coordination within the tetradic reference frame. Survivors were detected in homologous (n = 5) and cross-hemispheric (n = 5) geometries, involving sensorimotor, frontal, temporal, parietal, cingulate, and occipital regions (see Figure 7d; Table 7). The two condition targets showed greater redundancy during *Matching* than *Mirroring*. Response-period contrasts varied by condition: one target within *Mirroring* was greater during the Follower period, while two within *Matching* were greater during the Leader period. In the comparison averaged across conditions, four targets were greater during the Follower period and one during the Leader period. These effects occurred before and after response-period onset.

The separate response-period × condition interaction yielded two cross-hemispheric redundancy targets. The *Matching*–*Mirroring* difference was greater during the Follower than Leader period in the left caudal middle frontal cortex (theta; 435–606 ms), but greater during the Leader than Follower period in the left inferior temporal cortex (low alpha; −475 to −343 ms).

### 2.6 Cross-Method Synthesis

Across analytic levels, a consistent condition structure organizes the data: *Matching* (controlled, inhibited coordination) and *Mirroring* (automatic, resonant coordination) represent complementary poles of an attention–inhibition gating mechanism that governs dyadic sensorimotor coordination from behaviour through to inter-brain information structure.

At the behavioural level, this structure is expressed as a role-differentiated RT asymmetry: Followers are strongly sensitive to the *Mirroring*/*Matching* condition, while Leaders are comparatively insensitive. At the tonic neural level, *Matching* imposes a sustained energetic demand concentrated in frontoparietal and salience networks, with no Role × Condition effects surviving FDR correction. Phasically, *Matching* showed greater source-level power than *Mirroring* within significant condition clusters aligned to response-period onset in the alpha, beta, and gamma ranges. Role-specific clusters further delineated a perception–action–feedback sequence: anticipatory theta–alpha activity and peri-response beta activity around Leader responses, followed by high-alpha and high-beta effects after Follower response-period onset. Viewed as a dyadic sequence, these clusters suggest a Leader-driven anticipatory cascade extending into action execution and a Follower-mediated phase of post-response gating and rebound that may prepare the system for subsequent interaction.

At the level of inter-brain information, ΦID decomposition reveals role-specific atomic contributions: Leader-unique predictive information in superior temporal regions and Follower-unique predictive information in insular and superior frontal regions. The expression of these contributions differs across the two modes of coordination. *Matching* showed greater redundancy than *Mirroring* in two targets and greater Follower-unique information in the insula; within *Matching*, two other redundancy targets were greater during the Leader period. Within *Mirroring*, synergy and one redundancy target were greater during the Follower period. The shared predictive scaffold and emergent dyadic information therefore show different surviving role-period patterns under *Matching* and *Mirroring*, as opposed to a uniform effect across the interaction space.

Together, these findings characterize the baseline architecture of leader–follower coordination as a multiscale system in which a global condition structure is evident at the aggregate level, alongside asymmetric, role-specific organization in time-resolved neural dynamics and inter-brain information decomposition. This convergence provides the empirical grounding for the tetradic framework and establishes normative reference parameters for the planned clinical extension of this paradigm (ClinicalTrials.gov ID: NCT06978803).

## 3. Methods

### 3.1 Participants

Two datasets were acquired.

The first is a sample of 32 adults (16 mixed-sex dyads; 12 men, 20 women; mean age 20.8 years, range 18–40; 30 right-handed) recruited from within the Department of Psychology, University of Manitoba. All subjects were enrolled in Introduction to Psychology at the University of Manitoba and received research participation credits. Inclusion required normal or corrected-to-normal vision and that subjects were not made aware of the experiment’s purpose. All participants provided written informed consent under protocols approved by the Psychology/Sociology Research Board (Protocol HS19206 (P2015:153)).

The second is another healthy control sample of 32 adults (16 mixed-sex dyads; 12 men, 19 women, 1 non-binary; mean age 27.2 years, range 18–57; 31 right-handed) who took part additionally in the hyperscanning EEG experiment at the Douglas Mental Health University Institute. Participants were fluent in English or French; dyads were matched for language preference. Inclusion required age 18–60 and the absence of any neurological or psychiatric diagnosis by self-report. All participants provided written informed consent under protocols approved by the Douglas Mental Health University Institute Research Ethics Board (Protocol 2022-595).

### 3.2 Experimental Design and Task

Dyadic sensorimotor coordination was assessed using a leader–follower stimulus–response task implemented during simultaneous dual-brain EEG hyperscanning (for experimental paradigm see Figure 8). The task employed a 2 × 2 within-subject factorial design crossing Role (Leader/Follower) and Condition (*Mirroring*/*Matching*). Participants were seated face-to-face across an 80-cm square table, their physical midlines equidistant, defining the orthogonal intersection of the self–other and left–right axes.

**Figure 8.**
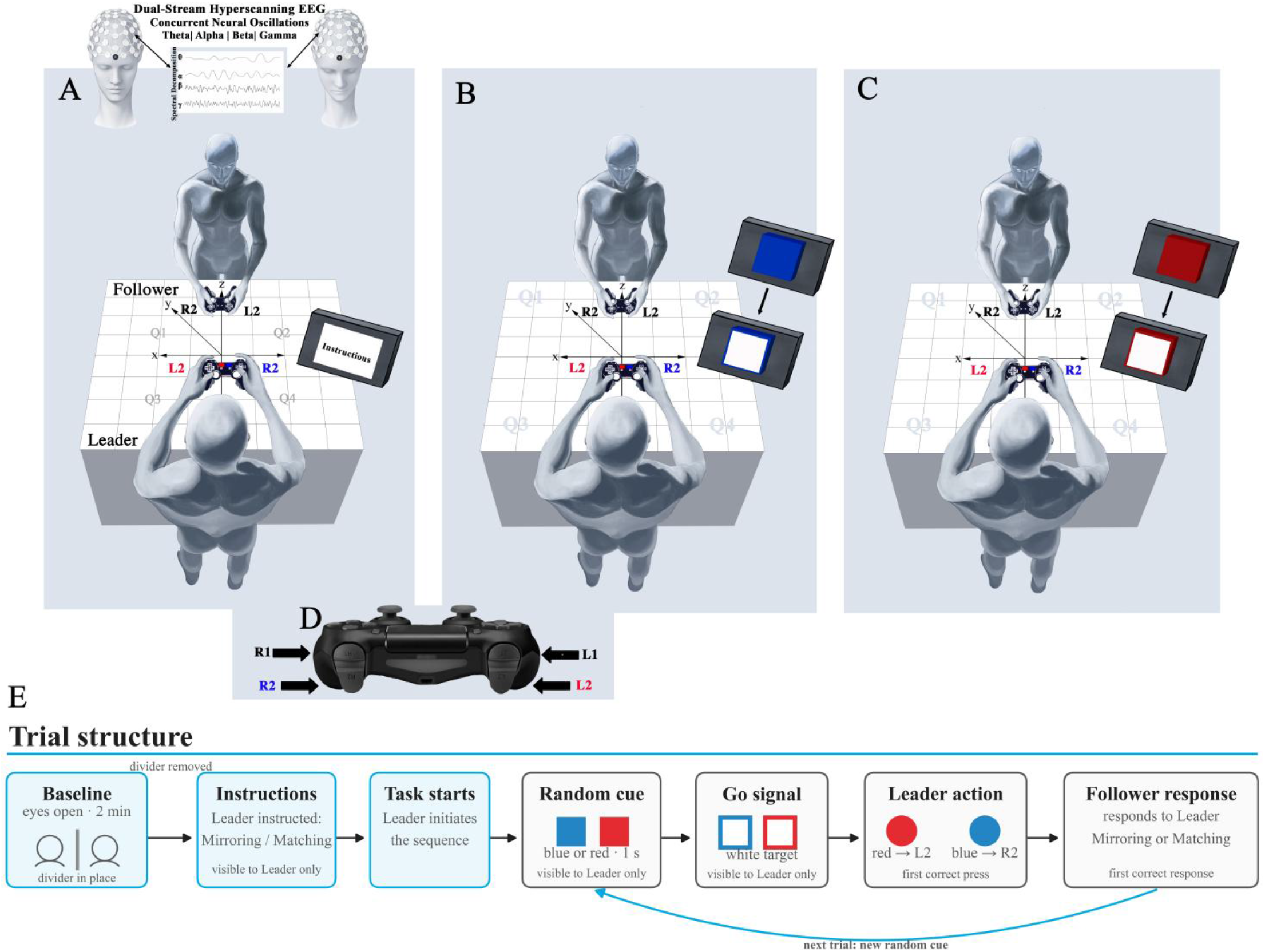
Leader–follower sensorimotor coordination task with EEG hyperscanning. A pair of 128-electrode geodesic EEG sensor nets is fitted to a dyad for simultaneous dual-stream neural recording (A). The pair is positioned face-to-face within a shared x–y–z spatial reference frame (A). This multiscale organization is subdivided into left and right, as well as self and other. This fourfold symmetry forms the tetradic field that underpins the geometries of interaction. To map the changing configural dynamics within this field, a 2 × 2 within-subjects design is employed, crossing Role (Leader/Follower) and Condition (*Mirroring*/*Matching*). (E) The experimental sequence begins with a two-minute eyes-open baseline, followed by block-specific instructions and task initiation. The task comprises four blocks of 120 trials; Leader and Follower roles switch after Block 2, and conditions follow the sequence *Mirroring*–*Matching*–*Matching*–*Mirroring* (Table 8). At the beginning of each block, instructions are displayed on the monitor, read aloud by the Leader, and verbally affirmed by the Follower. Each trial begins with a randomly selected blue or red cue presented for 1 s, followed by a white target bordered by the original cue colour, which serves as the go signal (B, C; arrows indicate stimulus progression). At go-signal onset, the Leader responds using a lateralized controller trigger—L2 for a red cue and R2 for a blue cue (D). Leader response time is measured from go-signal onset to the first correct trigger press; following an incorrect press, timing continues until the correct response is executed. The Follower then produces the response specified by the block condition. In *Mirroring*, the Follower presses the trigger opposite to the Leader’s response (R2 following L2 or L2 following R2), producing cross-hemibody coordination across the dyad (Q1–Q3 and Q2–Q4). In *Matching*, the Follower presses the same trigger as the Leader (R2 following R2 or L2 following L2), producing homologous hemibody coordination (Q1–Q4 and Q2–Q3). Because motor output is contralaterally controlled and the participants are positioned face-to-face, homologous trigger pairings occupy contralateral positions in shared space (opposite sides of the shared left–right field), whereas opposite-trigger pairings occupy ipsilateral positions (the same side of that field) (see Figure 2 for the geometric mapping). Follower response time is measured from the Leader’s first correct response to the Follower’s first correct mapped response; following an incorrect response, timing continues until the required configuration is achieved. The next trial then begins with a newly randomized cue. Adapted from Olarewaju (2021).

**Table 8.** Experimental Block Structure.

| **Block** | **P1 Role** | **P2 Role** | **Condition** | **Trials** |
|---|---|---|---|---|
| 1 | Leader | Follower | Mirroring | 120 |
| 2 | Leader | Follower | Matching | 120 |
| 3 | Follower | Leader | Matching | 120 |
| 4 | Follower | Leader | Mirroring | 120 |
| **Total** |  |  |  | **480** |

***Note.*** P1 = Participant 1; P2 = Participant 2. Role assignment (P1 vs. P2) was randomized at session start. Role order across blocks was fixed by this assignment; the condition sequence was *Mirroring–Matching–Matching–Mirroring*. Blocks 1–4 each comprised 120 trials (480 total).

Under *Mirroring*, Followers reproduced the Leader’s movement in visual mirror coordinates, such that the Leader’s right-hand action corresponded to the Follower’s left-hand response, and vice versa. Under *Matching*, Followers reproduced the Leader’s movement in anatomical coordinates, such that right-hand actions were matched with right-hand responses and left-hand actions with left-hand responses.

Both participants responded using L2/R2 triggers on PlayStation 5 controllers, logged by E-Prime 3.0 with millisecond precision and time-stamped against EEG event markers. Participants were instructed to respond as quickly and accurately as possible and to treat each trial as a joint action. Trial progression depends on the completion of a correct dyadic response, actively monitored by Leaders. Thus, recorded latencies capture the time required to achieve a valid joint configuration.

Each dyad completed four experimental blocks of 120 trials each (480 trials total). Role order was fixed by participant assignment (see Table 8): Participant 1 served as Leader in Blocks 1–2 and Follower in Blocks 3–4; Participant 2 assumed the complementary order. The condition sequence was Mirroring–Matching–Matching–Mirroring, so each participant completed one block of each condition in each role. A four-block practice phase (30 trials per block) was completed before EEG net application. A 2-minute resting-state baseline was recorded with participants separated by a midline divider immediately before the experimental task. Note that role order was not counterbalanced across dyads; the implications of this design constraint are discussed in Section 4.

### 3.3 EEG Acquisition

Continuous EEG was acquired simultaneously from both dyad members using a dual-stream hyperscanning configuration (Magstim EGI Geodesic EEG System 400; two independent Net Amps 400 amplifiers). Neural signals were recorded via 128-channel HydroCel Geodesic Sensor Nets (saline-based) at 1,000 Hz. Electrode impedances were maintained below 50 kΩ. Temporal alignment between streams was achieved using the EGI Hyperscanning clock synchronization device and a shared event-marker infrastructure, yielding sub-millisecond inter-stream synchrony.

### 3.4 Behavioural Statistical Analysis

#### 3.4.1 Aggregate analysis

Condition-wise median RTs (NRT = 32) were analyzed using a 2 × 2 repeated-measures ANOVA (within-subject factors: Role, Condition) with Bonferroni-corrected post-hoc comparisons. The same procedure was applied to a combined behavioural cohort (NRT = 64) formed by pooling the current hyperscanning sample with an independent dataset collected under equivalent procedures (Olarewaju, 2021) to assess cross-cohort replicability (see Supplementary section S1 for procedural-equivalence documentation). Primary-cohort RT contrasts were also tested with dyads as independent units (Supplementary Section S1.3).

#### 3.4.2 Trial-level linear mixed-effects model

Trial-level log-transformed RTs were analyzed using a linear mixed-effects model (LMM) with fixed effects for Role, Condition, and their interaction (sum-coded: Leader = −0.5, Follower = +0.5; *Matching* = −0.5, *Mirroring* = +0.5). Random effects included participant intercepts, role and condition slopes were supported by convergence, and a dyad-level variance component; dyad fixed effects replaced dyadic random intercepts when the latter were unstable. The Role × Condition interaction was tested by likelihood-ratio comparison of nested models under identical random-effects structures. Estimates were back-transformed from log to millisecond scale via exponentiation; approximate Nakagawa R^2^ (marginal and conditional) was reported.

#### 3.4.3 Complementary analyses

Within-participant Welch-type contrasts were computed per participant to characterize individual-level consistency of the *Mirroring* advantage. Bayesian LMMs (Bambi / PyMC backend; see Supplementary section S2 for prior specification and MCMC parameters) provided complementary uncertainty estimates; practical equivalence was evaluated using a region of practical equivalence (ROPE) of ±5% on the multiplicative RT scale.

### 3.5 EEG Preprocessing

All preprocessing was performed in MNE-Python (v1.11.0) within the HyPyP (v0.5.0) hyperscanning framework (Ayrolles et al., 2021). Continuous data were notch-filtered at 60 Hz (and 120 Hz harmonic) and band-pass filtered 1–100 Hz (zero-phase FIR, Hamming window; full filter parameters in Supplementary section S3). Data were re-referenced offline to the common average. Channels with flat signals, intermittent dropouts, or high-amplitude noise were identified visually and excluded from ICA fitting (retained for later interpolation).

Filtered, re-referenced data were segmented into epochs (−1.0 to +1.0 s) aligned to task markers initiating the Leader and Follower response periods. For each dyad member, separate epochs were aligned to the onset of the corresponding Leader or Follower response period. Artifact correction used joint ICA (Extended Infomax; n = 20 components; HyPyP joint decomposition across both dyad members), followed by AutoReject (union strategy; 50 μV threshold) for residual noisy channels and epochs. Ocular artifact invariance across conditions was verified via GEE analysis of trial-level blink rates (see Supplementary section S3). Six frequency bands were retained for source-level analyses: theta (4–7 Hz), low-α (8–10 Hz), high-α (10–12 Hz), low-β (13–20 Hz), high-β (20–30 Hz), and Gamma-γ (30–45 Hz).

### 3.6 EEG Source Reconstruction and Parcellation

Source reconstruction was performed in MNE-Python using the FreeSurfer *fsaverage* template. A cortical source space was defined on an octahedral (*oct6*) mesh (∼4,098 dipoles per hemisphere). A three-layer boundary element model (BEM: brain, skull, scalp) served as the forward model; inverse solutions were computed using dynamic statistical parametric mapping (dSPM) with a grand-average noise covariance matrix estimated from the pre-stimulus baseline (details in Supplementary section S4). Source estimates were parcellated into 68 cortical regions of interest (ROIs) using the Desikan–Killiany atlas. For tonic analyses, band power was estimated from ROI time courses using Welch’s method. For phasic and information theoretic analyses, time-resolved power was computed via Morlet wavelet convolution. The ΦID analysis used a separately retained set of 12,218 paired epochs and source estimates generated with a rebuilt shared inverse operator. Its epoch membership and source inputs were therefore not identical to those used for the tonic and phasic analyses; the ΦID frequency bins are specified in Section 3.9.

A subset of NEEG = 30 participants (15 dyads) was included in source-level analyses following exclusion of two participants from one dyad for technical recording failures. Source-level analyses were restricted to this subset throughout; behavioural analyses used the full NRT = 32 cohort.

### 3.7 Tonic EEG: Mixed-Effects ROI × Band Models

Tonic (trial-averaged) oscillatory power across the 408 ROI × frequency-band features (68 ROIs × 6 bands) was modelled using trial-level mixed-effects models with Role, Condition, and Role × Condition as fixed effects and a within-participant z-scored RT covariate to partial out speed–power correlations. Random effects included participant intercepts. Multiple-comparison control used Benjamini–Hochberg FDR (q ≤ .05) applied separately to each of the four effects— Condition, Role, Role × Condition, and RT—across 408 features. Spatial overlap across roles was then quantified using the Jaccard index computed on these independently FDR-corrected survivor sets. RT-adjusted tonic contrasts were also tested at the dyad level (Supplementary Section S5.1).

### 3.8 Phasic EEG: Spatiotemporal Cluster Permutation Testing

Phasic, time-resolved source power was analyzed using spatiotemporal cluster-based permutation tests (MNE-Python; 5,000 permutations; cluster-level FWER α = .05 within each band × contrast map). In the primary analysis, no correction across the 18 maps was applied. Epochs spanned −1.0 to +1.0 s around response-period onset; spatial adjacency was derived from the Desikan–Killiany parcellation centroid distances. Three orthogonal contrasts were tested within the 2 × 2 factorial design: Condition (*Matching* − *Mirroring*), Role (Leader − Follower), and Role × Condition. The Role × Condition interaction was defined as [(Follower: Matching − Mirroring) − (Leader: Matching − Mirroring)] / 2; positive values indicate a larger Matching–Mirroring difference during Follower response periods, and negative values indicate a larger difference during Leader response periods. Effect sizes (Cohen’s |dᶻ|) were computed at each significant cluster vertex. A complementary space-frequency (ROI × band) cluster analysis on time-averaged power confirmed Condition effects. EMG-prone ROI exclusion and frequency-specific SNR checks are described in Supplementary Sections S6.1–S6.2. In a separate RT-controlled sensitivity analysis, trial-level log1p power was adjusted for within-participant standardized RT at each ROI × band × time point before the cluster tests were repeated (Supplementary Section S6.3). An additional conservative check applied Bonferroni correction across the 18 band × contrast maps, treating the primary and RT-controlled analyses as separate families (Supplementary Section S6.4).

To test whether the response-period-aligned Condition effects contained temporal variation beyond a sustained difference, we subtracted each participant’s full-epoch mean *Matching*– *Mirroring* contrast at each ROI and band, then repeated the six Condition cluster tests using the original adjacency and 5,000 permutations (Supplementary Section S6.5). Cluster inference was also repeated with dyads as sign-flip units (Supplementary Section S6.6).

### 3.9 Integrated Information Decomposition (ΦID)

Inter-brain informational structure was characterized using Integrated Information Decomposition (ΦID; Luppi et al., 2022), implemented in the phyid framework with a Gaussian/correlation-based estimator and minimum mutual information (MMI) redundancy. The use and interpretation of MMI redundancy are described in Supplementary Section S7.6. ΦID decomposes the mutual information between the dyad’s joint past and future states, defined here as (S = (Xt−τ, Yt−τ); T = (Xt+τ, Yt+τ)), into a 16-atom information lattice. Here, τ = 1 sample at 1000 Hz, corresponding to a 1 ms offset on either side of each local time point and a 2 ms past–future separation, was used to define a local transition in time-resolved source-level band-power envelope dynamics (see Supplementary Sections S7.2–S7.5 for tensor construction, τ-specific surrogate controls, history/τ embedding selection, selection-adjusted surrogate controls, signal-timescale diagnostics, and chronological task-half comparisons).

This analysis focused on four hypothesis-driven persistence atoms corresponding to the tetradic information geometries that constrain the dyadic system and are experimentally expressed through the Leader/Follower × *Mirroring*/*Matching* task structure: (1) synergy-preserving information (sts), (2) leader-unique-preserving information (xtx), (3) follower-unique-preserving information (yty), and (4) redundancy-preserving information (rtr). The remaining twelve off-diagonal atoms, which describe transformations between informational modes across the past-future transition, were not included in the inferential model (see Supplementary Section S8.2 for the rationale for restricting inference to diagonal persistence atoms).

For each epoch, X was assigned to the task Leader and Y to the Follower: event suffix 1 assigned X = P1 and Y = P2, whereas suffix 2 assigned X = P2 and Y = P1. The Leader/Follower event prefix identified the sampled response period. ΦID was computed separately for homologous pairings (Q1–Q4, Q2–Q3) and cross-hemispheric pairings (Q1–Q3, Q2–Q4), corresponding to the principal inter-brain geometries of the tetradic face-to-face configuration. For ΦID, the included integer-Hz frequencies were theta (4–7 Hz), low-α (8–9 Hz), high-α (10–12 Hz), low-β (13–19 Hz), high-β (20–29 Hz), and gamma (30–45 Hz).

ΦID time series were tested using per-ROI temporal cluster-based permutation tests (5,000 permutations; cluster-forming α = .05; FWER controlled across time within each ROI × band × atom × geometry map). Four contrasts were tested: Matching − Mirroring; Leader − Follower within Mirroring; Leader − Follower within Matching; and Leader − Follower averaged across conditions. A separate response-period × condition interaction tested whether the Matching– Mirroring difference varied between Leader and Follower response periods (see Supplementary Section S7.1 for definitions).

A three-stage control framework was applied: primary permutation inference, a trial-shuffle null (disrupting trial correspondence while preserving within-trial structure), and a time-shift null (disrupting temporal alignment while preserving within-participant autocorrelation). Geometry overlap was identified before surrogate controls; homologous and cross-hemispheric targets were then evaluated separately, each receiving its own robustness tier. Time-shift tests were conducted for targets with a group trial-shuffle p ≤ .10. Feature-level robustness tiers were then defined by cross-dyad recurrence under the surrogate controls. Tier A features had trial-shuffle support in at least four dyads and time-shift support in at least three dyads. Tier B features had trial-shuffle support in at least four dyads without meeting both Tier A requirements. Tier A and Tier B together constituted the strict survivor set.

Additional sensitivity analyses examined lag, signal timescale, history length, and differences between the early and late chronological task halves. The lag and history analyses included corresponding surrogate controls and recurrence tiers. The task-half comparison examined unique-information targets across the concurrent changes in participant identity, chronological half, and condition order. The four main contrasts and response-period × condition interaction were evaluated separately (Supplementary Sections S7.3–S7.5).

To assess stability across interacting pairs, an across-dyad recurrence analysis was applied to the strict survivor set by quantifying the number of dyads exhibiting each effect under surrogate control procedures. Main-text results summarize the strict survivor architecture, whereas the primary target summaries and recurrence counts are provided in Supplementary Table S6a; the sensitivity results are reported in Supplementary Tables S6b–S6e.

### 3.10 Software and Reproducibility

All analyses were implemented in Python 3.11.11 (Bayesian LMM: Python 3.10.18) on Windows 11 (Intel i9-14900K, 64 GB RAM). Key packages: MNE-Python v1.11.0 (EEG preprocessing and source reconstruction), HyPyP v0.5.0 (hyperscanning QC and joint ICA), Bambi v0.17.2 / PyMC v5.27.1 (Bayesian LMM), ArviZ v0.23.4 (convergence diagnostics), phyid (ΦID). Fixed random seeds were used; intermediate outputs (cleaned epochs, source tensors, contrast definitions, permutation results) were saved with full metadata. E-Prime 3.0 controlled stimulus presentation and response logging.

## 4. Discussion

### 4.1 Summary of Principal Findings

This study characterized the baseline multiscale architecture of leader–follower sensorimotor coordination across four analytic levels. First, in the primary cohort, *Mirroring* produced a robust reaction-time advantage for Followers but a smaller, non-significant advantage for Leaders. The Leader advantage reached significance when the independent behavioural cohort was added. Second, tonic oscillatory power revealed a widespread Condition effect, predominantly *Matching* > *Mirroring*; no Role × Condition feature survived FDR correction. Third, phasic time-resolved analyses revealed *Matching* > *Mirroring* Condition clusters in alpha, beta, and gamma bands across epochs aligned to response-period onset, alongside role-related activity consistent with anticipation around Leader responses and post-response gating and rebound around Follower responses. Fourth, ΦID identified a role-specific predictive architecture with targets before and after response-period onset: Leader-unique information in superior temporal regions, Follower-unique information in insular and superior frontal regions, sparse synergy in the right rostral anterior cingulate cortex, and a broader Redundancy scaffold across homologous and cross-hemispheric mappings involving sensorimotor, frontal, temporal, parietal, cingulate, and occipital regions.

Together, these findings address all four stated aims and converge on a coherent account: a Matching-dominant Condition pattern is evident in tonic power and significant phasic condition clusters, while role-specific structure emerges in time-resolved neural dynamics and inter-brain information decomposition.

### 4.2 Aim 1: Behavioural Asymmetry: Role-Dependent Attention–Inhibition Gating

The Follower *Mirroring* advantage reflects role-dependent modulation of a shared automatic imitation field within the dyadic system. In the *Mirroring* configuration, spatial correspondence between effectors across the face-to-face axis directly maps perception onto action, facilitating response selection and reducing latency. In the *Matching* configuration, this correspondence must be overridden, requiring inhibitory control and incurring a measurable response time cost. These effects are consistent with established accounts of Automatic Imitation (Heyes, 2011; Ito, 2023; Wilt et al., 2024) and related stimulus–response compatibility phenomena, including the Simon Effect (Campbell et al., 2023; Simon, 1969) and Joint Simon Effect (Dolk et al., 2014), which demonstrate that action selection is systematically shaped by the spatial alignment between observed and executed movements (Sebanz et al., 2006).

Within this framework, both Leaders and Followers are embedded in the same correspondence-driven field; however, role-specific task constraints modulate how these automatic tendencies are expressed. Followers, whose responses are contingent on the Leader’s action, operate directly within this interpersonal mapping and therefore exhibit a strong *Mirroring* advantage when correspondence is preserved. Leaders, although subject to the same underlying compatibility structure, respond to a tangential go-signal and are therefore less behaviourally constrained by the observed action at lower sample sizes. With increased power, a smaller but reliable *Mirroring* advantage emerges in Leaders, confirming that automatic imitation operates in both roles but is differentially expressed as a function of role-dependent control demands.

### 4.3 Aim 2: Tonic Neural Activity: A Role-Invariant Condition Polarity

The dominance of *Matching* > *Mirroring* across frontoparietal, salience, and default mode network features is consistent with *Matching* exerting greater sustained demand on executive control: suppressing automatic spatial correspondence requires continuous top-down inhibitory regulation, reflected in elevated time-averaged alpha, beta, and gamma power (Buckner et al., 2008; Cole et al., 2013; Lundqvist et al., 2024; Menon et al., 2022; Schimmelpfennig et al., 2023; Seeley et al., 2007; Ulloa, 2022; Vincent et al., 2007). This pattern exhibited strong role-invariance, with near-complete spatial overlap across roles for the dominant contrast.

These results indicate that both Leaders and Followers share the block-level energetic cost of sustained controlled versus automatic processing, even though their moment-to-moment selection demands differ. Tonic measures therefore primarily index the shared metabolic architecture of Condition: five of 408 features showed a Role main effect, but no Role × Condition interactions survived correction (Table 4). Time-resolved and information-theoretic analyses further characterize role-specific organization. Independent role-specific overlap analysis is detailed in Supplementary section S8.1.

### 4.4 Aim 3: Role-Specific Phasic Dynamics

The *Matching* > *Mirroring* condition pattern identified in tonic averages was also expressed across the **period-aligned** epoch. Tonic averaging captures the aggregate condition difference, while spatiotemporal cluster analysis identifies its anatomical and temporal extent across the **period-aligned** epoch.

Crucially, this condition-related pattern coexists with role-specific phasic dynamics that may reflect anticipatory preparation around Leader responses and post-response gating and rebound around Follower responses, in keeping with established distinctions between predictive motor preparation and post-response inhibitory regulation (Deco et al., 2008; Friston, 2010; Hari & Kujala, 2009; Ivry & Spencer, 2004; Jensen & Mazaheri, 2010; Miller & Cohen, 2001). Estimated cluster-mean |*d*ᶻ| values across the significant condition, Role, and Role × Condition clusters were approximately 0.46–0.51 (Tables 5 and 6). This functional interpretation remains qualified by the experimental design: Leader and Follower epochs are centered on successive response-period onsets within a trial, so the Role contrast cannot fully separate role from temporal context.

### 4.5 Aim 4: ΦID: Role-Specific Predictive Architecture

The ΦID decomposition provides an information-theoretic account of inter-brain role asymmetry. Unique-X, localized to superior temporal regions and, in a separate interaction, the left frontal pole, reflects Leader-specific predictive information expressed across both Leader and Follower response periods. These findings suggest distinct temporal and condition-dependent frontal contributions to the dyadic predictive architecture, consistent with executive-control (Miller & Cohen, 2001) and predictive coding (Friston, 2010) frameworks. Unique-Y, **l**ocalized to the left insula and right superior frontal cortex, reflects Follower-unique predictive information whose condition- and response-period-dependent expression is consistent with **t**he Follower’s adaptive updating of partner-derived signals and a prediction-error correction process that modulates ongoing coordination and action selection (Friston, 2010; Llinás & Roy, 2009; Schwartz, 2016; Wolpert et al., 1995). Synergy, localized to the right rostral anterior cingulate cortex, reflects emergent dyadic information within a region associated with cognitive–affective integration (Tang et al., 2019). The Redundancy component, distributed across homologous and cross-hemispheric sensorimotor, frontal, temporal, parietal, cingulate, and occipital regions, suggests that the shared predictive scaffold is expressed across a broader cortical architecture and changes with condition and response period.

### 4.6 Cross-Method Synthesis

Across analytic levels, a coherent multiscale organization emerges: the *Mirroring*– *Matching* distinction is expressed as a role-dependent behavioural RT difference and a Matching-dominant neural power pattern at tonic and condition-level phasic scales, while further role-specific structure appears in time-resolved dynamics and inter-brain information decomposition. This organization is consistent with an initiation–response–feedback cycle, with a Leader-driven anticipatory cascade preceding action and Follower-mediated post-response gating and rebound that may reorganize the dyadic state. Consistent with this structure, ΦID identifies a sparse predictive architecture spanning Unique-X, Unique-Y, Synergy, and Redundancy, with targets occurring both before and after response-period onset.

Together, these findings show that condition-level organization and role-specific dynamics coexist across analytic resolutions.

### 4.7 Limitations

The experimental design imposes a structured ordering of roles across blocks. This introduces a temporal asymmetry between role and block position (early vs late interaction). Chronological task-half comparisons of five strict unique-information targets, spanning the planned reversal of participant roles, found no early–late differences surviving Holm correction. Detailed methods and full results are provided in the Supplementary section S7.5 and Table S6b. Accordingly, the design does not fully dissociate role from temporal context. A fully counterbalanced within-subject role-swap design, dissociating role from interaction history, remains a priority for future study.

A second consideration concerns generalizability of the ΦID effects. The strict survivor set reflects the imposition of surrogate-recurrence criteria, yielding a sparse subset of temporally and statistically robust predictive dependencies. While this selectivity strengthens internal validity by isolating features that persist under perturbation, replication in an independent cohort is required to establish the stability of these effects across samples. Finally, the dimensionality of the ΦID analysis imposes practical constraints on exhaustive exploration of the full feature space.

### 4.8 Conclusion

We articulate and test a tetradic account of dyadic sensorimotor coordination, organized by a role-asymmetric attention–inhibition gate whose expression is scale-dependent. The *Mirroring/Matching* polarity establishes a robust behavioural anchor for interpersonal gating, embedded within role-specific neural signatures and inter-brain information geometry, which together provide a multiscale basis for clinical comparison in the registered Phase 2 extension of this study involving schizophrenia patient–healthy control dyads (ClinicalTrials.gov ID: NCT06978803).

This reproducible pipeline examines interpersonal coordination without collapsing its variability across scales or reducing it to isolated neural or behavioural effects (Bolis et al., 2023; De Felice et al., 2025; Dumas, 2022; Olarewaju et al., 2023). Such innovation is of particular importance in psychiatry, where reliance on group-level averages under ergodic assumptions obscures the heterogeneous individual- and dyadic-level dynamics that contribute to clinical variation (Fisher et al., 2018; Loth et al., 2016; Meyer-Lindenberg, 2023; Öngür & Paulus, 2025; Schilbach, 2016; Tong, 2019).

Taken together, the tetradic framework provides a multiscale reference frame for evaluating real-time dyadic coordination across normative and clinical samples, while offering a broader architecture for modelling how adaptive systems maintain individuality, exchange information with others, and reorganize through interaction.

## Ethics statement

The behavioural-only cohort was approved by the University of Manitoba Psychology/Sociology Research Board (Protocol HS19206 [P2015:153]). The hyperscanning cohort was approved by the Douglas Mental Health University Institute Research Ethics Board (Protocol 2022-595). All participants provided written informed consent.

## Data and code availability statement

Analysis scripts covering the principal behavioural, tonic EEG, phasic EEG, and information-decomposition analyses, with selected sensitivity checks, are available at https://github.com/Carbonbygravity/tetradic-dynamics. Participant-level EEG analysis inputs and behavioural response-time data may be requested from the corresponding author, subject to applicable consent, ethics, and institutional data-sharing requirements.

## Acknowledgments

We thank the volunteer research team, including Laurence LaFlamme, Sara Toca, Niska Dholabhai, Mitia Andrieux, Laetitia Wang, Ivo Leiva, and Nora Gürke, for their contributions to data collection and study coordination. We thank Dr. Annemarie Wolff and Rémy Ramadour for their assistance during manuscript preparation. We also thank Dr. Jason Leboe-McGowan, Dr. James Frederick Hare, and the University of Manitoba, Faculties of Psychology and Biological Sciences, for their early support in enabling this research. Finally, we thank Aleksandra Atanasova for contributing illustratively to Figures 2 and 8.

This work was supported by research funding awarded to Lena Palaniyappan (LP) and Emmanuel Olarewaju (EO). LP received support from the Monique H. Bourgeois Chair in Developmental Disorders; a New Investigator Supplement from the Healthy Brains, Healthy Lives (HBHL) initiative at McGill University; a salary award from the Fonds de recherche du Québec–Santé (FRQS); the Graham Boeckh Foundation through the Douglas Research Centre, McGill University; the Quebec Bio-Imaging Network (QBIN; grant 35450); and the Canada Foundation for Innovation John R. Evans Leaders Fund, matched by the Government of Quebec, for the project “Studying the Neural Basis of Interpersonal Interactions in Severe Mental Illnesses” (grant 43849). EO received support through an HBHL Fellowship and a Graduate Excellence Recruitment Award from McGill University.

## Declaration of generative AI use

During the preparation of this manuscript, the authors used OpenAI Codex (GPT-6 Sol), accessed in 2026, to assist with code development, debugging, and data-analysis verification. The authors reviewed, verified, and edited all AI-assisted outputs and take full responsibility for the accuracy and integrity of the final manuscript, analyses, and conclusions.

## Author Contributions

**Emmanuel Olarewaju:** Conceptualization; Methodology; Investigation; Data curation; Formal analysis; Validation; Interpretation; Writing – original draft; Writing – review & editing; Visualization; Project administration. **Lena Palaniyappan:** Conceptualization; Methodology; Validation; Interpretation; Writing– review & editing; Funding acquisition; Resources. **Guillaume Dumas:** Conceptualization; Methodology; Validation; Interpretation; Formal analysis; Writing – review & editing.

