## Supplemental Material for "Tetradic Dynamics of Dyadic Sensorimotor Coordination: A Multiscale EEG Hyperscanning Study"

**Supplementary Materials**

**Contents**

**Supplementary S1.** Behavioural Quality Control, Cohort Documentation, and Dyad-Level Sensitivity

**Supplementary S2.** Bayesian Linear Mixed-Effects Model Specification

**Supplementary S3.** EEG Filter Parameters and Ocular Artifact Verification

**Supplementary S4.** Source Reconstruction: Noise Covariance and Forward-Model Details

**Supplementary S5.** Tonic EEG: Top Results, Full Results File, and Dyad-Level Sensitivity

**Supplementary S6.** Phasic EEG Results and Sensitivity Analyses

**Supplementary S7. ΦID: Contrast Definitions, Tensor Construction, and Strict Survivor Results**

**Supplementary S8.** Extended Discussion Notes

### Supplementary S1. Behavioural Quality Control, Cohort Documentation, and Dyad-Level Sensitivity

#### S1.1 Trial-Level Exclusion Criteria and Counts

Behavioural RT data were pre-processed using non-exclusive exclusion criteria applied independently within each participant × role × condition cell. Trials were excluded if reaction time was (1) faster than 150 ms (floor for voluntary motor response), (2) more than 3 SDs above the participant-level cell mean, or (3) outside the P1–P99 range of the participant-level distribution. Criteria were applied non-exclusively (multiple criteria could flag the same trial without double-counting). All exclusions were at the individual-participant level; no joint dyadic latency criterion was applied.

Of 15,360 total trials ($N_{RT}$ = 32; 480 trials/participant), 752 trials (4.90%) were removed, yielding 14,608 retained trials. Post-cleaning trial counts ranged from 114 to 116 per participant × condition cell (median imbalance = 3 trials; maximum = 16). No systematic differences in exclusion rate were observed across Role or Condition cells.

#### S1.2 Combined Behavioural Cohort: Procedural Equivalence

The $N_{RT}$ = 64 combined cohort comprised the current hyperscanning sample (N = 32; collected in 2024) and an independent behavioural dataset from Olarewaju (2021) Experiment 1A (N = 32). The 2021 dataset was collected without concurrent EEG. Key procedural parameters were equivalent across cohorts, as documented in Supplementary Table S1.

### Table S1. Procedural Equivalence of the Two Behavioural Cohorts

| Parameter | Hyperscanning Cohort (N = 32) | Behavioural Cohort (N = 32) |
|---|---|---|
| Collection year | 2024 | 2018 |
| Site | Douglas MHUI, Montréal | U of M, Manitoba |
| EEG concurrent | Yes | No |
| Task software | E-Prime 3.0 | E-Prime 2.0 |
| Controller | PS5 DualSense | PS4 DualShock |
| Trial structure | 2 × 2 within-subject | 2 × 2 within-subject |
| Trials/condition/person | 120 | 120 |
| RT definition (Leader) | Stimulus onset | Stimulus onset |
| RT definition (Follower) | Leader response | Leader response |

*Note.* PS5 = PlayStation 5 DualSense controller (L2/R2 trigger inputs). Both cohorts used identical stimulus-mapping rules for Mirroring and Matching. A Cohort fixed effect was included in the N = 64 ANOVA as a sensitivity check; the Role × Condition interaction pattern was invariant to its inclusion.

### S1.3 Dyad-Level Behavioural RT Sensitivity Analysis

To assess the behavioural contrasts with dyads as independent units, we calculated Condition, Role, and Role × Condition contrasts from each participant's four cell-median response times. The two partners' contrasts were averaged equally within each of 16 dyads. Each dyad-level contrast was assessed using a two-sided one-sample t test. The three planned sensitivity-test p values were not adjusted for multiplicity.

The Condition contrast (Matching − Mirroring) remained positive, indicating slower responses during Matching and a Mirroring RT advantage (mean = 39.51 ms, 95% CI [21.27, 57.75], $t(15) = 4.62$, $p = .000335$). The Role and Role × Condition contrasts did not reach the .05 threshold ($p = .1155$ and .1604, respectively; Table S8, panel A).

### Supplementary S2. Bayesian Linear Mixed-Effects Model Specification

#### S2.1 Model Structure and Priors

Bayesian LMMs were fit to trial-level log-transformed RTs using the Bambi (v0.17.2) interface with a PyMC (v5.27.1) backend. The fixed-effects structure mirrored the frequentist specification:

*log_RT ∼ Role × Condition + (Role + Condition + Role:Condition | Participant_ID) + (1 | Dyad)*

Sum-coded predictors were used throughout (Leader = −0.5, Follower = +0.5; Matching = −0.5, Mirroring = +0.5). Weakly informative priors were assigned: intercept ~ Normal (0, 0.5); fixed effects ~ Normal(0, 0.2); residual and random-effect SDs ~ half-Student-t($\nu = 3$, $\sigma = 0.5$). These priors are weakly informative relative to the observed effect magnitude (main condition contrast on log scale ≈ 0.14) and impose mild shrinkage.

#### S2.2 MCMC Sampling and Convergence

Models were sampled with 4 chains, 3,000 draws per chain, 3,000 tuning steps (target accept = 0.9; random seed = 42). Convergence was assessed via ArviZ (v0.23.4): all fixed-effect parameters achieved $\hat{R} < 1.01$ and effective sample sizes ESS > 1,000. Trace plots showed no pathological features. Posterior predictions were generated for all Role × Condition cells and back-transformed via exponentiation to millisecond scale for interpretability.

#### S2.3 ROPE Definition and Practical-Equivalence Testing

The region of practical equivalence (ROPE) was defined as a ±5% multiplicative change on the raw RT scale, expressed on the log-RT scale as $\delta = \log(1.05) \approx 0.049$ log-ms. Two practically meaningful thresholds were evaluated: (1) P(Follower Mirroring − Matching ≤ −0.05 ratio), and (2) P(Δms ≤ −40 ms). Frequentist two one-sided tests (TOST) with ±25 ms equivalence bounds were applied to participant-level raw RT differences as a complementary equivalence check.

### Supplementary S3. EEG Filter Parameters and Ocular Artifact Verification

#### S3.1 Band-Pass and Notch Filter Specifications

Continuous EEG data were filtered using MNE-Python (v1.11.0) with the parameters in Table S2.

**Table S2. EEG Filter Specifications**

| Parameter | Value |
|---|---|
| Filter type | Zero-phase FIR (firwin design) |
| Window | Hamming (passband ripple: 0.0194; stopband attenuation: 53 dB) |
| Band-pass passband | 1–100 Hz |

| | |
|---|---|
| Lower cutoff (−6 dB) | 0.5 Hz (transition bandwidth: 1 Hz) |
| Upper cutoff (−6 dB) | 112.5 Hz (transition bandwidth: 25 Hz) |

*Note.* Filter length: 3,301 samples (3.301 s at 1,000 Hz). Notch filters at 60 Hz and 120 Hz (FIR firwin). All filters applied to continuous data before epoching.

#### S3.2 Ocular Artifact Invariance Verification

Following ICA and AutoReject, residual eye-movement artifacts were assessed to confirm that blink rates did not vary systematically across experimental conditions. Trial-level blink counts were extracted from synchronized EOG channels. A repeated-measures ANOVA, complemented by paired t-tests and trial-level generalized estimating equations (GEE) with Role and Condition as predictors, confirmed no significant differences in blink rate across the 2 × 2 factorial cells. This rules out condition-correlated EOG contamination as an explanation for observed EEG effects.

### Supplementary S4. Source Reconstruction: Noise Covariance and Forward-Model Details

#### S4.1 Noise Covariance Estimation

A grand-average noise covariance matrix was estimated by pooling pre-stimulus baseline intervals (−1.0 to 0 s) across all retained epochs and participants. Individual-participant noise covariance matrices were regularised using a fixed regularisation parameter ($\lambda = 0.1$) to improve numerical stability. The grand-average matrix was then used to define a common inverse operator applied to all participants, ensuring comparability of source estimates across individuals.

#### S4.2 BEM Conductivity and Forward Solution

The three-layer boundary element model (BEM) used the following conductivity values (standard MNE defaults): brain = 0.3 S/m, skull = 0.006 S/m, scalp = 0.3 S/m. A minimum source–

sensor distance of 5 mm was imposed to reduce numerical instability from superficial dipoles. Sensor coordinates were defined using the standard HydroCel 129-channel montage; a template head-to-MRI transformation was applied for alignment with the *fsaverage* reference space.

#### S4.3 Source Space and Parcellation

Cortical source space used an *oct6* octahedral mesh (~4,098 dipoles/hemisphere). Source estimates (dSPM) were parcellated into 68 ROIs using the Desikan–Killiany atlas. ROI time series were computed as the mean of dipole time courses within each parcel, weighted by the dSPM solution. Table S3 lists all 68 ROIs with their assigned large-scale network.

**Table S3. Desikan–Killiany ROIs and Large-Scale Network Assignments**

| Network | ROI Labels (bilateral) |
|---|---|
| Mirror-Neuron System | pars opercularis; pars triangularis; inferior parietal; precentral; postcentral; superior temporal |
| Salience Network | insula; rostral anterior cingulate; caudal anterior cingulate |
| Frontoparietal Network | Rostral middle frontal; caudal middle frontal; lateral orbitofrontal; superior frontal; inferior parietal; supramarginal |
| Default Mode Network | medial orbitofrontal; frontal pole; posterior cingulate; isthmus cingulate; precuneus; inferior parietal; angular gyrus; temporal pole |
| Sensorimotor | precentral; postcentral; paracentral; supramarginal |
| Temporal/STS | superior temporal; middle temporal; inferior temporal; bankssts; transverse temporal; temporal pole |
| Other | All remaining Desikan–Killiany ROIs not listed above (see MNE-Python atlas documentation for full list) |

*Note.* ROI labels use FreeSurfer/Desikan–Killiany naming convention. Some ROIs appear in multiple networks reflecting their distributed function; for network-engagement analyses, ROIs were assigned to their primary network. Full bilateral ROI set: 34 ROIs × 2 hemispheres = 68 total.

### Supplementary S5. Tonic EEG: Top Results, Full Results File, and Dyad-Level Sensitivity

#### Table S4. FDR-Significant ROI × Band Mixed-Effects Results

| A. Top 10 Most Significant Features (by FDR-corrected p-value) | | | | | | | | |
|---|---|---|---|---|---|---|---|---|
| ROI | Band | Effect | beta (log) | z | LRT $\chi^2$ | p_LRT | p_FDR | Direction |
| rostral middle frontal-rh | gamma | condition | 8.50e−05 | 23.90 | 559.19 | 1.26e−123 | 5.15e−121 | Matching |
| frontal pole-rh | gamma | condition | 7.77e−04 | 23.82 | 555.75 | 7.06e−123 | 1.44e−120 | Matching |
| frontal pole-rh | high beta | condition | 7.69e−04 | 23.35 | 534.20 | 3.45e−118 | 4.69e−116 | Matching |
| Rostral anterior cingulate-rh | gamma | condition | 2.24e−04 | 22.91 | 514.96 | 5.29e−114 | 5.39e−112 | Matching |
| Rostral anterior cingulate-rh | high beta | condition | 2.56e−04 | 21.71 | 463.44 | 8.59e−103 | 7.01e−101 | Matching |
| Superior frontal-rh | gamma | condition | 1.20e−05 | 20.77 | 424.14 | 3.07e−94 | 2.09e−92 | Matching |
| Caudal middle frontal-rh | gamma | condition | 3.70e−05 | 18.53 | 363.02 | 6.21e−81 | 3.62e−79 | Matching |
| Caudal middle frontal-lh | gamma | condition | 3.40e−05 | 18.44 | 349.09 | 6.70e−78 | 3.42e−76 | Matching |
| Rostral anterior cingulate-lh | gamma | condition | 2.13e−04 | 18.40 | 334.11 | 1.23e−74 | 5.56e−73 | Matching |
| frontal pole-lh | high beta | condition | 5.86e−04 | 18.34 | 331.90 | 3.71e−74 | 1.51e−72 | Matching |

| B. Summary Distribution of Significant Features (Total N = 285) | | | | |
|---|---|---|---|---|
| **Band** | **Total Sig.** | **Condition** | **Role** | **ROI Coverage (%)** |
| Low Beta | 63 | 62 | 1 | 92.6% |
| Gamma | 62 | 61 | 1 | 91.2% |
| High Beta | 62 | 62 | 0 | 91.2% |
| Theta | 40 | 36 | 3 | 58.8% |
| Low Alpha | 29 | 29 | 0 | 42.6% |
| High Alpha | 29 | 29 | 0 | 42.6% |

*Note.* Participant-level mixed-effects estimates on log-transformed power (N = 30). Total features tested per effect: 408 (68 ROIs × 6 bands), yielding 1,632 fixed-effect tests across four effects. Significance was determined via Likelihood Ratio Test (LRT) with Benjamini–Hochberg False Discovery Rate (BH–FDR) correction at α = 0.05. Section A displays the top 10 most robust features by corrected probability. Section B summarises the density of significant effects across frequency bands. The complete feature-level results, including all 1,632 effect tests and 285 LRT BH–FDR-significant results, are available at https://github.com/Carbonbygravity/tetradic-dynamics/blob/main/tonic-eeg/results/mixedlm_rt_cov_results_fdr.csv. Positive direction indicates higher power in the Matching condition relative to the Mirroring condition. The theta total includes one RT effect, which is not shown in the Condition and Role columns.

#### S5.1 Dyad-Level Tonic EEG Sensitivity Analysis

We reassessed tonic EEG using dyads as independent units (13,145 trials with usable RT; 30 participants in 15 dyads; 408 ROI × band features). Trial-level power was transformed with log1p. For each participant and feature, we calculated the four Role × Condition cell means and their Condition, Role, and interaction contrasts. We examined unadjusted cell means and two RT-adjusted versions: one using a common within-participant RT slope per feature and another using participant-specific slopes. RT adjustment evaluated each cell mean at that participant's mean RT. The partners' contrasts were averaged equally within each dyad, followed by two-sided one-sample t tests across 15 dyads. Benjamini–Hochberg FDR correction was applied separately to the 408 features for each effect.

Condition estimates favoured Matching in 346 of 408 features with the common RT slope and 347 of 408 with participant-specific slopes. No Condition, Role, or interaction feature survived dyad-level FDR correction in either RT-adjusted version (Table S8, panel B). The unadjusted version yielded a similar Condition direction and also had no FDR survivors. This dyad-level procedure differs from the trial-level mixed model used for the primary analysis, so the difference in survivor counts cannot be attributed to dyad grouping alone. The fixed condition sequence is unchanged by this analysis.

**Supplementary S6. Phasic EEG Results and Sensitivity Analyses**

**S6.1 EMG-Prone Cortical Regions**

To test whether high-frequency clusters depended on potentially EMG-sensitive temporal regions, we repeated the 30-participant high-beta and gamma analyses after excluding two sets of eight ROIs. One set comprised the temporal pole, inferior temporal cortex, banks of the superior temporal sulcus, and transverse temporal cortex bilaterally; the second comprised the temporal pole, fusiform, inferior temporal, and middle temporal cortices bilaterally. The Matching > Mirroring Condition clusters survived both exclusions (high beta: cluster-level FWER $p = .0088$ and .0092; gamma: $p = .0054$ under both). The pre/peri-response high-beta Leader > Follower cluster also survived both ($p = .0232$ and .0242). The post-response high-beta Follower > Leader cluster survived only the second exclusion ($p = .0402$; first exclusion, $p = .0578$). Thus, the Condition clusters did not depend on these excluded ROIs, although this test cannot rule out residual EMG elsewhere.

**S6.2 Frequency-Specific SNR Verification**

Signal-to-noise ratios were verified for each frequency band by comparing the 1/f spectral background against band-averaged power. Bands showing insufficient SNR (defined as band power below 1.5× the interpolated 1/f baseline at that frequency) were flagged. No bands were excluded from the reported analyses; the γ band (30–45 Hz) showed the lowest median SNR (ratio = 2.1) but remained above threshold in all participants.

**Table S5a: Spatiotemporal Cluster-Based Permutation Tests (ROI × Time)**

| Band | Contrast | Cluster | p (FWER) | Direction | Time (ms) | Mass (Σt) | ROIs (n) |
|---|---|---|---|---|---|---|---|
| Gamma | Condition | 1 | 0.0090 | Matching > Mirroring | −1000 to 1000 | 140405.58 | 35 |

| Band | Contrast | Cluster | p (FWER) | Direction | Time (ms) | Mass (Σt) | ROIs (n) |
|---|---|---|---|---|---|---|---|
| High Alpha | Condition | 1 | 0.0014 | Matching > Mirroring | −1000 to 1000 | 46900.32 | 33 |
| High Alpha | Interaction | 1 | 0.0292 | Follower contrast > Leader contrast | 398 to 726 | 4570.29 | 13 |
| High Alpha | Role | 1 | 0.0402 | Leader > Follower | −718 to −277 | 8654.49 | 16 |
| High Alpha | Role | 2 | 0.0060 | Leader > Follower | 15 to 775 | 14228.34 | 29 |
| High Beta | Condition | 1 | 0.0122 | Matching > Mirroring | −1000 to 1000 | 107033.20 | 38 |
| High Beta | Role | 1 | 0.0262 | Leader > Follower | −459 to 208 | 9053.19 | 20 |
| High Beta | Role | 2 | 0.0482 | Follower > Leader | 422 to 856 | -7203.46 | 29 |
| Low Alpha | Condition | 1 | 0.0004 | Matching > Mirroring | −1000 to 1000 | 58746.19 | 38 |
| Low Alpha | Role | 1 | 0.0122 | Leader > Follower | −723 to −268 | 11318.33 | 18 |
| Low Beta | Condition | 1 | 0.0076 | Matching > Mirroring | −1000 to 1000 | 89242.55 | 37 |
| Low Beta | Role | 1 | 0.0044 | Leader > Follower | −322 to 456 | 13677.88 | 24 |
| Theta | Role | 1 | 0.0148 | Leader > Follower | −763 to −275 | 18299.99 | 27 |

### Table S5a (continued): ROIs involved by cluster

| Band \| contrast \| cluster | ROIs involved |
|---|---|
| Gamma \| Condition \| 1 | Caudal anterior cingulate-lh, caudal anterior cingulate-rh, caudal middle frontal-lh, caudal middle frontal-rh, frontal pole-lh, frontal pole-rh, inferior parietal-rh, insula-rh, isthmus cingulate-rh, lateral orbitofrontal-lh, lateral orbitofrontal-rh, medial orbitofrontal-lh, medial orbitofrontal-rh, paracentral-lh, paracentral-rh, parahippocampal-rh, pars opercularis-lh, pars opercularis-rh, pars orbitalis-lh, pars orbitalis-rh, parstriangularis-lh, parstriangularis-rh, postcentral-lh, postcentral-rh, posterior cingulate-lh, posterior cingulate-rh, precentral-rh, precuneus-rh, rostral anterior cingulate-lh, rostral anterior cingulate-rh, rostral middle frontal-lh, rostral middle frontal-rh, superior frontal-lh, superior frontal-rh, superior parietal-rh |
| High Alpha \| Condition \| 1 | bankssts-lh, caudal anterior cingulate-lh, caudal anterior cingulate-rh, caudal middle frontal-lh, cuneus-lh, cuneus-rh, entorhinal-lh, frontal pole-rh, fusiform-lh, inferior temporal-lh, insula-lh, lateral occipital-lh, lateral orbitofrontal-lh, lateral orbitofrontal-rh, medial orbitofrontal-lh, medial orbitofrontal-rh, middle temporal-lh, parahippocampal-lh, pars opercularis-lh, pars orbitalis-lh, parstriangularis-lh, posterior cingulate-lh, precentral-lh, precuneus-lh, rostral anterior cingulate-lh, rostral anterior cingulate-rh, rostral middle frontal-lh, superior frontal-lh, superior parietal-lh, superior temporal-lh, supramarginal-lh, temporal pole-lh, transverse temporal-lh |
| High Alpha \| Interaction \| 1 | bankssts-lh, cuneus-rh, entorhinal-lh, fusiform-lh, inferior temporal-lh, isthmus cingulate-lh, lateral occipital-lh, lingual-lh, middle temporal-lh, parahippocampal-lh, precuneus-lh, superior parietal-lh, temporal pole-lh |
| High Alpha \| Role \| 1 | entorhinal-rh, frontal pole-lh, frontal pole-rh, inferior temporal-rh, lateral orbitofrontal-lh, lateral orbitofrontal-rh, medial orbitofrontal-lh, medial orbitofrontal-rh, pars orbitalis-lh, pars orbitalis-rh, parstriangularis-lh, rostral anterior cingulate-lh, rostral anterior cingulate-rh, rostral middle frontal-rh, temporal pole-lh, temporal pole-rh |

| Band \| contrast \| cluster | ROIs involved |
|---|---|
| High Alpha \| Role \| 2 | Caudal anterior cingulate-lh, caudal middle frontal-lh, cuneus-lh, cuneus-rh, entorhinal-rh, fusiform-lh, fusiform-rh, insula-rh, lateral occipital-lh, lateral occipital-rh, lingual-lh, lingual-rh, middle temporal-rh, paracentral-lh, paracentral-rh, parahippocampal-lh, pericalcarine-lh, pericalcarine-rh, postcentral-lh, posterior cingulate-lh, posterior cingulate-rh, precentral-lh, precentral-rh, precuneus-lh, precuneus-rh, superior parietal-lh, superior temporal-rh, temporal pole-rh, transverse temporal-rh |
| High Beta \| Condition \| 1 | Caudal anterior cingulate-lh, caudal anterior cingulate-rh, caudal middle frontal-lh, caudal middle frontal-rh, cuneus-lh, frontal pole-lh, frontal pole-rh, insula-rh, isthmus cingulate-lh, isthmus cingulate-rh, lateral orbitofrontal-lh, lateral orbitofrontal-rh, medial orbitofrontal-lh, medial orbitofrontal-rh, paracentral-lh, paracentral-rh, parahippocampal-lh, parahippocampal-rh, pars opercularis-lh, pars opercularis-rh, pars orbitalis-lh, pars orbitalis-rh, parstriangularis-rh, postcentral-lh, postcentral-rh, posterior cingulate-lh, posterior cingulate-rh, precentral-lh, precentral-rh, precuneus-rh, rostral anterior cingulate-lh, rostral anterior cingulate-rh, rostral middle frontal-lh, rostral middle frontal-rh, superior frontal-lh, superior frontal-rh, superior parietal-lh, superior parietal-rh |
| High Beta \| Role \| 1 | Caudal anterior cingulate-lh, caudal anterior cingulate-rh, caudal middle frontal-lh, caudal middle frontal-rh, frontal pole-lh, insula-lh, lateral orbitofrontal-lh, medial orbitofrontal-rh, paracentral-lh, paracentral-rh, postcentral-lh, posterior cingulate-lh, posterior cingulate-rh, precentral-lh, precentral-rh, rostral anterior cingulate-lh, rostral anterior cingulate-rh, superior frontal-lh, superior temporal-lh, supramarginal-lh |
| High Beta \| Role \| 2 | bankssts-rh, caudal anterior cingulate-rh, caudal middle frontal-lh, caudal middle frontal-rh, frontal pole-lh, fusiform-rh, insula-lh, lateral orbitofrontal-lh, medial orbitofrontal-rh, middle temporal-rh, paracentral-rh, parstriangularis-lh, postcentral-lh, postcentral-rh, posterior cingulate-lh, posterior cingulate-rh, precentral-rh, rostral anterior cingulate-lh, rostral anterior cingulate-rh, rostral middle frontal-lh, rostral middle frontal-rh, superior frontal-rh, superior temporal-lh, supramarginal-lh, supramarginal-rh, temporal pole-lh, temporal pole-rh, transverse temporal-lh, transverse temporal-rh |
| Low Alpha \| Condition \| 1 | bankssts-lh, caudal anterior cingulate-lh, caudal anterior cingulate-rh, caudal middle frontal-lh, cuneus-lh, cuneus-rh, entorhinal-lh, entorhinal-rh, fusiform-lh, fusiform-rh, inferior temporal-lh, inferior temporal-rh, insula-lh, lateral occipital-lh, lateral occipital-rh, lateral orbitofrontal-lh, lateral orbitofrontal-rh, medial orbitofrontal-lh, medial orbitofrontal-rh, middle temporal-lh, middle temporal-rh, parahippocampal-lh, pars opercularis-lh, pars orbitalis-rh, parstriangularis-lh, posterior cingulate-lh, posterior cingulate-rh, precentral-lh, precuneus-lh, rostral anterior cingulate-lh, rostral anterior cingulate-rh, superior frontal-lh, superior parietal-lh, superior temporal-lh, supramarginal-lh, temporal pole-lh, temporal pole-rh, transverse temporal-lh |
| Low Alpha \| Role \| 1 | entorhinal-lh, entorhinal-rh, frontal pole-lh, frontal pole-rh, lateral orbitofrontal-lh, lateral orbitofrontal-rh, medial orbitofrontal-lh, medial orbitofrontal-rh, pars opercularis-lh, pars orbitalis-lh, pars orbitalis-rh, parstriangularis-lh, rostral anterior cingulate-lh, rostral anterior cingulate-rh, rostral middle frontal-lh, rostral middle frontal-rh, temporal pole-lh, temporal pole-rh |

| Band \| contrast \| cluster | ROIs involved |
|---|---|
| Low Beta \| Condition \| 1 | Caudal anterior cingulate-lh, caudal anterior cingulate-rh, caudal middle frontal-lh, caudal middle frontal-rh, entorhinal-rh, frontal pole-rh, fusiform-lh, insula-lh, insula-rh, isthmus cingulate-lh, lateral orbitofrontal-lh, lateral orbitofrontal-rh, medial orbitofrontal-lh, medial orbitofrontal-rh, paracentral-lh, parahippocampal-lh, pars opercularis-lh, pars opercularis-rh, pars orbitalis-lh, pars orbitalis-rh, parstriangularis-lh, parstriangularis-rh, postcentral-lh, postcentral-rh, posterior cingulate-lh, posterior cingulate-rh, precentral-lh, precentral-rh, precuneus-lh, rostral anterior cingulate-lh, rostral anterior cingulate-rh, rostral middle frontal-lh, rostral middle frontal-rh, superior frontal-lh, superior frontal-rh, superior temporal-rh, transverse temporal-lh |
| Low Beta \| Role \| 1 | bankssts-rh, caudal middle frontal-lh, cuneus-lh, cuneus-rh, isthmus cingulate-rh, lateral occipital-lh, lateral occipital-rh, lingual-lh, lingual-rh, paracentral-lh, paracentral-rh, parahippocampal-rh, pericalcarine-lh, pericalcarine-rh, postcentral-lh, postcentral-rh, posterior cingulate-lh, posterior cingulate-rh, precentral-lh, precentral-rh, precuneus-lh, precuneus-rh, superior parietal-lh, superior parietal-rh |
| Theta \| Role \| 1 | Caudal anterior cingulate-rh, caudal middle frontal-lh, caudal middle frontal-rh, entorhinal-lh, entorhinal-rh, frontal pole-lh, frontal pole-rh, inferior temporal-lh, insula-lh, lateral orbitofrontal-lh, lateral orbitofrontal-rh, medial orbitofrontal-lh, medial orbitofrontal-rh, pars opercularis-lh, pars opercularis-rh, pars orbitalis-lh, pars orbitalis-rh, parstriangularis-lh, parstriangularis-rh, rostral anterior cingulate-lh, rostral anterior cingulate-rh, rostral middle frontal-lh, rostral middle frontal-rh, superior frontal-rh, temporal pole-lh, temporal pole-rh, transverse temporal-lh |

*Note.* Cluster-based permutation tests used 5,000 permutations (two-sided, $\alpha = .05$) and ROI adjacency. p-values are cluster-level familywise error-corrected within each band × contrast map; no correction across maps was applied. Condition is Matching − Mirroring; Role is Leader − Follower response-event epochs; the interaction compares the Follower and Leader condition contrasts. The Direction column gives the observed sign of each contrast. ROI membership describes cluster extent, not separate significance for each ROI.

**Table S5b. Space × Frequency Cluster Analyses (ROI × Band)**

| Contrast | Cluster | p-value | Direction | Peak ROI | Band | Elements |
|---|---|---|---|---|---|---|
| Condition | 1 | 0.0252 | Matching > Mirroring | Caudal anterior cingulate-rh | Low Beta | 20 |
| Condition | 2 | 0.0216 | Matching > Mirroring | Frontal pole-rh | High Beta | 22 |
| Condition | 3 | 0.0126 | Matching > Mirroring | Frontal pole-rh | Gamma | 25 |
| Interaction | — | — | No significant clusters | — | — | — |
| Role | — | — | No significant clusters | — | — | — |

**ROIs Involved**

**Cluster 1 (Low Beta):** caudal anterior cingulate-lh, caudal anterior cingulate-rh, caudal middle frontal-lh, caudal middle frontal-rh, lateral orbitofrontal-lh, lateral orbitofrontal-rh, medial orbitofrontal-lh, medial orbitofrontal-rh, pars opercularis-lh, pars opercularis-rh, pars orbitalis-lh, pars orbitalis-rh, parstriangularis-lh, posterior cingulate-rh, precentral-rh, rostral anterior cingulate-lh, rostral anterior cingulate-rh, rostral middle frontal-lh, rostral middle frontal-rh, superior frontal-lh

**Cluster 2 (High Beta):** caudal anterior cingulate-lh, caudal anterior cingulate-rh, caudal middle frontal-lh, caudal middle frontal-rh, frontal pole-lh, frontal pole-rh, lateral orbitofrontal-lh, lateral orbitofrontal-rh, medial orbitofrontal-lh, medial orbitofrontal-rh, paracentral-rh, pars opercularis-rh, pars orbitalis-lh, pars orbitalis-rh, postcentral-rh, posterior cingulate-rh, precentral-rh, rostral anterior cingulate-lh, rostral anterior cingulate-rh, rostral middle frontal-lh, rostral middle frontal-rh, superior frontal-lh

**Cluster 3 (Gamma):** caudal anterior cingulate-lh, caudal anterior cingulate-rh, caudal middle frontal-lh, caudal middle frontal-rh, frontal pole-lh, frontal pole-rh, insula-rh, medial orbitofrontal-lh, medial orbitofrontal-rh, paracentral-rh, pars opercularis-lh, pars opercularis-rh, pars orbitalis-lh, pars orbitalis-rh, parstriangularis-rh, postcentral-rh, posterior cingulate-lh, posterior cingulate-rh, precentral-rh, rostral anterior cingulate-lh, rostral anterior cingulate-rh, rostral middle frontal-lh, rostral middle frontal-rh, superior frontal-lh, superior frontal-rh

*Note.* Space × frequency cluster permutation testing performed on time-averaged source power (ROI × band). 5,000 permutations; α = .05; family-wise error controlled across 408 features (68 ROIs × 6 bands). Direction "Matching > Mirroring" indicates greater time-averaged source power in the Matching condition.

#### S6.3 RT-Controlled Phasic Sensitivity Analysis

Trial-level source power was matched to reaction time (RT) for the 30 participants in the source-level analysis. Of 14,220 available trials, 13,145 had usable RT values. For each participant and each ROI × frequency-band × time-point feature, log1p-transformed trial power was regressed on within-participant standardized RT. Power was adjusted to RT z = 0 before averaging trials into the verified Leader–Matching, Leader–Mirroring, Follower–Matching, and Follower–Mirroring cells. The resulting participant-level cell estimates were tested using the same 18 band × contrast spatiotemporal cluster tests as the primary analysis (5,000 permutations; cluster-level familywise-error correction within each map).

All five Matching > Mirroring Condition clusters remained significant within their respective maps. Seven role-related or interaction clusters also met the within-map threshold, although their locations and time windows did not uniformly reproduce the primary clusters. The full set of significant RT-controlled clusters is reported in Table S7. These results support the condition direction after accounting for trial-level RT, while indicating less stable role-specific cluster detail.

#### S6.4 Sensitivity to Correction Across Maps

As a separate, conservative multiplicity check, each saved within-map cluster p value was multiplied by 18, the number of frequency-band × contrast maps in each analysis, and capped at 1. The primary and RT-controlled analyses were treated as separate families. This Bonferroni calculation did not rerun the cluster tests or alter their reported within-map p values.

In the primary analysis, two of 13 within-map significant clusters survived correction across 18 maps: the low-alpha and high-alpha Matching > Mirroring Condition clusters. In the RT-controlled analysis, three of 12 survived: the low-alpha and high-alpha Matching > Mirroring Condition clusters and a peri-response low-beta Leader > Follower cluster. No primary Role or Role × Condition cluster survived this additional across-map correction. Table S7 reports both p values for every within-map significant cluster.

**Table S7. Phasic sensitivity analyses: within-map and across-map cluster p values**

| Analysis | Band | Contrast | Window (ms) | Within-map p | Across-map p | Survives |
|---|---|---|---|---|---|---|
| Primary | Theta | Role | -763 to -275 | 0.0148 | 0.2664 | No |
| Primary | Low Alpha | Condition | -1000 to 1000 | 0.0004 | 0.0072 | Yes |
| Primary | Low Alpha | Role | -723 to -268 | 0.0122 | 0.2196 | No |
| Primary | High Alpha | Condition | -1000 to 1000 | 0.0014 | 0.0252 | Yes |
| Primary | High Alpha | Role | -718 to -277 | 0.0402 | 0.7236 | No |
| Primary | High Alpha | Role | 15 to 775 | 0.0060 | 0.1080 | No |
| Primary | High Alpha | Interaction | 398 to 726 | 0.0292 | 0.5256 | No |
| Primary | Low Beta | Condition | -1000 to 1000 | 0.0076 | 0.1368 | No |
| Primary | Low Beta | Role | -322 to 456 | 0.0044 | 0.0792 | No |
| Primary | High Beta | Condition | -1000 to 1000 | 0.0122 | 0.2196 | No |
| Primary | High Beta | Role | -459 to 208 | 0.0262 | 0.4716 | No |
| Primary | High Beta | Role | 422 to 856 | 0.0482 | 0.8676 | No |
| Primary | Gamma | Condition | -1000 to 1000 | 0.0090 | 0.1620 | No |
| RT-controlled | Theta | Role | -742 to -312 | 0.0226 | 0.4068 | No |
| RT-controlled | Theta | Role | 540 to 1000 | 0.0496 | 0.8928 | No |
| RT-controlled | Low Alpha | Condition | -1000 to 1000 | 0.0002 | 0.0036 | Yes |
| RT-controlled | Low Alpha | Role | -711 to -257 | 0.0168 | 0.3024 | No |
| RT-controlled | High Alpha | Condition | -1000 to 1000 | 0.0018 | 0.0324 | Yes |
| RT-controlled | High Alpha | Role | -38 to 702 | 0.0080 | 0.1440 | No |
| RT-controlled | Low Beta | Condition | -1000 to 1000 | 0.0066 | 0.1188 | No |
| RT-controlled | Low Beta | Role | -224 to 530 | 0.0016 | 0.0288 | Yes |
| RT-controlled | High Beta | Condition | -1000 to 1000 | 0.0134 | 0.2412 | No |
| RT-controlled | High Beta | Role | -447 to 265 | 0.0304 | 0.5472 | No |
| RT-controlled | Gamma | Condition | -1000 to 1000 | 0.0122 | 0.2196 | No |
| RT-controlled | Gamma | Interaction | -1000 to -650 | 0.0464 | 0.8352 | No |

*Note.* The table includes clusters with within-map cluster-level FWER $p < .05$. Across-map p = min(1, 18 × within-map p), calculated separately for the primary and RT-controlled analyses. "Survives" indicates across-map $p < .05$. Positive Condition clusters indicate Matching > Mirroring; the reported low-beta Role cluster indicates Leader > Follower. Time windows are referenced to the corresponding response-period onset.

**S6.5 Temporal Centering of the Condition Contrast**

To assess whether the full-epoch Condition clusters could be explained solely by a sustained Matching–Mirroring difference, we subtracted each participant's mean Condition contrast over −1000 to +1000 ms separately for each ROI and band. We then repeated the six Condition cluster tests using the original ROI adjacency and 5,000 permutations, with cluster-level FWER correction within each band map. This check removes an epoch-constant difference; it is not pre-cue baseline normalization.

After centering, a positive high-beta cluster (−561 to −77 ms; $p = .0278$; 9 ROIs) and a negative gamma cluster (+885 to +1000 ms; $p = .0330$; 28 ROIs) were significant within their respective maps. No other band yielded a significant cluster. Positive and negative describe deviations from each participant's full-epoch Matching–Mirroring mean, not the direction of the original, uncentered contrast. Thus, the data retain limited within-epoch variation beyond a constant Condition difference; this check alone does not establish a separate response-evoked mechanism.

**S6.6 Dyad-Level Phasic EEG Sensitivity Analysis**

We repeated the phasic Condition, Role, and Role × Condition tests using dyads as the independent units. The two partners' participant-level contrast maps were averaged within each of 15 dyads. Spatiotemporal cluster tests used whole-dyad sign flips, the same ROI adjacency, and 5,000 permutations per band × contrast map. Cluster-level familywise error was controlled within each map; no additional correction across the 18 maps was applied in this sensitivity analysis.

Thirteen clusters met the within-map threshold: five Condition, six Role, and two Role × Condition clusters (Table S8, panel C). All five Condition clusters retained the Matching > Mirroring direction and spanned the full response-period-aligned epoch. Several Role clusters before response-period onset and the high-alpha interaction after onset were also present, but the detailed role-related pattern was not identical to the primary analysis. The late high-beta Follower > Leader Role cluster after onset did not reach the dyad-level within-map threshold. A high-beta interaction before onset appeared in this sensitivity analysis and is not treated as a primary finding.

#### Table S8. Dyad-Level Sensitivity Analyses of Behavioural RT, Tonic EEG, and Phasic EEG

#### A. Behavioural RT (16 dyads)

| Contrast | Mean (ms) | 95% CI (ms) | t(15) | p |
|---|---|---|---|---|
| Condition | 39.51 | 21.27 to 57.75 | 4.62 | .000335 |
| Role | −37.32 | −84.93 to 10.29 | −1.67 | .1155 |
| Role × Condition | 32.45 | −14.39 to 79.30 | 1.48 | .1604 |

#### B. Tonic EEG (15 dyads; 408 features per effect)

| Cell-mean version | Effect | Positive / 408 | Raw p < .05 | BH-FDR q < .05 |
|---|---|---|---|---|
| Unadjusted | Condition | 347 | 95 | 0 |
| Unadjusted | Role | 248 | 29 | 0 |
| Unadjusted | Role × Condition | 193 | 11 | 0 |
| Common RT slope | Condition | 346 | 96 | 0 |
| Common RT slope | Role | 255 | 33 | 0 |
| Common RT slope | Role × Condition | 185 | 15 | 0 |
| Participant-specific RT slopes | Condition | 347 | 94 | 0 |
| Participant-specific RT slopes | Role | 265 | 23 | 0 |
| Participant-specific RT slopes | Role × Condition | 168 | 13 | 0 |

#### C. Phasic EEG (15 dyads; within-map significant clusters)

| Contrast | Band | Time window (ms) | Cluster p | ROIs |
|---|---|---|---|---|
| Condition | Low alpha | −1000 to +1000 | .0016 | 37 |
| Condition | High alpha | −1000 to +1000 | .0016 | 35 |
| Condition | Low beta | −1000 to +1000 | .0118 | 39 |
| Condition | High beta | −1000 to +1000 | .0160 | 32 |
| Condition | Gamma | −1000 to +1000 | .0130 | 32 |
| Role | Theta | −768 to −293 | .0066 | 27 |
| Role | Low alpha | −741 to −271 | .0048 | 18 |
| Role | High alpha | −715 to −260 | .0234 | 15 |
| Role | High alpha | +23 to +546 | .0008 | 31 |
| Role | Low beta | −319 to +421 | .0022 | 22 |
| Role | High beta | −471 to +227 | .0308 | 19 |
| Role × Condition | High alpha | +416 to +722 | .0210 | 9 |
| Role × Condition | High beta | −831 to −341 | .0362 | 10 |

*Note.* In panel A, positive Condition values indicate slower RT during Matching; p values were not adjusted across the three contrasts. In panel B, "positive" indicates the specified contrast direction; raw-p counts are not FDR-significant findings. In panel C, positive Condition denotes Matching > Mirroring, positive Role denotes Leader-response > Follower-response, and positive interaction denotes a larger Matching − Mirroring contrast around

Follower than Leader responses. Panel C p values are cluster-level familywise error-corrected within each map, not across the 18 maps.

### Supplementary S7. ΦID: Contrast Definitions, Tensor Construction, and Strict Survivor Results

#### S7.1 ΦID Contrast Definitions

Four contrasts were constructed from the four response-period × condition groups (Leader–Mirroring [LMR], Leader–Matching [LMA], Follower–Matching [FMA], Follower–Mirroring [FMR]):

*Condition (Matching − Mirroring)*: [(LMA + FMA) − (LMR + FMR)] / 2

*Leader − Follower within Mirroring*: LMR − FMR

*Leader − Follower within Matching*: LMA − FMA

*Leader − Follower averaged across conditions*: [(LMR + LMA) − (FMA + FMR)] / 2

A separate response-period × condition interaction was tested as (LMA − LMR) − (FMA − FMR). Positive interaction values indicate a larger Matching–Mirroring difference during Leader than Follower response periods. These contrasts define dyad-wise difference time series applied per ROI for each ΦID atom, frequency band, and source configuration (homologous, cross-hemispheric) in the temporal cluster permutation tests.

#### S7.2 ΦID Tensor Construction

For each dyad, source-level ROI time series were transformed to Morlet power at 1-Hz intervals from 4 to 45 Hz and averaged within the six frequency bands specified in Section 3.9. The band-power means were then log-transformed [log(1 + power)]. Epochs were grouped by Leader or Follower response period and Mirroring or Matching condition, with X assigned to the task Leader and Y to the Follower for each epoch according to its event suffix. For each ROI pairing, frequency band and time point, the four selected ΦID atoms were estimated across paired trials using the Gaussian/MMI estimator. The four estimator variables were scaled by their across-trial standard deviations. Homologous and cross-hemispheric pairings were estimated separately, yielding dyad-level values indexed by response-period × condition group, ROI, band, time and atom for each geometry. The primary analysis used $\tau = 1$ sample (1 ms at 1,000 Hz), corresponding to a 2 ms past–future separation; time points spanned −999 to +999 ms relative to response-period onset.

**Table S6a. Integrated Information Decomposition (ΦID) — Feature-level statistics for Tier A and Tier B targets**

| Tier | ΦID | Geometry | Contrast | Frequency Band | ROI | Time Window (ms) | Duration (ms) | Dyads Passing Trial-Shuffle | Dyads Passing Time-Shift |
|---|---|---|---|---|---|---|---|---|---|
| Four condition and response-period contrasts | | | | | | | | | |
| Tier A | rtr | H | Matching > Mirroring | theta | postcentral-rh | −999 to −879 | 120 | 4 | 6 |
| Tier A | yty | C | Matching > Mirroring | high beta | insula-lh | 27 to 197 | 170 | 4 | 5 |
| Tier B | rtr | C | Matching > Mirroring | low beta | medialorbitofrontal-lh | 373 to 541 | 168 | 5 | 0 |
| Tier A | sts | H | Follower > Leader, Mirroring | theta | rostralanteriorcingulate-rh | −950 to −813 | 137 | 5 | 4 |

| Tier | ΦID | Geometry | Contrast | Frequency Band | ROI | Time Window (ms) | Duration (ms) | Dyads Passing Trial-Shuffle | Dyads Passing Time-Shift |
|---|---|---|---|---|---|---|---|---|---|
| Tier B | rtr | C | Follower > Leader, Mirroring | gamma | isthmuscingulate-rh | 235 to 409 | 174 | 4 | 1 |
| Tier A | rtr | C | Leader > Follower, Matching | low alpha | paracentral-rh | −195 to −82 | 113 | 4 | 3 |
| Tier B | rtr | H | Leader > Follower, Matching | low beta | isthmuscingulate-lh | 231 to 408 | 177 | 4 | 2 |
| Tier A | rtr | H | Follower period > Leader period | gamma | lateraloccipital-rh | −228 to −60 | 168 | 4 | 5 |
| Tier A | rtr | H | Follower period > Leader period | low beta | supramarginal-lh | −735 to −615 | 120 | 4 | 4 |
| Tier A | rtr | H | Follower period > Leader period | high alpha | temporalpole-rh | −193 to −88 | 105 | 4 | 8 |
| Tier A | rtr | C | Leader period > Follower period | high alpha | entorhinal-rh | 221 to 337 | 116 | 4 | 6 |
| Tier A | rtr | C | Follower period > Leader period | high alpha | parstriangularis-lh | −20 to 91 | 111 | 4 | 4 |

| Tier | ΦID | Geometry | Contrast | Frequency Band | ROI | Time Window (ms) | Duration (ms) | Dyads Passing Trial-Shuffle | Dyads Passing Time-Shift |
|---|---|---|---|---|---|---|---|---|---|
| Tier A | xtx | H | Leader period > Follower period | theta | bankssts-lh | 217 to 321 | 104 | 5 | 3 |
| Tier B | xtx | C‡ | Follower period > Leader period | low beta | superiortemporal-rh | 156 to 233 | 77 | 5 | 2 |
| Tier B | sts | H | Follower period > Leader period | theta | rostralanterior cingulate-rh | −951 to −774 | 177 | 5 | 2 |
| Tier B | yty | H | Follower period > Leader period | low alpha | superiorfrontal-rh | 555 to 622 | 67 | 4 | 0 |
| Separate response-period × condition interaction | | | | | | | | | |
| Tier B | xtx | H‡ | Follower Δ > Leader Δ | gamma | frontalpole-lh | −743 to −631 | 112 | 4 | 1 |
| Tier B | rtr | C | Follower Δ > Leader Δ | theta | caudalmiddlefrontal-lh | 435 to 606 | 171 | 4 | 0 |
| Tier A | rtr | C | Leader Δ > Follower Δ | low alpha | inferiortemporal-lh | −475 to −343 | 132 | 4 | 7 |

*Note.* This table reports 16 targets from the four condition and response-period contrasts and three from the separate response-period × condition interaction. Time windows are relative to response-period onset. H = homologous; C = cross-hemispheric. For the interaction, Δ denotes Matching minus Mirroring. Tier A denotes targets meeting trial-shuffle recurrence in at least four dyads and time-shift recurrence in at least three dyads. Tier B meets the trial-shuffle criterion but not both Tier A criteria. ‡The pairing was identified in both geometries before surrogate controls; only the geometry shown reached Tier A or B.

#### S7.3 Lag, Embedding, and Signal-Timescale Robustness for ΦID

A potential concern in time-resolved ΦID analyses is that a unit lag may capture near-instantaneous statistical dependence or local autocorrelation rather than predictive dynamics, particularly when the selected lag is shorter than the relevant timescales of the signal under analysis (Ragwitz & Kantz, 2002; Wibral et al., 2013). This concern is especially relevant for band-limited power-envelope signals, because filtering and envelope extraction introduce temporal smoothness that can influence lag-dependent information-theoretic estimates (Weber et al., 2017). More generally, choices regarding delay, embedding, filtering, and sampling should be evaluated explicitly when distinguishing predictive dependence from local signal continuity (Fraser & Swinney, 1986; Ragwitz & Kantz, 2002). Accordingly, $\tau = 1$ was interpreted as the finest sampled local past–future transition within the response-period-aligned dyadic trajectory.

##### S7.3.1 Primary ΦID lag definition

In the primary ΦID analysis, the dyadic transition was defined as:

$S = [X(t - \tau), Y(t - \tau)]$ and $T = [X(t + \tau), Y(t + \tau)]$

where X denotes the role-canonical Leader signal and Y denotes the role-canonical Follower signal. With data sampled at 1000 Hz, the primary setting $\tau = 1$ corresponds to adjacent past/future samples around each local time point. This was treated as a local time-resolved transition, not as a full oscillatory-cycle embedding or an estimate of physiological transmission delay.

In this formulation, ΦID evaluates whether the dyad's local past–future structure preserves synergistic, redundant, leader-unique, or follower-unique information geometry around the response period. This interpretation is consistent with the broader ΦID framework, which decomposes statistical dependencies between a multivariate system's past and future into distinct redundant, unique, and synergistic information modes (Mediano et al., 2025; Varley, 2023). Varley (2023) further demonstrated that both average and local ΦID quantities can reveal temporally evolving information-dynamic structure in neural data.

Accordingly, the present time-resolved analysis interprets the surviving effects as local past–future information geometries within the response-period-aligned dyadic trajectory.

**S7.3.2 Tau-specific surrogate validation**

To test whether effects were specific to $\tau = 1$, we recomputed ΦID for the 19 primary Tier A/B targets across longer $\tau$ values:

$\tau = 1, 2, 4, 8, 16, 32$

These correspond to past–future separations of 2, 4, 8, 16, 32, and 64 ms. Each target was evaluated over a common time window across $\tau$ values. Trial-shuffle and time-shift controls were conducted separately at each $\tau$, using the target's primary contrast direction and the same recurrence criteria as the primary analysis. The four main contrasts (16 targets) and the separate response-period × condition interaction (three targets) were summarized separately.

All 19 targets retained their contrast direction across the tested $\tau$ values, although their robustness tiers varied. Among the four main contrasts, the number meeting Tier A or B criteria decreased from 16 at $\tau = 1$ to five at $\tau = 32$; for the interaction, it decreased from three to one. The right postcentral theta target used a common-support window of −968 to −879 ms, compared with its primary window of −999 to −879 ms. Target-level tiers and surrogate recurrence counts by $\tau$ are provided in Supplementary Table S6c.

### S7.4. History Embedding, Selection-Adjusted Surrogates, and Signal-Timescale Diagnostics

Following the τ sensitivity analysis in Section S7.3, we examined history/τ selection for 17 primary Tier A/B targets: 14 from the four condition and response-period contrasts and three from the separate response-period × condition interaction. Two boundary targets were excluded because their windows did not accommodate the full candidate grid. Signal-timescale diagnostics were examined for 11 primary Tier A targets.

#### S7.4.1 History/tau embedding selection

To address the concern that fixing history length to one sample may under-embed a band-limited envelope, we performed a Ragwitz-style history/tau selection diagnostic (Ragwitz & Kantz, 2002). Candidate embeddings varied both history depth and τ:

*history_length* = 1, 2, 3

$\tau = 1, 2, 4, 8, 16, 32, 64$

For each candidate, local KNN prediction error was used to evaluate how well the role-canonical dyadic past predicted the dyadic future. Selection used fivefold cross-validation of whole trials, with $k = 10$ and normalization based on each training fold. Scores were weighted equally across actors, response-period × condition groups and dyads, with a maximum of 25,000 prediction points per candidate. Because the intended ΦID interpretation depends on a two-source/two-target geometry, multistep history was compressed into one role-specific history state per participant:

$$\text{Leader history state} = \text{mean}[X(t - \tau), X(t - 2\tau), ..., X(t - H\tau)]$$

$$\text{Follower history state} = \text{mean}[Y(t - \tau), Y(t - 2\tau), ..., Y(t - H\tau)]$$

This preserved the dyadic ΦID source structure:

[Leader history state, Follower history state] → [Leader future, Follower future]

For all 17 eligible targets, the observed-data selection was *history_length* = 1 and τ = 1.

#### S7.4.2 Selection-adjusted surrogate validation

Because embedding selection itself can capitalize on noise, the history/τ selection procedure was repeated for each surrogate before surrogate ΦID was computed. The null distributions therefore included both the disruption of trial correspondence or temporal alignment and the freedom to select history/τ. Trial-shuffle controls used 100 repetitions; time-shift controls were conducted for targets meeting the group trial-shuffle criterion of $p \leq .10$. The original contrast direction and across-dyad recurrence criteria were retained, with the four main contrasts and interaction summarized separately.

Among the 14 eligible main-contrast targets, the history-specific controls yielded eight Tier A, five Tier B and one Tier C classification. The three interaction targets yielded one Tier A and two Tier B classifications (Supplementary Table S6d).

The homologous right rostral anterior cingulate theta Synergy target illustrates a window-dependent exception. Its primary Leader–Follower contrast was Tier B over −951 to −774 ms. The history analysis used the shorter common-support interval −808 to −774 ms and yielded Tier C. The observed data and all 100 trial surrogates selected history_length = 1 and τ = 1; fixing that setting reproduced the shortened-window result. Thus, the change in tier followed the restriction of the averaging window, not selection of a different history or lag. This target did not meet the history-specific trial-shuffle criterion for time-shift testing. Its primary full-window Tier B classification remains unchanged.

#### S7.4.3 Signal-timescale diagnostics

Finally, we estimated signal-timescale diagnostics to contextualize the selected short lag. Following the use of mutual information for delay-coordinate selection in nonlinear time-series analysis (Fraser & Swinney, 1986), we computed autocorrelation zero crossings, autocorrelation first local minima, and mutual-information first local minima within each significant feature window and across the full epoch. These diagnostics were calculated separately for Leader and Follower signals in the 11 primary Tier A targets.

Across target- and actor-specific summaries, the median first mutual-information minimum was 4–5 ms within feature windows and 53.5–57 ms across the full epoch. Full-epoch median autocorrelation zero crossings ranged from 149.5 to 184.5 ms (Supplementary Table S6e). These measures describe different temporal spans of the band-power signals. The primary $\tau = 1$ setting represents a local past–future transition, not a full oscillatory-cycle embedding or an estimate of physiological transmission delay.

The history and signal-timescale analyses therefore provide complementary assessments of the local ΦID transition. Their sensitivity classifications are reported separately from the primary surrogate tiers.

**S7.5 Chronological Task-Half Comparisons for Unique-Information Targets**

Chronological task-half comparisons were applied to five primary Tier A/B unique-information targets: four from the main contrasts and one from the separate response-period × condition interaction. Participant 1 served as Leader in Blocks 1–2 and Participant 2 in Blocks 3–4; the condition sequence was Mirroring–Matching–Matching–Mirroring. Thus, participant identity, chronological half, and condition order changed together across the two halves.

For each target, ΦID was recomputed separately from the paired source inputs in the early and late halves, retaining X as the task Leader and Y as the Follower for each epoch. The target's original ROI pairing, frequency band, time window and $\tau = 1$ setting were retained. Its signed contrast was averaged over the time window, and the late-minus-early difference was calculated within each of 15 dyads.

Early–late differences were assessed using all 32,768 paired sign flips, with two-sided tests and Holm correction across the five targets. Effect sizes and 95% bootstrap intervals were also calculated. A companion analysis repeated the comparisons 100 times with trial counts matched between halves within each dyad and response-period × condition group. None of the five full-data comparisons met the corrected threshold (all Holm-adjusted p = 1.00; Supplementary Table S6b).

#### Table S6b. Chronological Task-Half Comparisons for Unique-Information ΦID Targets

| Analysis | Atom | Geometry; ROI pair | Band | Window (ms) | Late − early | $d^z$ | 95% bootstrap CI | Raw p | Holm p |
|---|---|---|---|---|---|---|---|---|---|
| Matching − Mirroring | yty | Cross-hemispheric; insula L–R | High beta | 27–197 | −0.0251 | −0.1805 | −0.0890 to 0.0451 | .4961 | 1.00 |
| Leader − Follower, averaged across conditions | xtx | Cross-hemispheric; superior temporal R–L | Low beta | 156–233 | 0.0481 | 0.2581 | −0.0396 to 0.1407 | .3394 | 1.00 |
| Leader − Follower, averaged across conditions | xtx | Homologous; bank of superior temporal sulcus L–L | Theta | 217–321 | 0.0058 | 0.0320 | −0.0843 to 0.0926 | .9015 | 1.00 |
| Leader − Follower, averaged across conditions | yty | Homologous; superior frontal R–R | Low alpha | 555–622 | −0.0081 | −0.0373 | −0.1217 to 0.0922 | .8859 | 1.00 |
| Separate response-period × condition interaction | xtx | Homologous; frontal pole L–L | Gamma | −743 to −631 | 0.0038 | 0.0171 | −0.1095 to 0.1017 | .9494 | 1.00 |

*Note.* Values are late-minus-early changes in each target's signed contrast, averaged over its stated time window across 15 dyads. Early and late refer to chronological Blocks 1–2 and 3–4. Two-sided p values were obtained using paired sign flips; Holm correction was applied across the five targets. The 95% intervals are bootstrap intervals. Participant identity, chronological half, and condition order changed together across the halves.

### Table S6c. τ-specific surrogate recurrence and robustness tiers for ΦID targets

| Target | τ=1 | τ=2 | τ=4 | τ=8 | τ=16 | τ=32 |
|---|---|---|---|---|---|---|
| **Four condition and response-period contrasts** | | | | | | |
| Matching–Mirroring; rtr; H; postcentral R; θ | A (4/6) | A (4/6) | A (4/6) | A (4/6) | C (3/6) | C (3/6) |
| Matching–Mirroring; yty; C; insula L; high β | A (4/5) | A (4/5) | A (4/5) | A (4/5) | A (4/5) | C (3/5) |
| Matching–Mirroring; rtr; C; medial orbitofrontal L; low β | B (5/0) | B (4/0) | B (4/0) | B (4/2) | B (4/1) | B (4/1) |
| Leader–Follower within Mirroring; sts; H; rostral anterior cingulate R; θ | A (5/4) | A (5/4) | A (5/4) | A (5/4) | A (5/4) | C (3/3) |
| Leader–Follower within Mirroring; rtr; C; isthmus cingulate R; γ | B (4/1) | B (4/2) | B (4/2) | B (4/2) | B (4/2) | B (4/2) |
| Leader–Follower within Matching; rtr; H; isthmus cingulate L; low β | B (4/2) | B (4/2) | B (4/2) | C (2/2) | C (2/2) | C (2/1) |
| Leader–Follower within Matching; rtr; C; paracentral R; low α | A (4/3) | A (4/3) | A (4/3) | C (3/3) | C (3/3) | C (3/4) |
| Leader–Follower averaged across conditions; xtx; C; superior temporal R; low β | B (5/2) | B (5/2) | B (5/2) | B (5/2) | B (5/2) | B (5/2) |
| Leader–Follower averaged across conditions; xtx; H; bank of superior temporal sulcus L; θ | A (5/3) | A (5/3) | A (5/3) | A (5/3) | A (4/3) | A (5/3) |
| Leader–Follower averaged across conditions; rtr; H; lateral occipital R; γ | A (4/5) | A (4/5) | A (4/5) | C (3/5) | C (3/5) | C (1/5) |
| Leader–Follower averaged across conditions; sts; H; rostral anterior cingulate R; θ | B (5/2) | B (5/2) | B (5/2) | B (5/2) | B (4/2) | C (3/3) |
| Leader–Follower averaged across conditions; yty; H; superior frontal R; low α | B (4/0) | B (4/0) | B (4/0) | B (4/0) | C (2/0) | C (2/1) |
| Leader–Follower averaged across conditions; rtr; H; supramarginal L; low β | A (4/4) | A (4/4) | A (4/4) | A (4/6) | C (3/5) | C (3/5) |
| Leader–Follower averaged across conditions; rtr; H; temporal pole R; high α | A (4/8) | A (4/8) | A (4/8) | A (4/7) | A (4/5) | C (3/5) |
| Leader–Follower averaged across conditions; rtr; C; entorhinal R; high α | A (4/6) | A (4/6) | A (4/6) | A (4/6) | C (2/5) | C (1/3) |
| Leader–Follower averaged across conditions; rtr; C; pars triangularis L; high α | A (4/4) | A (4/4) | A (4/4) | A (4/4) | A (5/4) | A (5/5) |
| **Separate response-period × condition interaction** | | | | | | |
| Interaction; xtx; H; frontal pole L; γ | B (4/1) | B (4/1) | B (4/1) | B (4/1) | B (4/1) | C (3/1) |
| Interaction; rtr; C; caudal middle frontal L; θ | B (4/0) | B (4/0) | B (4/0) | B (4/0) | C (3/0) | C (2/0) |
| Interaction; rtr; C; inferior temporal L; low α | A (4/7) | C (3/6) | A (4/6) | A (5/6) | A (4/6) | A (4/4) |

*Note.* H = homologous; C = cross-hemispheric; L = left; R = right. Each cell reports the τ-specific tier followed by the number of dyads meeting the trial-shuffle and time-shift criteria, respectively. Time-shift controls were conducted for all targets at each τ; a count of zero means tested without dyad-level recurrence. Tier A required trial-shuffle recurrence in at least four dyads and time-shift recurrence in at least three. Tier B met the trial-shuffle requirement without meeting both Tier A requirements; Tier C comprised the remaining targets. The right postcentral theta target used the common-support window −968 to −879 ms for this sweep, rather than its primary window −999 to −879 ms. Primary ΦID estimates and tiers remain as reported in Table S6a.

### Table S6d. History/τ embedding sensitivity and surrogate recurrence for ΦID targets

| Target | Primary tier | History tier | Trial-shuffle p (dyads) | Time-shift p (dyads) |
|---|---|---|---|---|
| **Four condition and response-period contrasts** | | | | |
| Matching–Mirroring; yty; cross-hemispheric; insula L; high β | A | A | .0099 (4) | .0099 (5) |
| Matching–Mirroring; rtr; cross-hemispheric; medial orbitofrontal L; low β | B | B | .0099 (5) | .0198 (0) |
| Leader–Follower within Mirroring; rtr; cross-hemispheric; isthmus cingulate R; γ | B | B | .0099 (4) | .0099 (1) |
| Leader–Follower within Matching; rtr; homologous; isthmus cingulate L; low β | B | B | .0099 (4) | .0495 (2) |
| Leader–Follower within Matching; rtr; cross-hemispheric; paracentral R; low α | A | A | .0495 (4) | .0396 (3) |
| Leader–Follower averaged across conditions; xtx; cross-hemispheric; superior temporal R; low β | B | B | .0495 (5) | .0198 (2) |
| Leader–Follower averaged across conditions; xtx; homologous; bank of superior temporal sulcus L; θ | A | A | .0495 (5) | .0396 (3) |
| Leader–Follower averaged across conditions; rtr; homologous; lateral occipital R; γ | A | A | .0099 (4) | .0099 (5) |
| Leader–Follower averaged across conditions; sts; homologous; rostral anterior cingulate R; θ | B | C | .2277 (1) | Not tested |
| Leader–Follower averaged across conditions; yty; homologous; superior frontal R; low α | B | B | .0099 (4) | .0495 (0) |
| Leader–Follower averaged across conditions; rtr; homologous; supramarginal L; low β | A | A | .0099 (4) | .0297 (4) |
| Leader–Follower averaged across conditions; rtr; homologous; temporal pole R; high α | A | A | .0099 (4) | .0099 (8) |
| Leader–Follower averaged across conditions; rtr; cross-hemispheric; entorhinal R; high α | A | A | .0099 (4) | .0099 (6) |
| Leader–Follower averaged across conditions; rtr; cross-hemispheric; pars triangularis L; high α | A | A | .0099 (4) | .0099 (4) |
| **Separate response-period × condition interaction** | | | | |
| Interaction; xtx; homologous; frontal pole L; γ | B | B | .0495 (4) | .0099 (1) |
| Interaction; rtr; cross-hemispheric; caudal middle frontal L; θ | B | B | .0198 (4) | .1089 (0) |
| Interaction; rtr; cross-hemispheric; inferior temporal L; low α | A | A | .0099 (4) | .0099 (7) |

*Note.* L = left; R = right. Values in parentheses are the numbers of dyads meeting the respective surrogate criterion; p values are group-level surrogate results. The observed-data embedding selection was H = 1 and τ = 1 for all 17 eligible targets. The homologous right rostral anterior cingulate theta sts target retained primary Tier B over −951 to −774 ms. Its history check used the shorter common-support window −808 to −774 ms and yielded Tier C. Both observed data and all trial surrogates selected H = 1, τ = 1; fixing those settings reproduced the shortened-window result. Its history-specific time-shift control was not tested because the trial-shuffle gate was not met. Two boundary targets—the homologous right postcentral theta rtr target and the homologous right rostral anterior cingulate theta sts target within Mirroring—were not eligible for the full history/τ grid. Primary estimates and tiers remain as reported in Table S6a.

### Table S6e. Signal-timescale diagnostics for primary Tier A ΦID targets

| Target | Actor | Span | ACF zero crossing | ACF first minimum | MI first minimum |
|---|---|---|---|---|---|
| Matching–Mirroring; rtr; homologous; postcentral R; θ | Leader | Feature | 45.0 [44.0–46.1] | 65.5 [64.5–65.6] | 4.5 [4.0–5.0] |
| | Follower | Feature | 44.8 [44.5–46.1] | 65.0 [64.5–65.5] | 4.5 [4.0–5.0] |
| | Leader | Full epoch | 150.2 [147.5–154.9] | 216.5 [214.8–224.9] | 56.5 [56.0–57.0] |
| | Follower | Full epoch | 149.5 [146.6–153.2] | 219.2 [215.5–224.5] | 56.5 [56.0–56.6] |
| Matching–Mirroring; yty; cross-hemispheric; insula L; high β | Leader | Feature | 53.5 [52.0–54.0] | 73.0 [71.6–74.9] | 5.0 [5.0–5.0] |
| | Follower | Feature | 52.0 [50.9–52.9] | 73.5 [71.1–74.2] | 5.0 [5.0–5.0] |
| | Leader | Full epoch | 167.8 [165.5–170.0] | 247.0 [244.5–250.6] | 55.2 [55.0–55.5] |
| | Follower | Full epoch | 167.5 [165.0–170.0] | 247.5 [243.8–254.8] | 55.0 [55.0–55.9] |
| Leader–Follower within Mirroring; sts; homologous; rostral anterior cingulate R; θ | Leader | Feature | 48.8 [48.5–50.0] | 75.5 [75.0–76.6] | 5.0 [4.5–5.0] |
| | Follower | Feature | 50.5 [47.9–52.0] | 76.8 [76.4–77.9] | 5.0 [4.5–5.0] |
| | Leader | Full epoch | 181.0 [168.1–194.6] | 241.0 [237.9–258.2] | 56.0 [55.5–56.4] |
| | Follower | Full epoch | 184.5 [174.6–201.9] | 256.0 [245.2–270.9] | 56.5 [55.8–56.8] |
| Leader–Follower within Matching; rtr; cross-hemispheric; paracentral R; low α | Leader | Feature | 46.0 [44.4–47.5] | 67.8 [66.8–69.0] | 4.0 [4.0–4.5] |
| | Follower | Feature | 47.2 [45.4–47.8] | 68.2 [67.4–69.2] | 4.5 [4.0–4.5] |
| | Leader | Full epoch | 161.0 [156.4–168.9] | 232.0 [227.6–244.6] | 54.0 [53.5–54.5] |
| | Follower | Full epoch | 165.5 [156.5–171.5] | 232.5 [225.8–238.2] | 53.5 [53.5–54.2] |
| Leader–Follower averaged across conditions; xtx; homologous; bank of superior temporal sulcus L; θ | Leader | Feature | 42.0 [40.9–43.8] | 62.2 [61.6–63.0] | 4.0 [4.0–4.0] |
| | Follower | Feature | 42.0 [40.8–43.1] | 61.2 [61.0–61.8] | 4.0 [4.0–4.0] |
| | Leader | Full epoch | 159.8 [155.0–162.4] | 230.5 [229.1–235.6] | 55.0 [54.6–55.0] |
| | Follower | Full epoch | 162.0 [157.5–165.1] | 229.2 [225.5–233.6] | 55.0 [54.5–55.5] |
| Leader–Follower averaged across conditions; rtr; homologous; lateral occipital R; γ | Leader | Feature | 51.5 [51.0–52.2] | 71.5 [70.2–73.1] | 5.0 [5.0–5.0] |
| | Follower | Feature | 52.0 [51.4–54.2] | 73.0 [72.0–74.2] | 5.0 [5.0–5.0] |
| | Leader | Full epoch | 161.2 [158.0–163.9] | 240.2 [235.2–244.1] | 55.5 [55.0–56.0] |
| | Follower | Full epoch | 160.5 [159.1–163.2] | 238.8 [234.5–243.2] | 55.5 [55.0–55.5] |

| Target | Actor | Span | ACF zero crossing | ACF first minimum | MI first minimum |
|---|---|---|---|---|---|
| Leader–Follower averaged across conditions; rtr; homologous; supramarginal L; low β | Leader | Feature | 47.5 [45.6–48.2] | 67.8 [67.5–69.0] | 5.0 [4.5–5.0] |
| | Follower | Feature | 47.2 [45.8–48.1] | 68.5 [67.8–69.0] | 5.0 [4.5–5.0] |
| | Leader | Full epoch | 161.5 [156.8–166.1] | 241.8 [239.6–245.2] | 57.0 [56.8–57.5] |
| | Follower | Full epoch | 159.8 [158.0–169.0] | 241.8 [238.1–246.1] | 57.0 [56.2–57.1] |
| Leader–Follower averaged across conditions; rtr; homologous; temporal pole R; high α | Leader | Feature | 43.5 [42.5–44.2] | 60.5 [60.0–61.0] | 4.0 [4.0–4.4] |
| | Follower | Feature | 43.0 [42.1–44.5] | 60.5 [59.8–61.0] | 4.0 [4.0–4.2] |
| | Leader | Full epoch | 162.8 [158.4–168.4] | 240.0 [235.5–247.2] | 55.5 [55.5–56.0] |
| | Follower | Full epoch | 158.8 [156.0–164.2] | 239.0 [234.5–246.6] | 55.2 [54.9–55.8] |
| Leader–Follower averaged across conditions; rtr; cross-hemispheric; entorhinal R; high α | Leader | Feature | 47.5 [46.5–49.2] | 68.0 [66.6–68.0] | 4.8 [4.2–5.0] |
| | Follower | Feature | 47.0 [45.5–49.5] | 68.5 [67.5–68.5] | 4.5 [4.0–5.0] |
| | Leader | Full epoch | 161.5 [156.5–171.0] | 237.2 [231.4–246.6] | 56.0 [55.5–56.5] |
| | Follower | Full epoch | 163.5 [159.2–169.4] | 240.0 [236.6–252.4] | 56.0 [55.9–56.2] |
| Leader–Follower averaged across conditions; rtr; cross-hemispheric; pars triangularis L; high α | Leader | Feature | 45.5 [44.1–46.9] | 64.5 [64.0–65.2] | 4.0 [4.0–4.2] |
| | Follower | Feature | 44.5 [44.0–45.1] | 65.2 [64.4–66.0] | 4.0 [4.0–4.2] |
| | Leader | Full epoch | 161.5 [157.0–169.4] | 239.2 [229.0–242.0] | 56.0 [55.5–56.2] |
| | Follower | Full epoch | 158.0 [155.0–164.2] | 236.2 [229.1–241.0] | 55.5 [55.5–56.0] |
| Separate response-period × condition interaction; rtr; cross-hemispheric; inferior temporal L; low α | Leader | Feature | 48.8 [47.0–49.0] | 73.8 [72.9–74.2] | 5.0 [5.0–5.0] |
| | Follower | Feature | 48.2 [46.8–49.6] | 74.2 [73.5–75.6] | 5.0 [5.0–5.0] |
| | Leader | Full epoch | 162.0 [157.6–164.6] | 239.8 [230.4–251.4] | 55.2 [54.2–55.5] |
| | Follower | Full epoch | 158.0 [156.4–160.4] | 239.8 [229.4–258.5] | 55.0 [54.2–55.5] |

*Note*. L = left; R = right; ACF = autocorrelation function; MI = mutual information. Values are median [first–third quartile] in milliseconds across 15 dyads. Feature windows are the saved target intervals; full epochs span −1,000 to +1,000 ms around response-period onset. Leader and Follower refer to the actor signals. The diagnostics describe signal timescales and do not alter the primary ΦID estimates or robustness tiers.

### S7.6. MMI Redundancy Choice and Interpretation of ΦID Atoms

ΦID was computed using the Gaussian/correlation-based estimator with minimum mutual information (MMI) redundancy and its double-redundancy extension (Barrett, 2015; Luppi et al., 2022; Mediano et al., 2025). MMI redundancy is a defensible operational choice for continuous Gaussian PID/ΦID analyses because it provides a tractable decomposition of redundant, unique, and synergistic information in continuous time-series data (Barrett, 2015). In two-source, single-target Gaussian PID, MMI defines redundancy as the minimum mutual information carried by either individual source about the target, which makes that redundancy estimate independent of the correlation structure between sources (Barrett, 2015).

Accordingly, ΦID atoms estimated under MMI should be interpreted as MMI-defined information geometries rather than as redundancy, uniqueness, or synergy under all possible PID redundancy functions.

**Supplementary S8. Extended Discussion Notes**

This section contains methodological detail and extended interpretation relocated from the main Discussion to maintain concision. Each sub-section is cross-referenced in the main text.

**S8.1 Tonic Neural Role-Invariance: Independent Per-Role FDR Validation**

To evaluate the spatial overlap of tonic Condition effects across roles, mixed-effects models were re-estimated independently within Leader and Follower subsets, with Benjamini–Hochberg FDR correction ($q \leq .05$) applied separately within each role. Spatial overlap was quantified using the Jaccard index on independently FDR-corrected ROI × band survivor sets. Under this procedure, spatial overlap was high but not complete. For the dominant contrast (Matching > Mirroring), overlap was near-complete (Jaccard ≈ 0.95; 63/66 ROIs), whereas the weaker contrast (Mirroring > Matching) showed more moderate overlap (Jaccard ≈ 0.57; 4/7 ROIs). Despite these differences in spatial extent, effect magnitudes were effectively identical across roles (median $|\Delta\beta| < 10^{-4}$). These results indicate that tonic condition effects are encoded within a predominantly shared neural architecture across roles, with minor role-dependent variation in the spatial distribution of weaker effects.

**S8.2 Restriction to Diagonal Persistence Atoms**

ΦID decomposes the mutual information between the dyad's joint past and future states into a 4 × 4 lattice of 16 atoms. In the present study, inference was restricted a priori to four diagonal persistence atoms: synergy-preserving information (sts), leader-unique-preserving information (xtx), follower-unique-preserving information (yty), and redundancy-preserving information (rtr). These atoms index whether the same information mode is maintained across the local past–future transition and were examined within the Leader/Follower × Mirroring/Matching task structure. The measurement space was thus hypothesis-driven, whereas the spatial, spectral, temporal, geometric, and contrast-specific localization of effects within that space was determined empirically.

The remaining twelve off-diagonal atoms describe transformations between information modes, such as redundant information becoming synergistic or role-specific information becoming redundant. Although theoretically meaningful, these atoms address a different inferential question concerning how information changes its mode of expression over time. Their inclusion would also substantially expand the analysis when crossed with regions, frequency bands, contrasts, source configurations, and surrogate procedures. Future work with hypotheses directed at information-mode transformations should extend this framework to the off-diagonal atoms.
